# Comparative lifespan and healthspan of nonhuman primate species common to biomedical research

**DOI:** 10.1101/2024.07.31.606010

**Authors:** Hillary F Huber, Hannah C Ainsworth, Ellen E Quillen, Adam Salmon, Corinna Ross, Adinda D Azhar, Karen Bales, Michele A Basso, Kristine Coleman, Ricki Colman, Huda S Darusman, William Hopkins, Charlotte E Hotchkiss, Matthew J Jorgensen, Kylie Kavanagh, Cun Li, Julie A Mattison, Peter W Nathanielsz, Suryo Saputro, Diana G Scorpio, Paul-Michael Sosa, Eric J Vallender, Yaomin Wang, Caroline J Zeiss, Carol A Shively, Laura A Cox

## Abstract

There is a critical need to generate age- and sex-specific survival curves to characterize chronological aging consistently across nonhuman primates (NHP) used in biomedical research. Sex-specific Kaplan-Meier survival curves were computed in 12 translational aging models: baboon, bonnet macaque, chimpanzee, common marmoset, coppery titi monkey, cotton-top tamarin, cynomolgus macaque, Japanese macaque, pigtail macaque, rhesus macaque, squirrel monkey, and vervet/African green. After employing strict inclusion criteria, primary results are based on 12,269 NHP that survived to adulthood and died of natural/health-related causes. A secondary analysis was completed for 32,616 NHP that died of any cause. Results show a pattern of reduced male survival among catarrhines (African and Asian primates), especially macaques, but not platyrrhines (Central and South American primates). For many species, median lifespans were lower than previously reported. An important consideration is that these analyses may offer a better reflection of healthspan than lifespan since research NHP are typically euthanized for humane welfare reasons before their natural end of life. This resource represents the most comprehensive characterization of sex-specific lifespan and age-at-death distributions for 12 biomedically relevant species, to date. These results clarify relationships among NHP ages and provide a valuable resource for the aging research community, improving human-NHP age equivalencies, informing investigators of expected survival rates, providing a metric for comparisons in future studies, and contributing to understanding of factors driving lifespan differences within and among species.

## Introduction

Nonhuman primates (NHPs) are genetically, physiologically, and behaviorally the best translational models for human aging as their genomes, developmental trajectory, reproductive strategies, and aging-related changes in physical function, cognitive function, and disease development are more similar to humans than those of other mammals.^1–4^ Yet, there is limited information regarding longevity in the NHPs most commonly used as translational models. Few studies have attempted cross-species comparisons and reports are often contradictory, likely due to the use of different methodological approaches (e.g., inclusion criteria). To determine how NHP ages correspond with human age, it is essential to fully characterize the demography of NHP longevity within each species, rather than focusing on individual reports of maximum longevity. Numerous publications list NHP maximum lifespans in tables that include a variety of other life history features, but few cite primary sources. This leads to overreporting of the same statistics without verifying the validity of the measure or the relevance to animals under study. For example, 37.5 years is often cited as the lifespan of baboons (*Papio hamadryas* spp.).^5–8^ However, tracing citations to the primary source reveals that this statistic comes from a single baboon that died at the Brookfield Zoo in 1972; the birth date is given as June 1, 1935 (one year after the zoo opened), but it is not documented whether this date is known or estimated.^9^ This estimate of maximum longevity in baboons is not particularly useful without additional context such as the number of baboons surviving to the maximum or knowledge of the median baboon lifespan. Median captive baboon lifespan has been reported as 21^10^ or 11^11^ years but the report of maximum longevity is more frequently cited. It is likely that the discrepancy in median baboon lifespan reflects differences in methodological approaches to data analysis. This example in baboons highlights how differences in analytic approaches across studies make it difficult to compare reports within or across species. The unclear and limited data on NHP lifespan, such as the reporting of maximum longevity to indicate “lifespan,” creates confusion in scientific analysis and in the peer review process.

Cross-species comparisons are a major goal of aging research since they can reveal factors contributing to variation in lifespans. Inconsistent lifespan estimates are problematic when looking at a single species, and the problem is compounded by cross-species comparisons. We address this knowledge gap by creating rigorous and reproducible survivorship data, identifying mortality risk and its relationship to biological age at different chronological ages, and examining the shape of mortality and healthspan curves across 12 captive NHP species. The initial dataset, prior to quality control and filtering, included lifespan data from 114,255 animals from 58 species at 15 institutions. We highlight that while maximum age is an easily reported statistic as it is purely observational, calculating median lifespan is more challenging, as methodological decisions about inclusion and exclusion criteria vary among studies, producing substantial discrepancies across cohorts and species. With the data herein, we have the unique ability to calculate survival probabilities using the same criteria for all 12 species, producing the most methodologically consistent cross-species comparison to date. The value of such a large dataset is the ability to filter the data to the most representative sample and retain adequate sample sizes for statistical analyses. In this study, survival curves were generated on animals that survived to at least adulthood (defined in Methods) because, as in most mammals including humans, risk of death in infancy is substantial and strongly biases the median lifespan. Primary results and comparisons by sex are built using data from animals that died of natural causes or were euthanized for clinical/health reasons. This report provides comprehensive data summaries and tools to improve biomedical research involving NHPs within and beyond the field of aging.

## Methods

### Species

Twelve NHP species for analyses are shown in **Table 1**. We are considering all members of the genus *Papio* a single species and considering Indian- and Chinese-origin rhesus macaques together, as captive research baboons have a high degree of morphotype mixing^12,13^ and captive rhesus are similarly highly admixed from these geographic source populations.^14^ We included chimpanzees (*Pan troglodytes* spp.), but it must be noted that biomedical research with great apes is heavily restricted across the world. Still, many retired chimpanzees reside at research facilities and they provide a valuable comparison since their estimated lifespan is between that of humans and the monkey species commonly found at biomedical research facilities. Similarly, while cotton-top tamarins (*Saguinus oedipus*) were at one time biomedical research models, they have not been used for that purpose since 2008 when deforestation resulted in animals being listed as critically endangered.

**Table 1.**
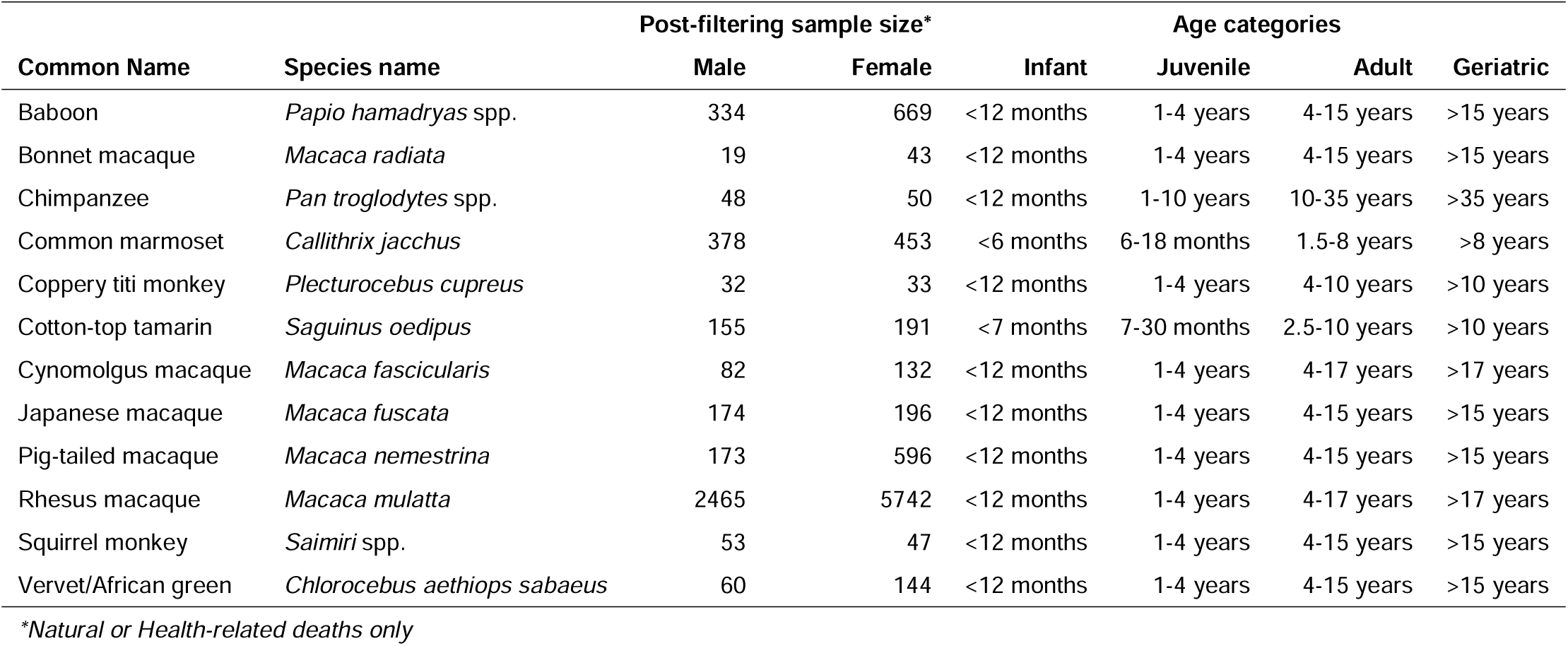
Sample sizes of primary analysis datasets and species-specific age categories. For each species, age categories and estimated age ranges are shown. ^33,44,45^

| Common Name | Species name | Post-filtering sample size* |  | Age categories |  |  |  |
| --- | --- | --- | --- | --- | --- | --- | --- |
|  |  | Male | Female | Infant | Juvenile | Adult | Geriatric |
| Baboon | *Papio hamadryas* spp. | 334 | 669 | <12 months | 1-4 years | 4-15 years | >15 years |
| Bonnet macaque | *Macaca radiata* | 19 | 43 | <12 months | 1-4 years | 4-15 years | >15 years |
| Chimpanzee | *Pan troglodytes* spp. | 48 | 50 | <12 months | 1-10 years | 10-35 years | >35 years |
| Common marmoset | *Callithrix jacchus* | 378 | 453 | <6 months | 6-18 months | 1.5-8 years | >8 years |
| Coppery titi monkey | *Plecturocebus cupreus* | 32 | 33 | <12 months | 1-4 years | 4-10 years | >10 years |
| Cotton-top tamarin | *Saguinus oedipus* | 155 | 191 | <7 months | 7-30 months | 2.5-10 years | >10 years |
| Cynomolgus macaque | *Macaca fascicularis* | 82 | 132 | <12 months | 1-4 years | 4-17 years | >17 years |
| Japanese macaque | *Macaca fuscata* | 174 | 196 | <12 months | 1-4 years | 4-15 years | >15 years |
| Pig-tailed macaque | *Macaca nemestrina* | 173 | 596 | <12 months | 1-4 years | 4-15 years | >15 years |
| Rhesus macaque | *Macaca mulatta* | 2465 | 5742 | <12 months | 1-4 years | 4-17 years | >17 years |
| Squirrel monkey | *Saimiri* spp. | 53 | 47 | <12 months | 1-4 years | 4-15 years | >15 years |
| Vervet/African green | *Chlorocebus aethiops sabaeus* | 60 | 144 | <12 months | 1-4 years | 4-15 years | >15 years |

\*Natural or Health-related deaths only

### Participating institutions

Data from eight United States National Primate Research Centers (NPRCs) are included: California (CNPRC), Emory (ENPRC), New England (NEPRC; this center is no longer open but we obtained archival data), Oregon (ONPRC), Southwest (SNPRC), Tulane (TNPRC), Washington (WaNPRC), and Wisconsin (WNPRC). Data also originated from Primate Research Center IPB University in Indonesia, Keeling Center for Comparative Medicine and Research at The University of Texas MD Anderson Cancer Center, National Institute on Aging Intramural Research Program, Sam and Ann Barshop Institute for Longevity and Aging Studies at UT Health San Antonio, Vervet Research Colony at Wake Forest University, and Yale University. **Supplementary Table S1** shows species sample sizes contributed by each institute. A data extraction standard operating protocol (SOP) was developed to ensure consistency among institutions. The SOP requested data from all NHPs that were born and died at the same institute going back through all historical records, along with sex, species, date of birth, date of death, and disposition (i.e., death) code and description. We received data from 27 species categories at the Duke Lemur Center, but ultimately did not include these data herein because they did not meet stage 1 filtering requirements of this study. We also note that life history profiles for these animals are published^15^ and the data are available for public download (https://lemur.duke.edu/duke-lemur-center-database/).

### Data Filtering and Quality Control

Received data were first processed via a series of quality control checks for non-NHP species labels, inconsistent or undefined codes, and duplicated records (e.g., ensuring one observation (date of birth and death) per animal in data). We attempted to resolve inconsistencies or undefined codes via follow-up with the original data source. Records that were unable to be resolved were removed from subsequent analyses. The resulting data were then parsed through a two-stage filtering process. Stage One filtering retained records with: 1) sex classified as male or female, 2) known date of birth (not estimated), and 3) survived at least 30 days (removing neonatal deaths). Species were then filtered to only include those which retained at least 150 animals. These Stage One filtered data yielded over 77,000 animals across 12 species. Stage Two filtering retained 1) animals that survived to adulthood using the National Institutes of Health Nonhuman Primate Evaluation and Analysis table of NHP life stages (**Table 1**).^16^ The earliest age listed as adult for each species was used, supplemented by additional references for two species not present in the table, chimpanzees^17^ and coppery titi monkeys.^18^ Stage Two filtering also implemented a date of birth (DOB) cutoff. This step was critical for survival analyses and lifespan inference as received data did not include records on alive animals. Removing later (more recent) births avoided skewing results towards earlier deaths, and inference was thus based on the dataset of animals that had greatest opportunity to live to their maximum ages (**Supplementary Figure S1**). The DOB threshold was implemented by retaining animals born before 2023 minus the number of years corresponding to the initial assessment of the 85th percentile of lifespan for that species (combined sexes; non-natural deaths as censored events). In total, this filtering stage yielded a dataset of 32,616 animals, across 12 species.

#### Defining censored events by death types

Given that these data did not include alive animals, for survival analyses, censored events were based on death type, as follows: 1) death types pertaining to research sacrifice and colony management were categorized as right censored events; 2) death types pertaining to natural causes or humane euthanasia for health reasons were coded as un-censored events. Right censoring is a statistical approach in survival analysis that enables inclusion of the knowledge that the subject survived at least to that point.^19^ Treating deaths related to research sacrifice and colony management as right-censored events enabled animals to contribute to the survivorship model up until age of censoring. That is, this accounts for the lack of knowledge of how long the animal would have lived until a natural or health-related death. The final Stage Two filtered dataset was comprised of 12,269 events and 20,347 censored events.

### Statistical analyses

We computed the Kaplan-Meier estimator^20^ of the survivorship function for each species and sex, using the ggsurvfit package^21^ in R version 4.1.2. Survival curves and median lifespan estimates were calculated for both including and excluding censored (research sacrifice; colony management death types) data. A critical analytic consideration was that censoring was greatly biased by sex. Thus, the primary analyses presented with comparisons by sex were limited to natural/health-related deaths only (no censored data). For many species, proportional hazards assumptions were violated (preventing usage of the cox-proportional hazards model), but since the primary analysis datasets were absent of censored events, analyses were not restricted to methods for censored data. The analysis plan followed one that was applicable across all twelve species of various sample sizes. For each species, maximum ages were compared between males and females using two analytic approaches. First, quantile regression models were analyzed in SAS version 9.2 using the QUANTREG procedure at the 25^th^, 50^th^, 75^th^, and 85^th^ maximum age percentiles with sex as the predictor and primate center was included as a covariate. Effects of sex at each percentile were tested using the Wald statistic and standard errors for regression coefficients were computed using resampling method (seed=12333). For each species, we also tested for differences in the maximum age distributions by sex using the nonparametric two-sample Kolmogorov-Smirnov test (ks.test function in R version 4.1.2), two-sided test p-values are reported.^20^ Finally, to evaluate the uniformity of the rate of decline across survivorship curves, we fit an exponential model (e^β^), separately, to the first and last quartiles of the Kaplan-Meier survival curves using the nonlinear least squares function in R (version 4.1.2), shown in **Supplementary Figure S2**. As β captures the function’s rate of decay, we illustrated trends across species, by sex, by plotting the magnitude of β for these two quartiles. Computations were performed using the Wake Forest University (WFU) High Performance Computing Facility.^22^

## Results

### Primary analyses

Sample counts of primary analysis datasets, featuring natural or health-related deaths only, are shown in **Table 1**. Maximum observed age including all types of deaths (e.g., research-related sacrifice, clinical/health-related euthanasia, and natural), as well as median age at death calculated from only natural and clinical deaths, are summarized by sex and species in **Table 2**. **Figure 1** shows the distribution of natural and clinical deaths, with medians, interquartile ranges, and proportions of data by sex and species. Combined survival curves for all 12 species in males and females are shown in **Figure 2**. To evaluate the rate of decline for the survivorship curves, across species, data from the first and last quartiles of the Kaplan-Meier survivorship function were fit to an exponential model that captures rate of decay (i.e., change in probability of death) (**Supplementary Figure S2**), and species were then compared within and between sexes. Comparing first and last quartiles illustrated that species predominantly experienced faster rates of death within the first quartile of adulthood. Comparing male and female rates of decline within both quartiles highlighted the faster rates of decline for males within the first quartile. However, in the last quartile, this pattern was nearly reversed; the majority of species (except cotton-top tamarin, vervet/African green monkey, and common marmoset) exhibited slower rates of decline in males compared to females (**Figure 3**).

**Figure 1.**
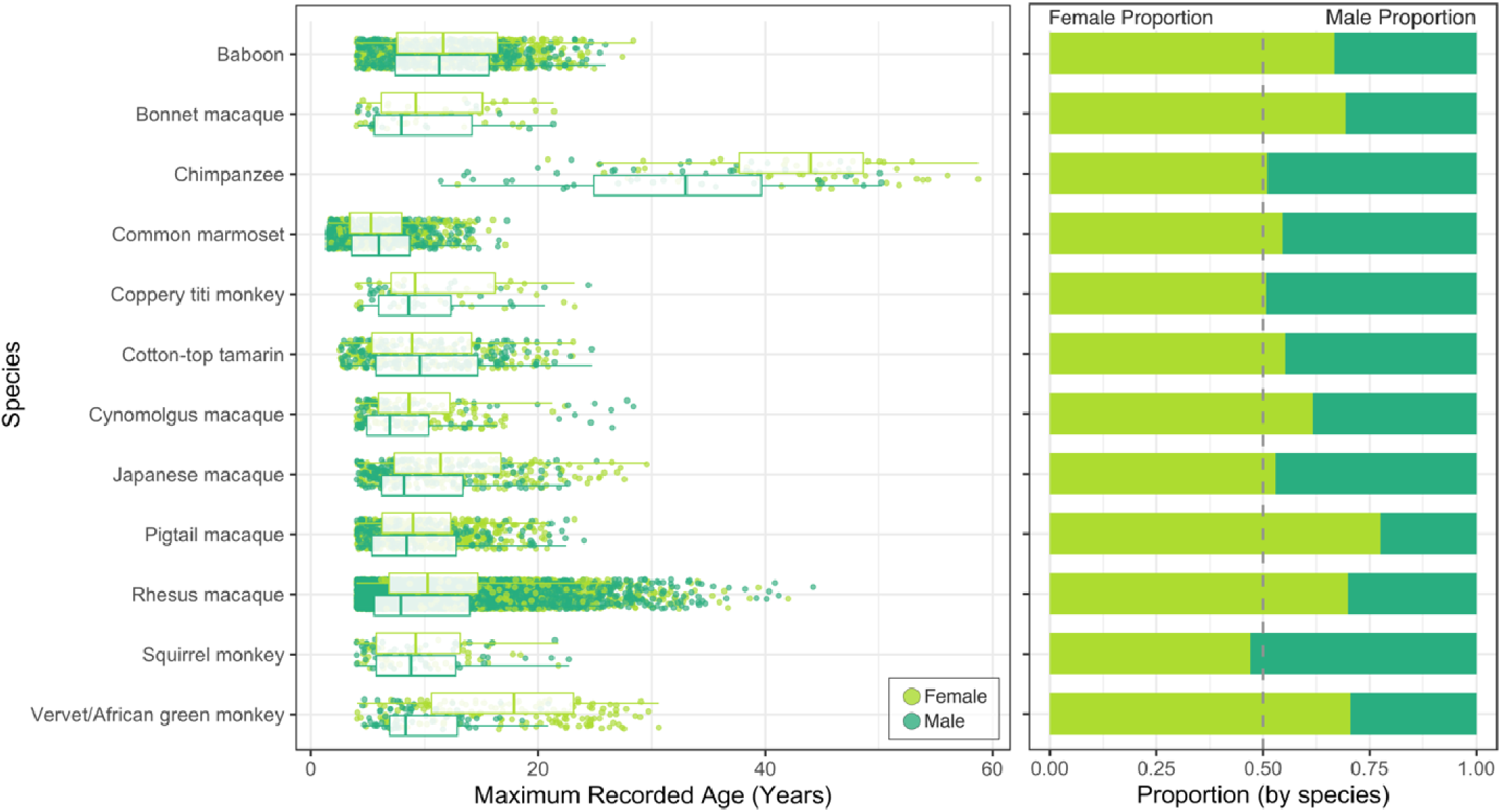
Distribution of natural and health-related euthanasia deaths by species. Boxplot overlay depicts median and interquartile range by species and sex. Proportion of data by sex and species also shown. The vertical dashed line denotes equal counts of males and females by species.

**Figure 2.**
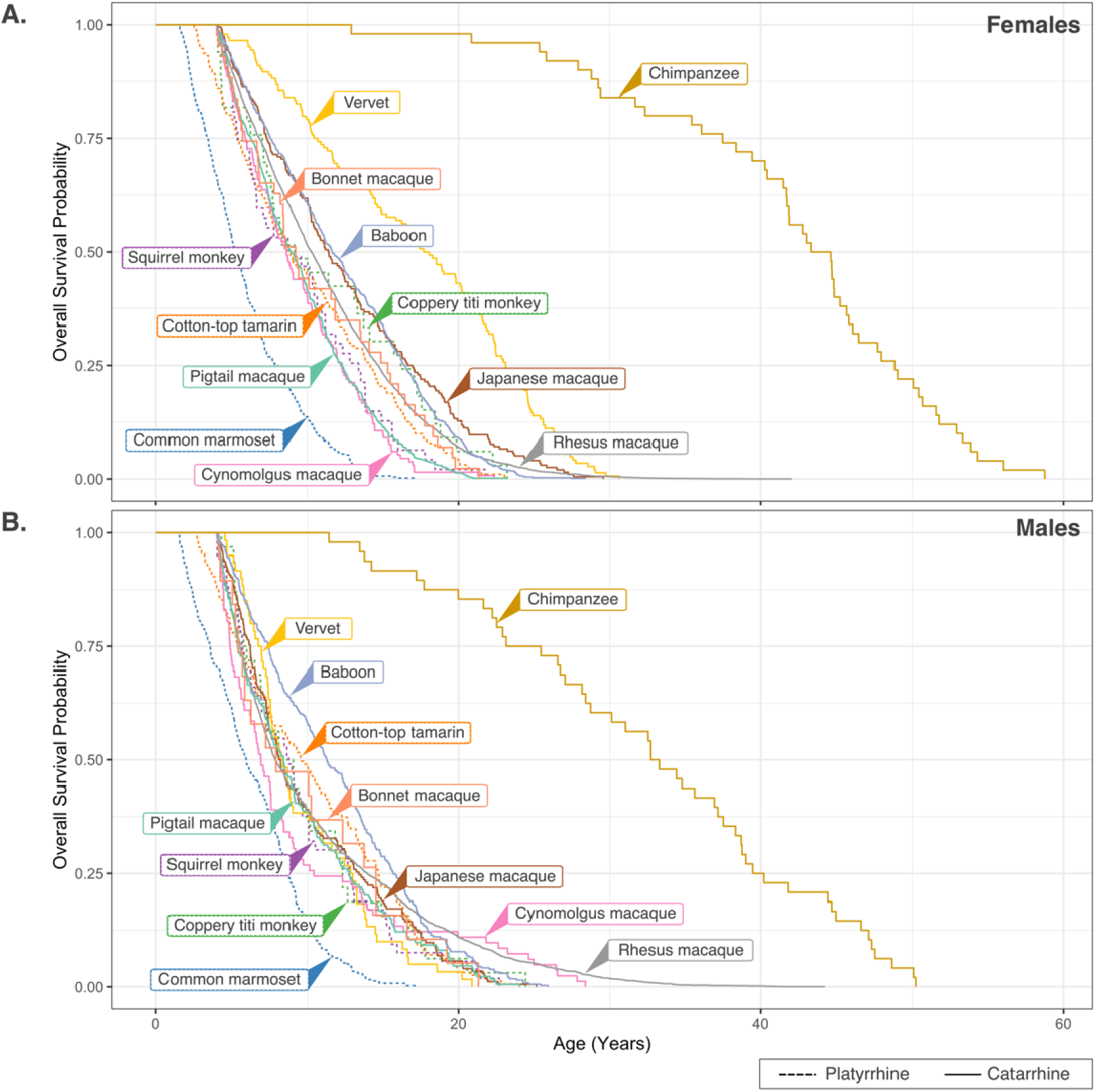
Survival curves for females (A) and males (B) of all 12 species. Data shown are for animals with deaths resulting from natural causes or humane euthanasia for health-related reasons.

**Figure 3.**
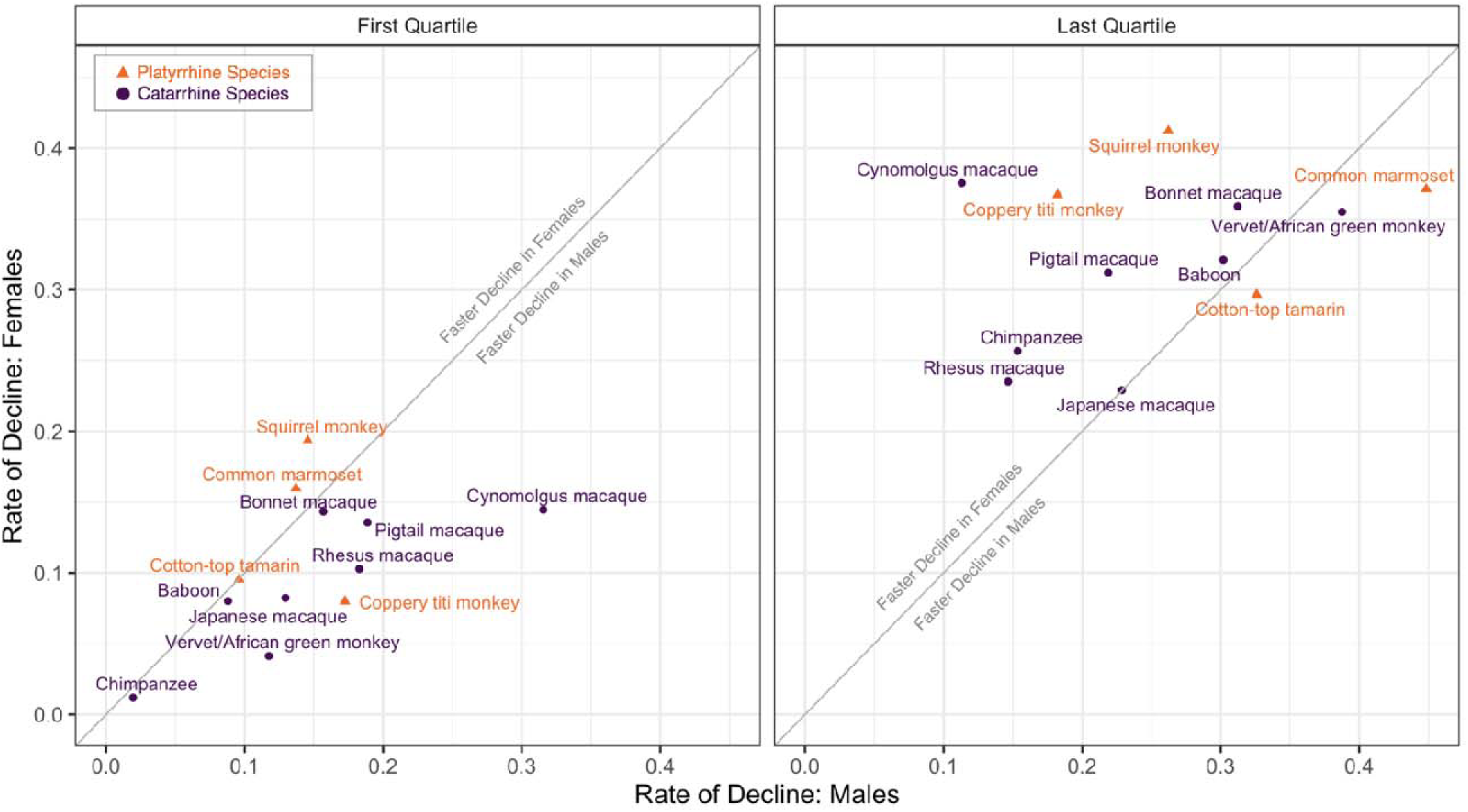
Comparison of rate of survivorship decline by quartile and sex. Rates of decline were calculated from fitting an exponential model to the first and last quartiles of the sex-specific Kaplan-Meier survival curves. Males and females are compared by quartile. Rate of decline was generally faster in males within the first quartile with the pattern nearly reversed by sex in the last quartile.

**Table 2.** Maximum and median age at death by sex and species.

| Common Name | Species name | Maximum observed age in years* |  | Median age at death in years (range)* |  |
| --- | --- | --- | --- | --- | --- |
|  |  | Male | Female | Male | Female |
| Baboon | *P. hamadryas* spp. | 30.3 | 30.6 | 11.29(10.41-12.47) | 11.65(11.08-12.44) |
| Bonnet macaque | *M. radiata* | 32.8 | 21.4 | 7.93(5.70-14.54) | 9.22(7.81-13.49) |
| Chimpanzee | *P. troglodytes* spp. | 53.3 | 58.8 | 33.00(28.41-38.33) | 43.96(41.66-45.82) |
| Common marmoset | *C. jacchus* | 17.3 | 17.1 | 5.97(5.41-6.74) | 5.31(4.92-5.66) |
| Coppery titi monkey | *P. cupreus* | 24.4 | 23.2 | 8.59(6.92-12.13) | 9.16(7.35-14.13) |
| Cotton-top tamarin | *S. oedipus* | 24.7 | 23.1 | 9.60(7.87-11.27) | 8.87(7.67-10.57) |
| Cynomolgus macaque | *M. fascicularis* | 28.4 | 23.5 | 6.93(6.21-8.18) | 8.62(7.72-9.84) |
| Japanese macaque | *M. fuscata* | 38.4 | 30.1 | 8.19(7.48-9.36) | 11.41(10.27-12.70) |
| Pig-tailed macaque | *M. nemestrina* | 27.9 | 29.2 | 8.43(7.49-9.12) | 8.96(8.43-9.59) |
| Rhesus macaque | *M. mulatta* | 44.2 | 42 | 7.89(7.65-8.24) | 10.26(10.03-10.49) |
| Squirrel monkey | *Saimiri* spp. | 22.7 | 21.8 | 8.78(6.97-10.09) | 9.22(6.55-11.19) |
| Vervet/African green | *C. aethiops sabaeus* | 24.1 | 30.6 | 8.34(7.57-10.71) | 17.87(15.24-20.23) |

\*Median age at death is calculated from natural and clinical deaths only; maximum observed age includes animals with any type of death. Maximum ages are from the current dataset only; there are known older animals of some of these species at research institutes, such as a 29-year-old titi monkey male at CNPRC and two 19-year-old male marmosets at SNPRC but these did not meet this study's filtering criteria (see methods).

For each species, individual survival curves are shown in **Figure 4** and species-specific, sex-based comparisons in **Table 3**. In most species, males showed reduced survival compared to females. Among vervets, Japanese macaques, and chimpanzees, males showed reduced survival at every age with a different overall distribution of age at death. *C*ynomolgus macaque and baboon males showed reduced survival compared to females at younger ages (25^th^ and 50^th^ percentiles), but there was no difference in survival at later stages of life. Rhesus macaque males showed reduced survival compared to females at the 25^th^, 50^th^, and 75^th^ percentiles, but females had lower age of survival at the 85^th^ percentile. There was a strong difference in the distribution of age at death between males and females (P-value=2.20x10^-16^). Pig-tailed macaque males showed reduced survival compared to females early in life (25%) but the sexes were similar at other ages. In contrast, females showed reduced survival compared to males at every age in common marmosets. Male and female survival was similar at every age with no difference in the distribution of age at death between sexes for cotton-top tamarins and squirrel monkeys. There was also no difference in distributions for coppery titi monkeys and bonnet macaques; however, the modest sample size for the species limits power to detect small differences.

**Figure 4.**
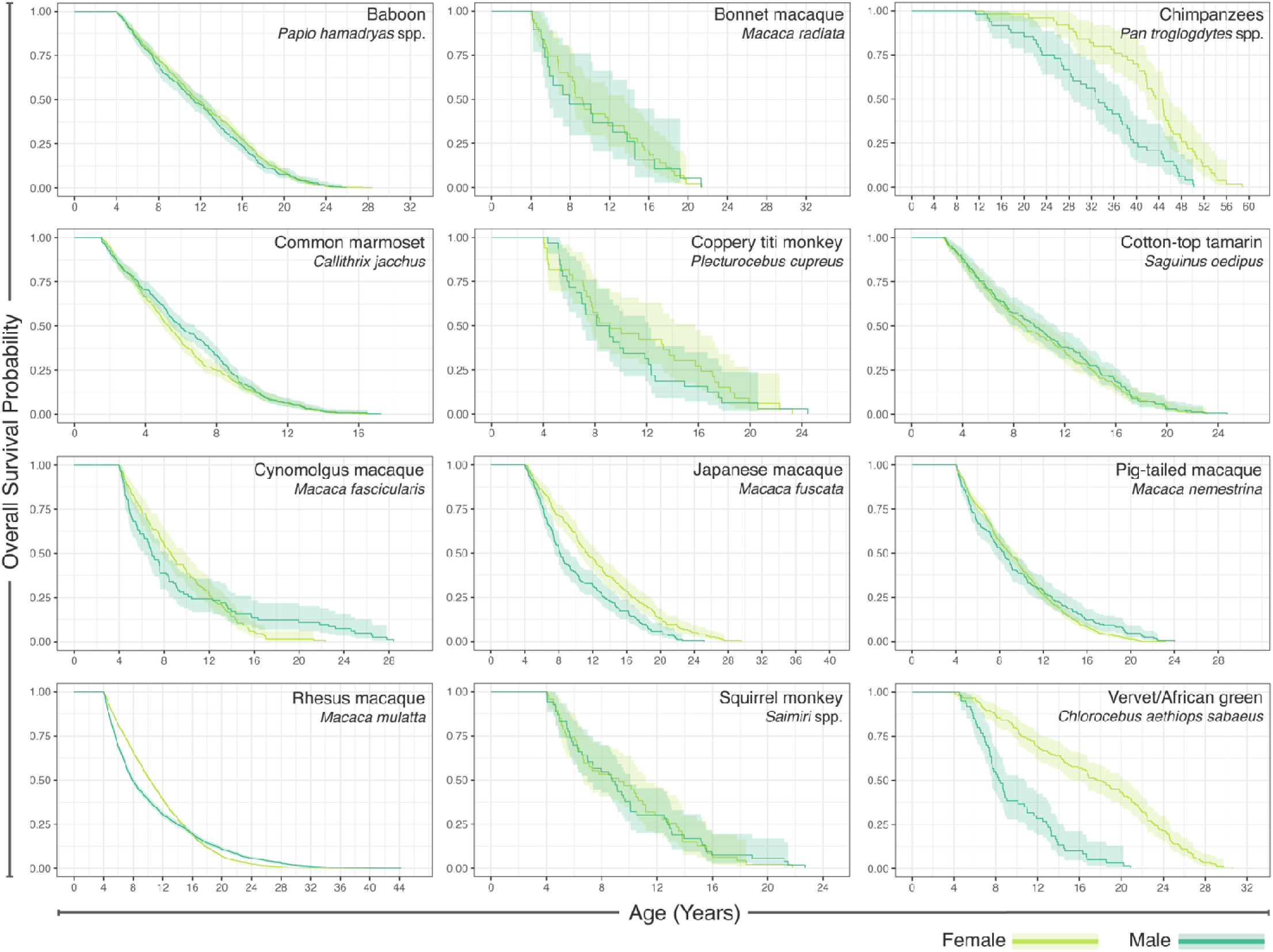
Kaplan-Meier survival curves by sex and species for natural deaths or humane euthanasia for health-related reasons. For each plot, the X-axis scaling (maximum age) is species-specific.

**Table 3.**
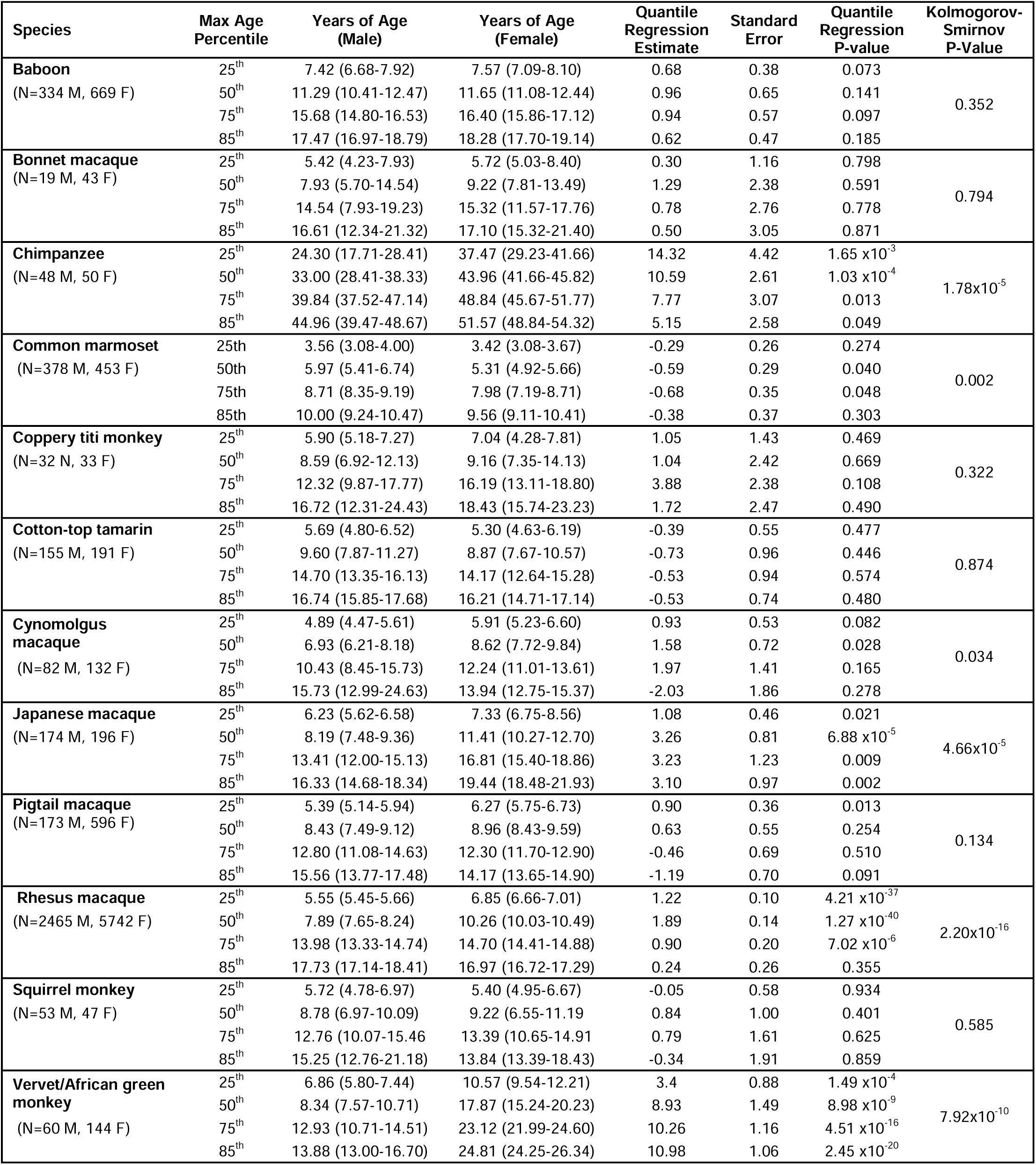
Sex-based comparisons of age by species. Quantile regression for 25^th^, 50^th^, 75^th^, and 85^th^ percentiles. Regression models adjusted for primate location (data source). Distribution of ages by sex were assessed using the Kolmogorov Smirnov test. Complete data used for analyses (natural or clinical deaths) with no censoring.

### Secondary analyses

*C*ensored data (deaths due to research sacrifice and colony management) were biased by sex (**Supplemental Figure S3**) and prevented statistical comparisons between males and females when including censored data.^19^ However, as a secondary analysis, survival curves that include censored events are presented for reference. **Supplemental Figure S4** features survival curves for each species separately with and without censored events adjacent to each other with additional details. Across species, inclusion of additional datapoints from censored events increased median lifespan estimates. We note that the high proportion of censored events (**Supplemental Figure S3**), especially in some species (i.e., greater than 50% of deaths in baboons, cynomolgus, pigtails, rhesus, squirrel monkeys, and vervets), yielded survivorship functions that never reach zero, limiting utility and inference for the full lifespan.

## Discussion

### Lifespan vs healthspan

A major consideration of note for this study is that few research NHPs live until natural death. Most are humanely euthanized due to study protocols or clinical determinations based on quality of life. The issues considered by veterinarians in making euthanasia decisions vary by facility and study protocol, but a common approach is to euthanize at the first diagnosis of major disease or injury requiring long-term treatment with reduced quality of life. Reasons for humane euthanasia may include such diverse conditions as advanced spinal or knee osteoarthritis, endometriosis, broken limbs, tumors, and meningitis – not all of which are the result of aging-related diseases. Therefore, we posit that these findings may be measuring healthspan rather than lifespan in NHP cohorts housed at research facilities. For our survival analyses, this potential limitation is partially mediated by our very large database, which enabled analyses even after removing experimental and other non-clinical deaths.

Supporting the idea that we are measuring healthspan rather than lifespan, for several species, typical age at onset of chronic disease is similar to the median lifespan estimates. Among baboons, age-related diseases are apparent around 9 years old (e.g., edema, kyphosis, prolapse, myocarditis), and by 12 years many more are evident (e.g., pancreatitis, stricture, lymphosarcoma).^23^ Median baboon lifespan in this report is 10.1 years for males and 11.1 years for females. Marmoset age-related diseases tend to emerge in animals >6 years old, including cardiovascular disease, diabetes, and neoplasias.^24^ Median marmoset lifespan in our study is 5.5 years in males and 5.0 years in females. Rhesus macaques are on average diagnosed with the first chronic condition at age 9.0 years and the second at age 10.7 years.^25^ Median rhesus lifespan in our study is 9.1 years in males and 10.6 years in females. Differences in veterinary care for these conditions mean that some pathologies in some species may be treated medically, whereas others proceed to veterinarian-suggested euthanasia. We speculate that zoo NHPs may be treated for more chronic conditions than research NHPs and would make a useful lifespan and healthspan comparison to humans.

The ability to make more accurate comparisons between NHP age and the human equivalent was a primary goal of the current analyses. Since the NHP estimates herein may be closer to healthspan than lifespan, it is useful to consider them in relation to human healthspan. The most frequently studied measures of human healthspan are deficit accumulation indices, which measure accumulation of health deficits and decline in physical function or frailty.^26–30^ In one study of 66,589 Canadians in the National Population Health Survey, accumulation of health deficits was gradual before age 46 years, with 40% of 45-50 year-olds having a frailty index score of 0 (no health deficits); starting at age 46, deficit accumulation was much more rapid, and at age 80, only 5% still had a score of 0.^30,31^ Among 73,396 people from the Longitudinal Ageing Study in India, average age of onset of any chronic disease was 53 years.^32^ We speculate that our NHP median lifespan estimates may align better with human onset and accumulation of health deficits, rather than human lifespan. However, our analysis does not address onset of health deficits, and we are unable to distinguish between which NHPs died at the end of their lifespan versus those which died at the end of their healthspan. Therefore, we are unable to make specific comparisons between human and NHP healthspans.

### Sources of variation within and between species

Our findings show great variation in adult life expectancy among all 12 species, in contrast to a prior cross-species analysis of six primate species that found little variation in adult survival.^33^ Many factors contribute to variation in adult survival. Some may assume that in captive research populations, quality of veterinary care is a major driving force. While this may have been important in the early years of NHP research, most species have been in captivity for decades and quality care is well defined. Institutional management practices are important factors, such as how decisions are made about euthanizing animals due to illness or reproductive capacity. Housing conditions are a likely influence on lifespan, as it is well known that individual versus paired versus group housing can have profound effects on health.^34–39^ The goals of the research are also important to consider. For example, rhesus monkeys have been the subjects in two longevity studies in which survival time was an outcome variable. Here, additional measures were taken to maintain older animals, which explains the extreme maximum age of rhesus macaques – 44.2 years – relative to other the other four macaque species, which show maximum ages in the 20s and 30s.^40,41^ Another potential source of bias is the way animals are selected for studies. NHPs go through health checks beforehand, and healthy animals may be preferentially selected. In our study, many of the longest-lived animals were excluded from lifespan calculations because their endpoints were research-related (**Supplementary Figure S3**). Thus, limiting the analyses to natural deaths seems to influence lifespan calculations towards younger ages.

Within species, life history features can influence lifespan. It has been proposed that reproductive strategies play an evolutionary role in regulating lifespan, since there may be tradeoffs between female fertility, investment in offspring, and longevity,^42^ although this long-held view has been challenged since the relationships between reproduction and longevity are not consistent across species.^43,44^ Adult body size also factors into survival because a longer period of growth will likely result in later reproductive maturity and a greater need for investment in offspring. In our data, common marmosets have the shortest maximum and median lifespan of all 12 species. Marmosets are also the smallest species (average weight 350-400 g), reach adulthood at the youngest age (1.5 years), and usually give birth to twins.^24,45^ However, cotton-top tamarins, the other small (average weight in captivity 565.7 g), quickly maturing (2.5 years at adulthood), twinning callitrichine^46^ in this study, has maximum and median lifespan resembling that of several larger bodied, slower maturing species that give birth to singletons, including squirrel monkeys, baboons, vervets, and macaques. It is unclear to what extent these patterns are driven by inherent species characteristics versus institutional practices, but it would be advantageous to explore this question in future studies.

Identifying physiological changes underlying the aging process and variation in lifespan and healthspan has been a major goal of the aging research community, leading to the concept of the hallmarks of aging. Nine hallmarks are now well established: genomic instability, telomere attrition, epigenetic alterations, loss of proteostasis, deregulated nutrient-sensing, mitochondrial dysfunction, cellular senescence, stem cell exhaustion, and altered intercellular communication.^47^ Five new hallmarks have been recently proposed: autophagy, microbiome disturbance, altered mechanical properties, splicing dysregulation, and inflammation.^48^ These hallmarks are thought to be molecular, cellular, and organismal level drivers of the aging process. Investigators have generated hypotheses about how the hallmarks of aging may influence lifespan within and between primate species. For example, oxidative stress is a trigger of cellular senescence and genomic instability.^49^ In a comparative analysis of 13 primate species with divergent body sizes and longevity, investigators studied reactive oxygen species production and oxidative stress resistance in cultured fibroblasts, finding some support for their hypothesis of a causal relationship with species longevity.^50^ Within species, investigators are also exploring how variation in the hallmarks contributes to individual lifespan differences. Telomere shortening has long been recognized as a marker of aging. Studies of calorie restriction in rhesus macaques have shown extension of lifespan, and investigators tested whether lifespan differences between groups could be explained by telomere length in several tissues, but interestingly, telomere length was associated with both age and sex, but not calorie restriction.^51^ The hallmarks of aging provide a productive foundation for guiding studies of the causal factors underlying lifespan variation.

### Sex-based differences

Among primates, males have been shown to have higher age-specific mortality than females throughout adulthood.^52^ We see this in some species included in the current study. One pattern is shorter lifespan among macaque males. Five macaque species (*Macaca* spp.) are reported here. In three species males have shorter median lifespan than females (cynomolgus, Japanese, and rhesus macaques). In pigtails, males have lower survival probability in early adulthood (25%) but similar survival probability at older ages, and in bonnet macaques male lifespan appears shorter in the curves and estimates, but sample size may be too small to detect a difference (female n=43, male n=19). This pattern seems to extend to all of the parvorder Catarrhini (Old World monkeys-Cercopithecoidea and apes-Hominoidea). Vervets have the largest sex-based differential with median age of 8.3 years for males and 17.9 years for females. For baboons, males show borderline lower survival probability at the 25^th^ and 75^th^ percentiles. Male chimpanzees also have lower survival probability relative to females at every life stage.

In contrast, in the parvorder Platyrrhini (Central and South American monkeys), there is generally no difference between males and females in survival estimates. For context, a phylogenetic tree for the 12 species in this study is shown in **Figure 5**.^53^ The exception is the common marmoset, with lower female survival at every age, replicating the findings of another marmoset report.^24^ The relatively short female marmoset lifespan is related to their high fertility rates.^42,45^ There are no differences in survival between males and females in coppery titi monkeys, squirrel monkeys, or cotton-top tamarins. A prior primate lifespan comparison that suggested female primates have longer lifespan than males included several catarrhine species but few data from platyrrhine species.^52^A recent study of coppery titi monkey lifespan showed a trend toward longer lifespan in males relative to females using the same population of monkeys in the current study but with different inclusion criteria.^18^

**Figure 5.**
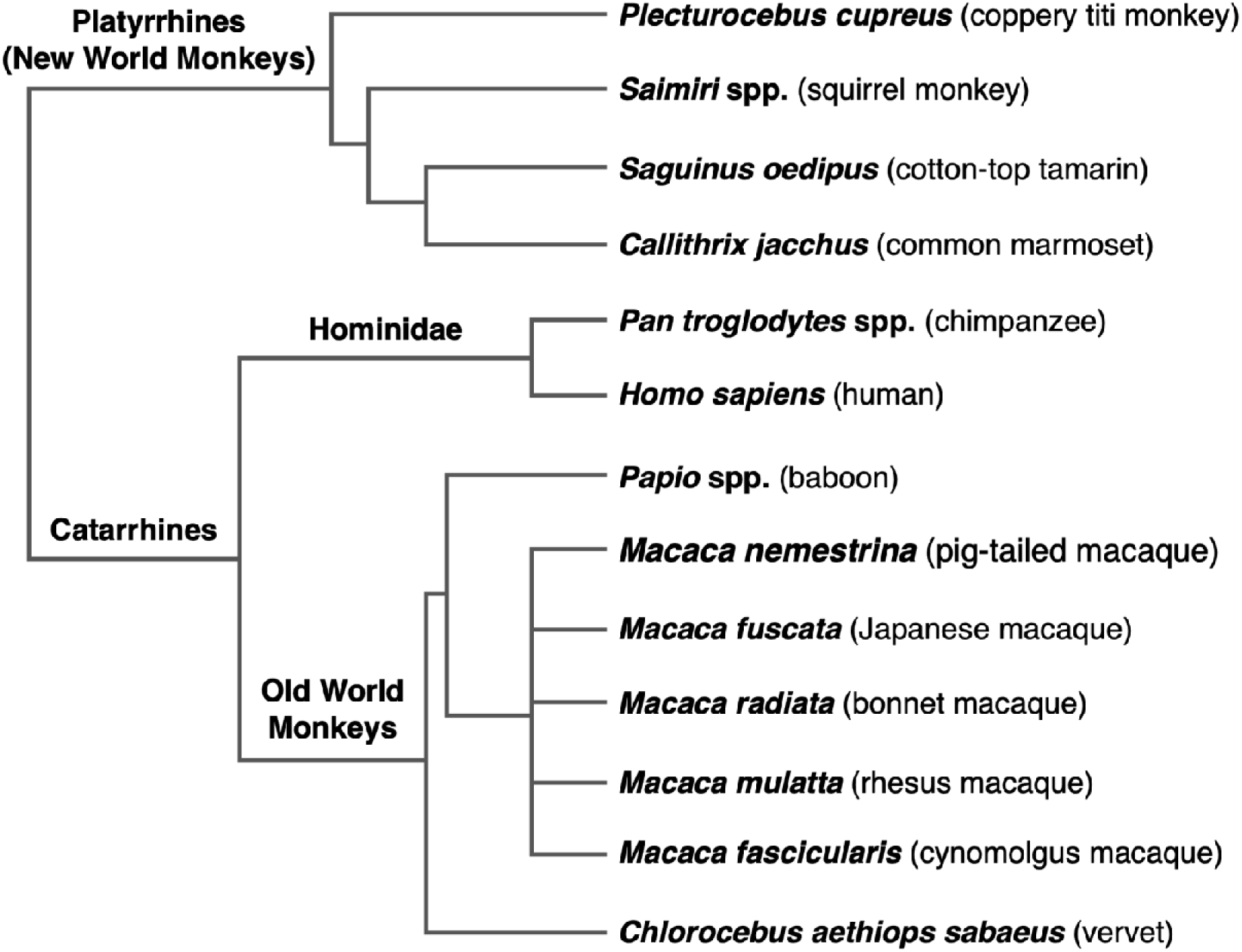
Phylogenetic tree of 12 species analyzed in study. This tree was generated with the 10kTrees Project and modified to match taxonomic names with those used in our study and to simplify the presentation.^42^ Only the 12 species studied herein are represented in the tree; there are many other sp primates in these clades not pictured.

It is difficult to know if the observed sex-based differences between catarrhine versus platyrrhine species are due to inherent species characteristics, institutional practices, or their interactions. For example, in catarrhine monkeys, it is common to house a single breeding or vasectomized male with multiple females. Fewer males than females are needed for breeding programs because males will mate with multiple females. In some species, especially baboons, males are much larger than females, requiring more space and resources. These factors and more mean males and females are not equally distributed and are subject to different animal selection practices in research institutions. The difference is also evident in the sample size. Before data filtering, the sample size included 44,704 females and 43,413 males. After data filtering, there were 8,296 females and 3,973 males. A larger proportion of the males were filtered out of the analyses because of research-related endpoints or humane euthanasia for management reasons, reflecting bias in how sexes are deployed in research.

### Comparison with prior reports of captive NHP lifespan

As mentioned in the introduction, captive baboon maximum lifespan has been reported as 37.5 years,^5–8^ and median lifespan as 21^10^ or 11^11^ years. Our median lifespan findings align with the lowest of those estimates, and close inspection of the methods used to arrive at that estimate reveals that the study employed similar inclusion and exclusion criteria as the current study.^11^ The 37.5 year estimate is based on a single zoo baboon^9^ and is a rare case of extreme maximum longevity. The 21-year baboon lifespan estimate uses different methods from the current study, such as inclusion of live animals as right censored datapoints.^10^ In another report that includes 4,480 zoo baboons, male *P. hamadryas* were estimated to live 13.2 years and females 17.1 years from birth.^33^ We expect that this difference is due to both methodological differences in calculating median lifespan and differences in the veterinary care for the small numbers of baboons in zoo settings, e.g., they frequently receive long-term treatment for chronic diseases. It may also be due to differences between hamadryas and the mixed baboons in our study. Prior reports of lifespan of rhesus macaques have hovered around a median lifespan of 25 years and maximum 40 years, but again, these studies employed right censored data approaches.^40,54–56^ In contrast, our median lifespan estimate for rhesus is 7.9 years in males and 10.3 years in females using data only from animals with known ages at death, rather than including ages from still living animals with a right censored approach. To highlight this methodological difference, we provide survivorship probabilities with censored data for reference (**Supplementary Figure S4**). A prior study of common marmosets at a single institution estimated median lifespan of 6.5 years in animals that survived to at least two years (compared with our starting age of 1.5 years).^24^ Another marmoset study from a different institution estimated median lifespan at four years in marmosets that survived for 60 days; the same study reported cotton-top tamarin median life expectancy of 7.2 years.^57^ Our estimates from marmosets at 4 different institutions are 5.3 years in females and 6.0 years in males. For cotton-top tamarins, our estimates of median lifespan (from animals living at one institution) are 9.6 years for males and 8.9 years for females. Chimpanzee median survival in a biomedical research population has been reported as 31.0 years in males and 38.8 years in females among individuals who reached 1 year of age.^58^ In a zoo population, male chimpanzees lived a median of 26.0 years and females 30.5 years from birth.^33^ Our estimates are 33.0 years in males and 44.0 years in females among individuals who reached ten years of age and are therefore fairly consistent with previous reports. For coppery titi monkeys, median lifespan has been reported as 14.9 years in males and 11.4 years in females among individuals surviving to 31 days,^18^ compared with our estimates of 8.6 years for males and 9.2 years for females. Once again, the differences between estimates in our studies and prior reports likely arise methodologically, such as choices made about age of inclusion and use of a right censored approach to include individuals still alive and/or those euthanized for research-related endpoints. A major strength of the current study is the use of uniform methods across 12 different NHP species.

### Importance of data filtering

This study highlights the necessity of thorough methodological documentation in NHP lifespan studies. As illustrated with our primary and secondary analyses, filtering and methodological decisions impact the results and interpretation. The simplest example is the minimum age threshold for computing the survivorship functions. Including juveniles dramatically lowers median lifespan due to high rates of juvenile mortality among primates. Additionally, by including only animals that were born and died at the same institute, it sometimes eliminated the oldest known individuals from the dataset, such as two 19-year-old SNPRC marmosets; however, these instances were rare in our very large sample. Decisions that greatly reduced our analysis sample size, such as date-of-birth (DOB) cutoffs, are a privilege of a large initial (pre-filtered) dataset. So, while the DOB cutoffs greatly reduced our final sample size, it removed bias associated with very early deaths (since our dataset did not include currently alive animals). Overall, given the impact of filtering decisions, we emphasize the need for robust reporting of the decision criteria in NHP survival studies. We encourage authors to follow the ARRIVE guidelines (Animal Research: Reporting of In Vivo Experiments; https://arriveguidelines.org/), a checklist for full and transparent reporting aimed at improving rigor, transparency, and reproducibility in animal research.^59^ In longevity research, it is particularly crucial to report inclusion and exclusion criteria in addition to the details of statistical approaches.

### Limitations

One limitation of the study is that the stringent inclusion criteria reduced our starting sample size by 86%. This was necessary to ensure appropriate comparisons across institutions and species. For example, some species (cynomolgus, pigtails, baboons) have a very high percentage of deaths by research sacrifice, rather than by natural or health-related causes. Including research-related deaths as right censored data results in highly skewed models with limited utility for these species (e.g., survival curves for female baboons do not converge past the median survivorship when including censored data). Further, censoring was biased by sex because of the differences in research utilization and breeding needs, statistically hindering the possibility of comparisons between males and females. Therefore, primary analyses were limited to data from natural or clinical deaths, eliminating the need for right censoring. Another constraint of the study is our limited knowledge of specific cause of death. Differences in institutional death coding systems make it difficult to easily determine cause of death, since some record systems group many types of deaths, while others have more granular codes to distinguish among death types. Furthermore, as previously described, variations in institutional practices can likely impose some differences on lifespan. While inclusion and assessment of specific practices (e.g., housing) are not explored within this study, institutional source was included within regression models to adjust for these potential effects.

## Conclusions

The need for comparative analyses of lifespans across species has been widely acknowledged.^60^ Investigators need access to reliable lifespan tables, survivorship graphs, and maximum lifespan measurements to conduct relevant translational aging studies. Here we provide the largest dataset yet assembled from captive research NHPs. These data provide a valuable comparative resource for translational NHP research, primary data on multispecies NHP lifespan in captivity, and context for consideration of morbidity and mortality in the study of diverse diseases.

## Acknowledgements & Sources of Funding

This work was supported by the National Institutes of Health: P40-OD010965 (MJJ), P51-OD011133 (CR; SNPRC), P51-OD011106 (RC; WNPRC), P51-OD011103 (EJV;NEPRC), P51-OD011104 (EJV,TNPRC), U42OD011123 (CEH; WaNPRC), P51OD010425 (CEH; WaNPRC), P51OD011092 (KC), OD011107 (KB), and the National Institute on Aging: R01AG087957 (EEQ), U19AG057758 (LAC), U34AGAG068482 (AS), P30AG013319 (AS), P30AG044271 (AS), R01AG050797 (AS), AG-067419 (BH), NIH-NIA Intramural Research Program (JAM), R24AG073199 (CS). Computations were performed using the Wake Forest University (WFU) High Performance Computing Facility, a centrally managed computational resource available to WFU researchers including faculty, staff, students, and collaborators.

## Data Availability

Raw, de-identified data are available via the password-protected database MIDAS (Monkey Inventory and DAta management of Samples), request for access available from https://midas.wakehealth.edu/MIDAS. The MIDAS database provides tools for species comparisons, which will make this a user-friendly resource accessible to researchers. Data accessible to approved users within MIDAS includes de-identified animal-level information used for analyses (e.g., date of birth, date of death, species, sex), as well as summary-level data such as the survivorship probabilities calculated in primary and secondary analyses. Data from all NHP in the manuscript are available within MIDAS, with the exceptions of data from chimpanzees and data from NHP residing at the Primate Research Center in Indonesia. Approved users who seek additional data not available within MIDAS should contact the authors directly. Data sharing will be limited to scientific uses.

## Code Availability

Analyses and summaries were computed using functions and libraries, as described in methods, in accordance with standard practices and their vignettes. Custom Code for fitting exponential curves to survival data is available in Supplementary Information and is available via MIDAS as described in Data Availability.

## Ethical Statement/Conflict of Interests

The authors declare no competing interests.

