## Supplemental document for "Comparative lifespan and healthspan of nonhuman primate species common to biomedical research"

**Supplementary Tables and Figures**

### **Table S1.** Number of animals contributed by each institution, by species, prior to sample filtering.

| **Common Name** | **Species name** | **Total** | **Barshop** | **CNPRC** | **ENPRC** | **Indonesia** | **Keeling** | **NEPRC** | **NIA** | **ONPRC** | **SNPRC** | **TNPRC** | **VRC** | **WaNPRC** | **WNPRC** | **Yale** |
| --- | --- | --- | --- | --- | --- | --- | --- | --- | --- | --- | --- | --- | --- | --- | --- | --- |
| Baboon | *P. hamadryas* spp. | 15893 | 0 | 42 | 0 | 0 | 0 | 188 | 0 | 213 | 13980 | 0 | 0 | 1470 | 0 | 0 |
| Bonnet macaques | *M. radiata* | 266 | 0 | 266 | 0 | 0 | 0 | 0 | 0 | 0 | 0 | 0 | 0 | 0 | 0 | 0 |
| Chimpanzee | *P. troglodytes* spp. | 581 | 0 | 0 | 27 | 0 | 356 | 0 | 0 | 0 | 185 | 13 | 0 | 0 | 0 | 0 |
| Common marmoset | *C. jacchus* | 6898 | 468 | 0 | 0 | 0 | 0 | 3237 | 0 | 0 | 1574 | 0 | 0 | 0 | 1619 | 0 |
| Coppery titi monkey | *P. cupreus* | 304 | 0 | 304 | 0 | 0 | 0 | 0 | 0 | 0 | 0 | 0 | 0 | 0 | 0 | 0 |
| Cotton-top tamarin | *S. oedipus* | 2609 | 0 | 0 | 0 | 0 | 0 | 2592 | 0 | 0 | 0 | 17 | 0 | 0 | 0 | 0 |
| Cynomolgus macaque | *M. fascicularis* | 5078 | 0 | 740 | 0 | 0 | 0 | 1218 | 0 | 205 | 2030 | 40 | 0 | 822 | 23 | 0 |
| Japanese macaque | *M. fuscata* | 1897 | 0 | 1 | 0 | 0 | 0 | 0 | 0 | 1884 | 0 | 0 | 0 | 12 | 0 | 0 |
| Pig-tailed macaque | *M. nemestrina* | 8906 | 0 | 0 | 0 | 82 | 0 | 80 | 0 | 59 | 0 | 142 | 0 | 8543 | 0 | 0 |
| Rhesus macaque | *M. mulatta* | 65042 | 0 | 17351 | 0 | 0 | 0 | 6041 | 180 | 14952 | 1282 | 21387 | 0 | 197 | 3587 | 65 |
| Squirrel monkey | *Saimiri* spp. | 1505 | 0 | 117 | 0 | 0 | 0 | 531 | 0 | 0 | 0 | 857 | 0 | 0 | 0 | 0 |
| Vervet/African green | *C. aethiops sabaeus* | 1848 | 0 | 0 | 0 | 0 | 0 | 83 | 0 | 0 | 0 | 168 | 1597 | 0 | 0 | 0 |

­­

Abbreviations: Sam and Ann Barshop Institute for Longevity and Aging Studies (Barshop); California National Primate Research Center (CNPRC); Emory National Primate Research Center (ENPRC); Primate Research Center IPB University in Indonesia (Indonesia); Keeling Center for Comparative Medicine and Research at The University of Texas MD Anderson Cancer Center (Keeling); New England Primate Research Center (NEPRC; this center is no longer open but we obtained archival data); National Institute on Aging (NIA) Intramural Research Program; Oregon National Primate Research Center (ONPRC); Southwest National Primate Research Center (SNPRC); Tulane National Primate Research Center (TNPRC); Vervet Research Colony of Wake Forest University (VRC); Washington National Primate Research Center (WaNPRC); Wisconsin National Primate Research Center (WNPRC); Yale University (Yale).

### **Figure S1:** Supportive data for date of birth cutoffs.

For analysis, data were pruned to animals born before a species-specific date of birth (DOB) cutoff. This prevented data skewing from recent deaths (in absence of information on alive animals). The 85^th^ percentile of maximum age in years was chosen to calculate the DOB cutoff (e.g., animals born before 2023 minus 85^th^ percentile were removed). Scatterplots show age at death by year of birth, including all death types. ‘Censored labels’ indicate deaths due to research sacrifice or colony management; ‘natural labels’ indicate natural/health-related causes. Tables show maximum age in years and percentiles calculated including deaths from natural causes and humane euthanasia for health reasons. For comparison, more stringent DOB thresholds are also shown (e.g., based on 95^th^, 98^th^ percentile max ages). These more stringent thresholds did not substantially alter the overall summary statistics (e.g., mean age); but did greatly reduce overall counts, motivating selection of 85^th^ percentile for DOB thresholds.

#### **Figure S1-A** Baboons

Alive animals not included. Asterisk indicates timespan is less than 5 years.

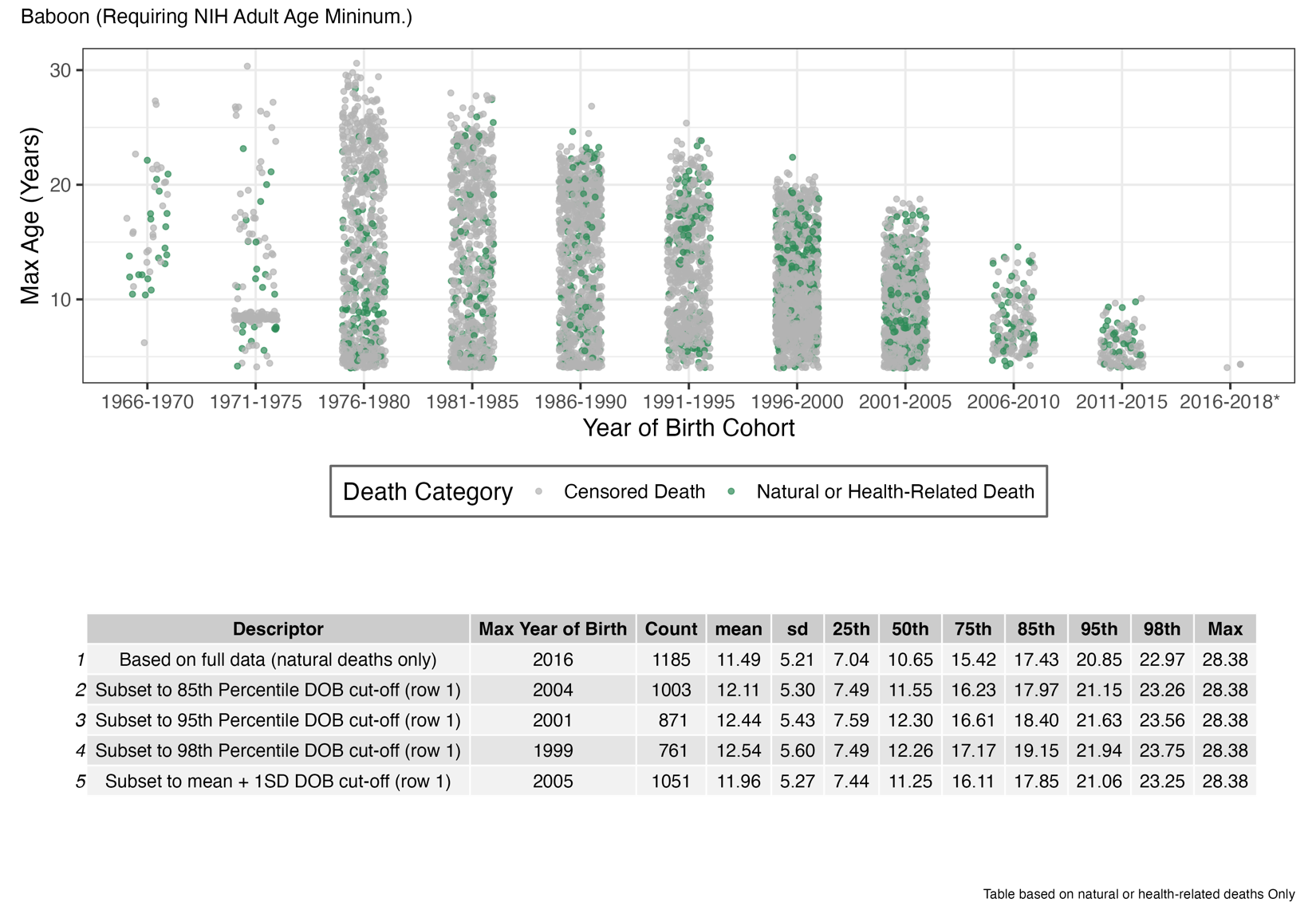

#### **Figure S1-C** Chimpanzees

Alive animals not included. Asterisk indicates timespan is less than 5 years.

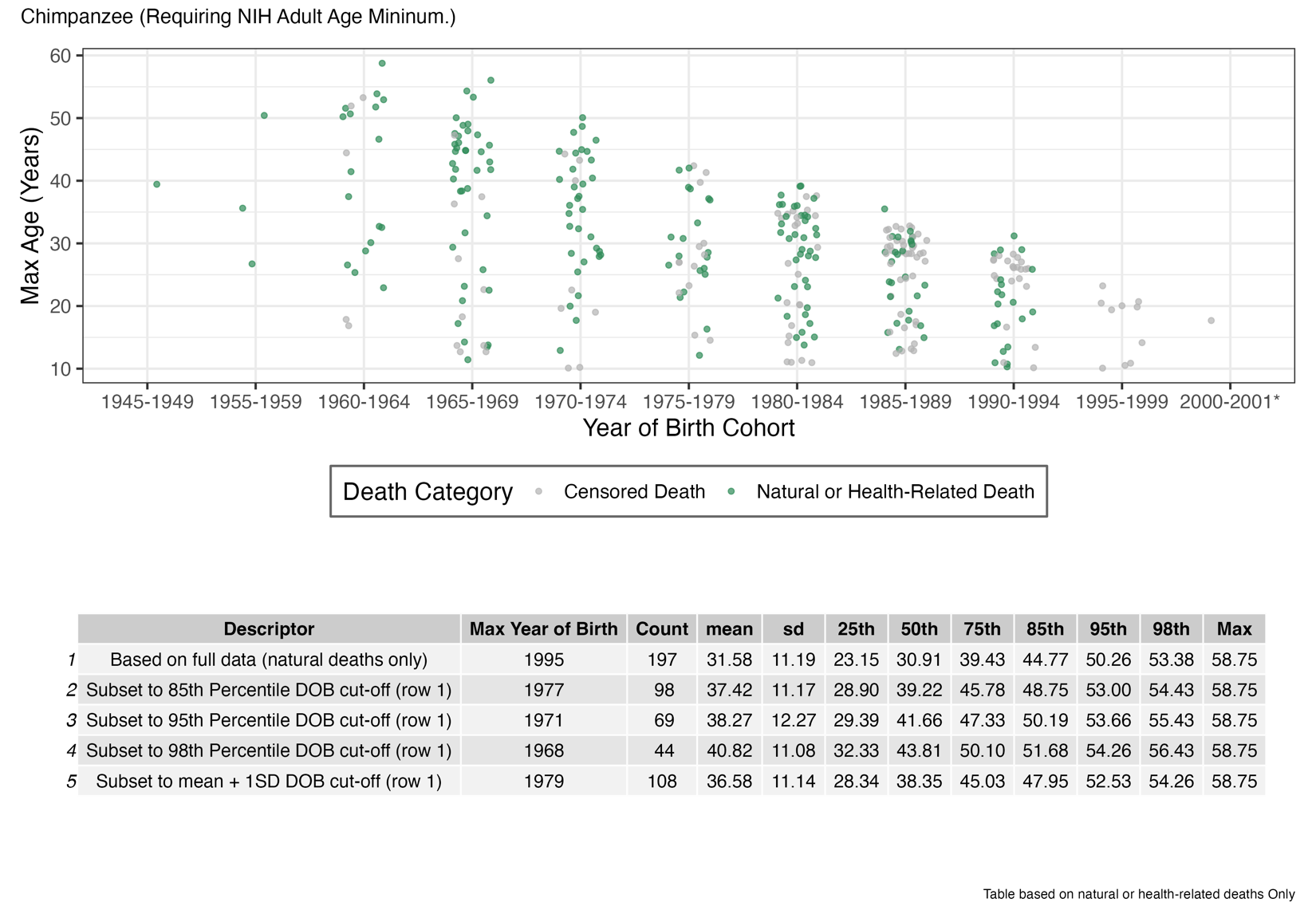

#### **Figure S1-B** Bonnet macaques

Alive animals not included.

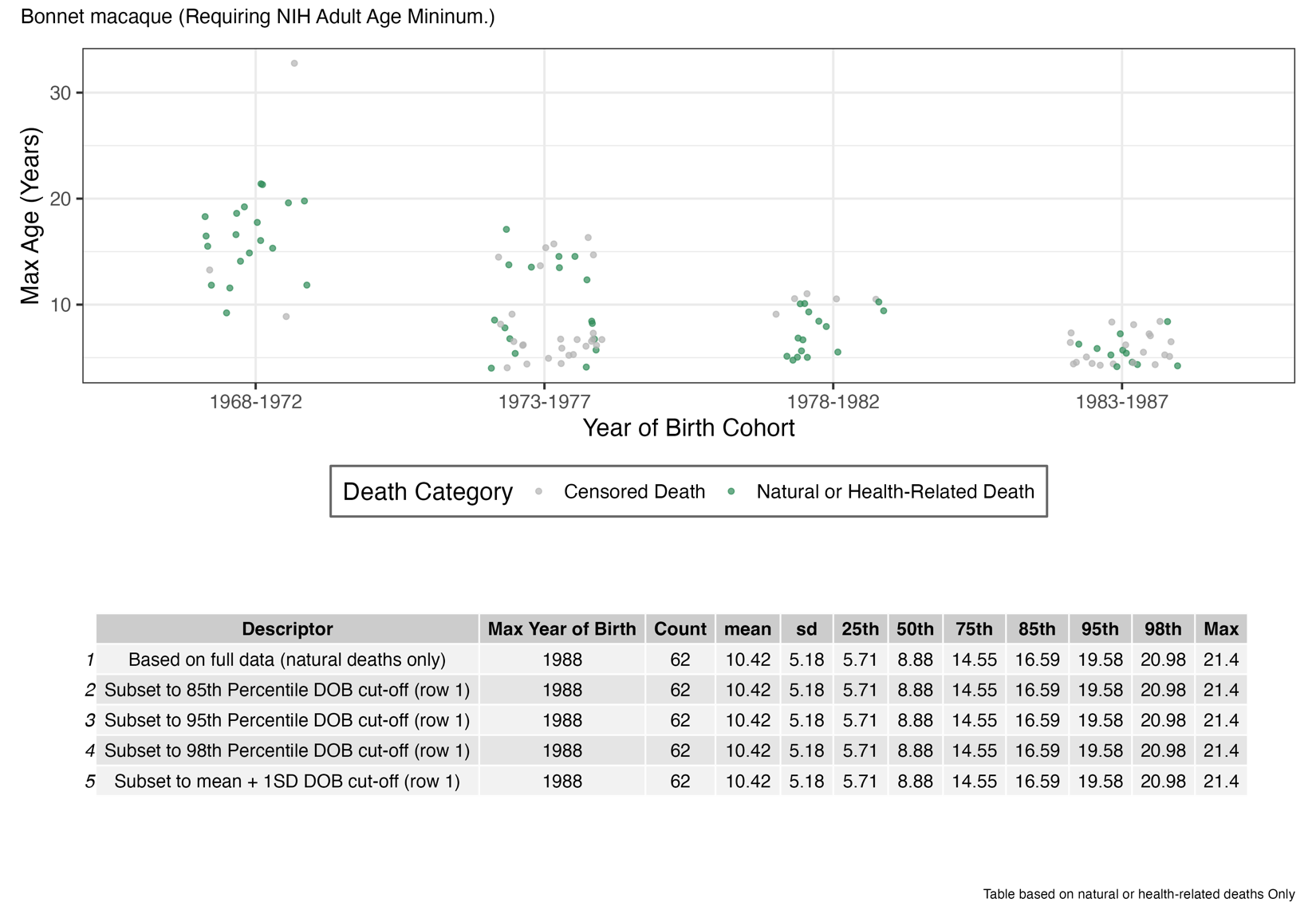

#### **Figure S1-D** Common marmoset

Alive animals not included. Asterisk indicates timespan is less than 5 years.

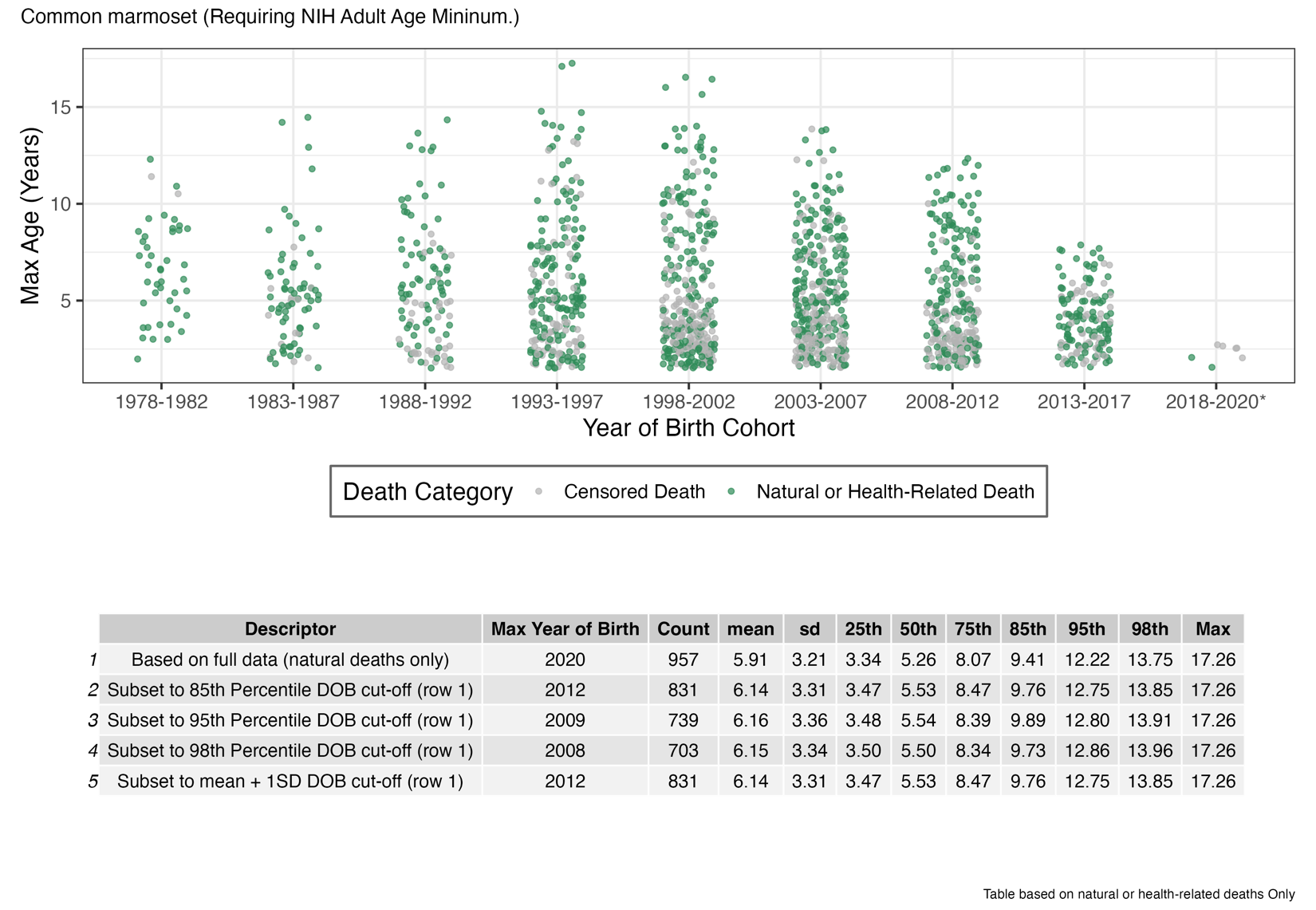

#### **Figure S1-E** Coppery titi monkeys

Alive animals not included. Asterisk indicates timespan is less than 5 years.

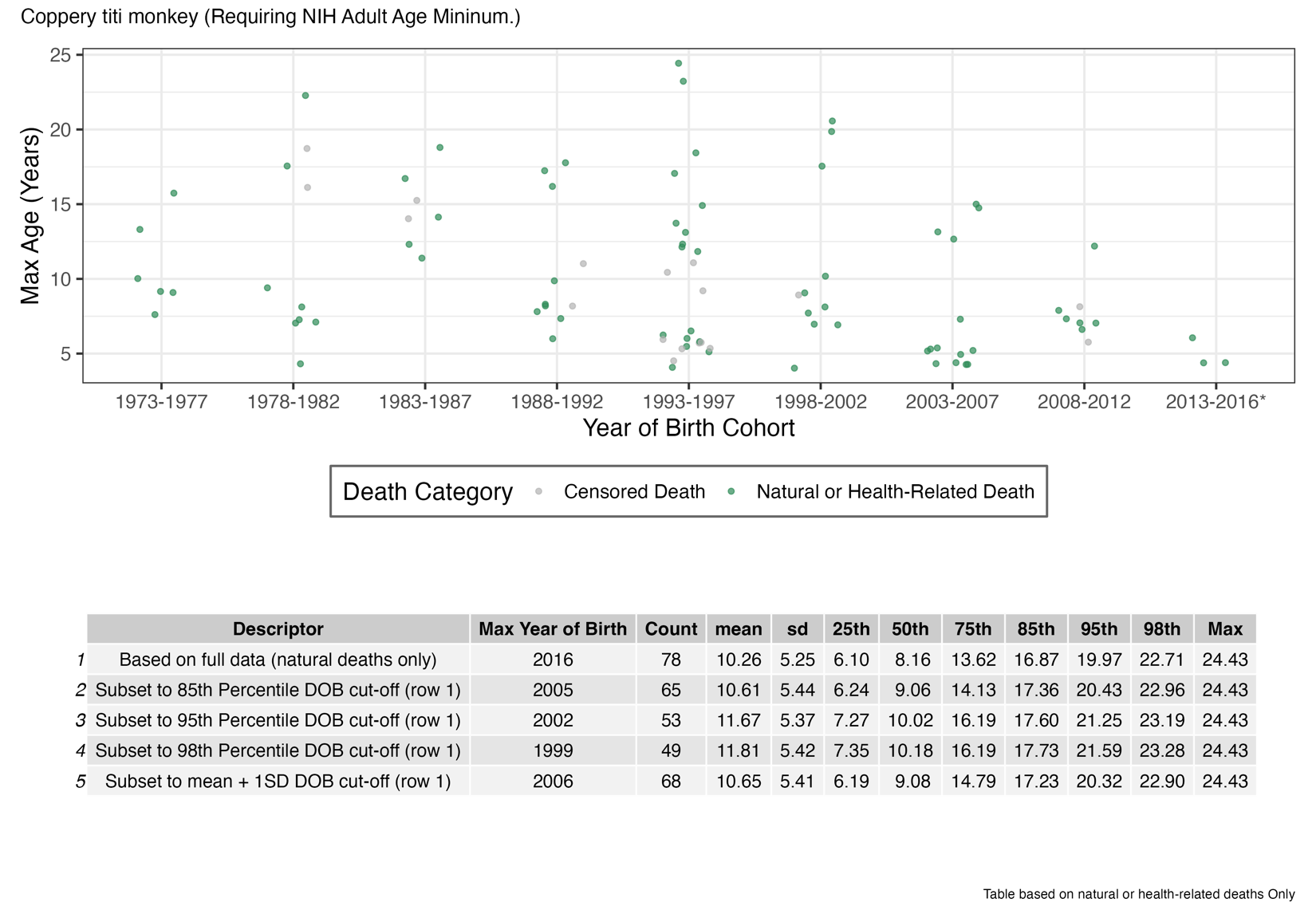

#### **Figure S1-F** Cotton-top tamarins

Alive animals not included. Asterisk indicates timespan is less than 5 years.

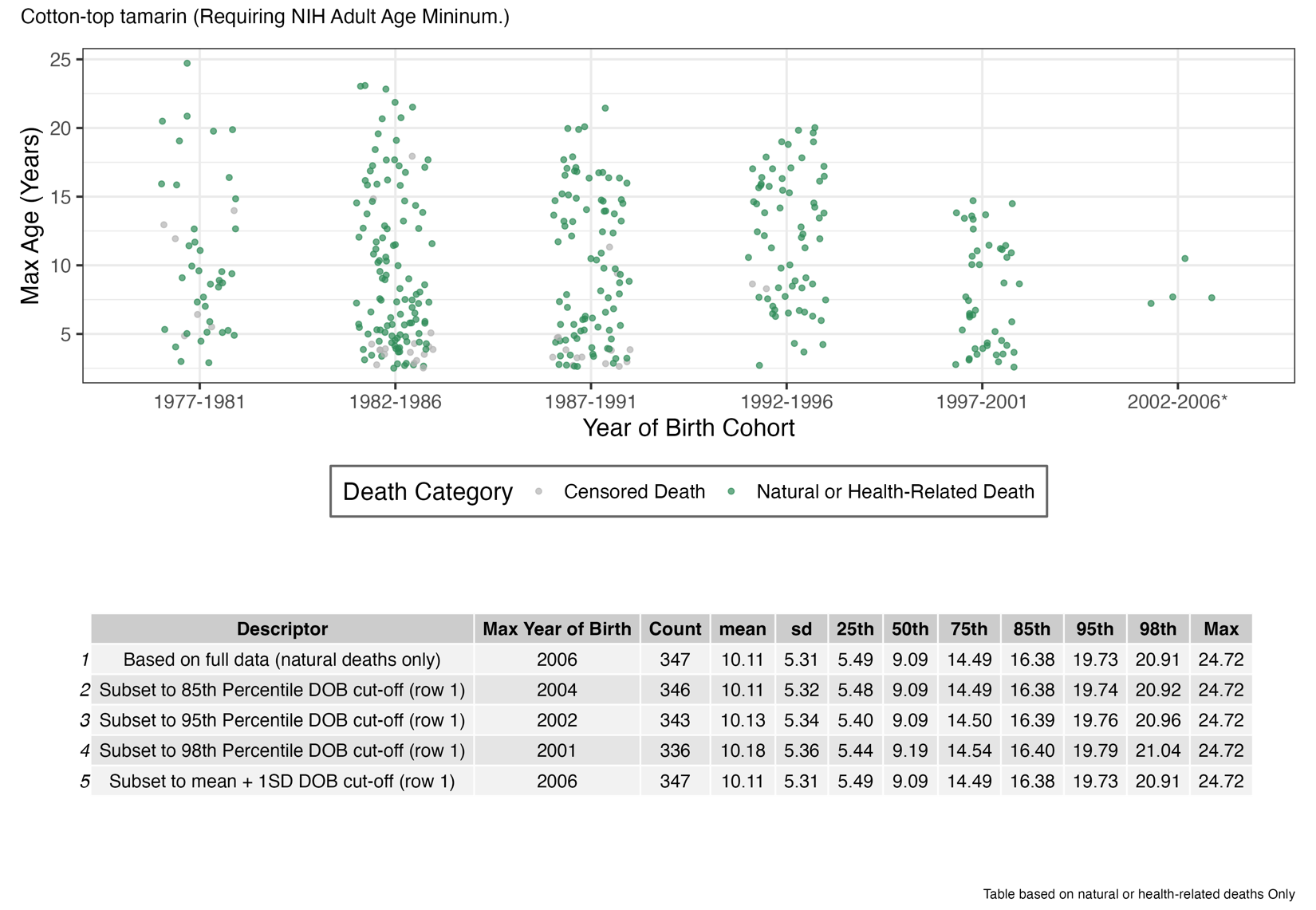

#### **Figure S1-G** Cynomolgus macaques

Alive animals not included. Asterisk indicates timespan is less than 5 years.

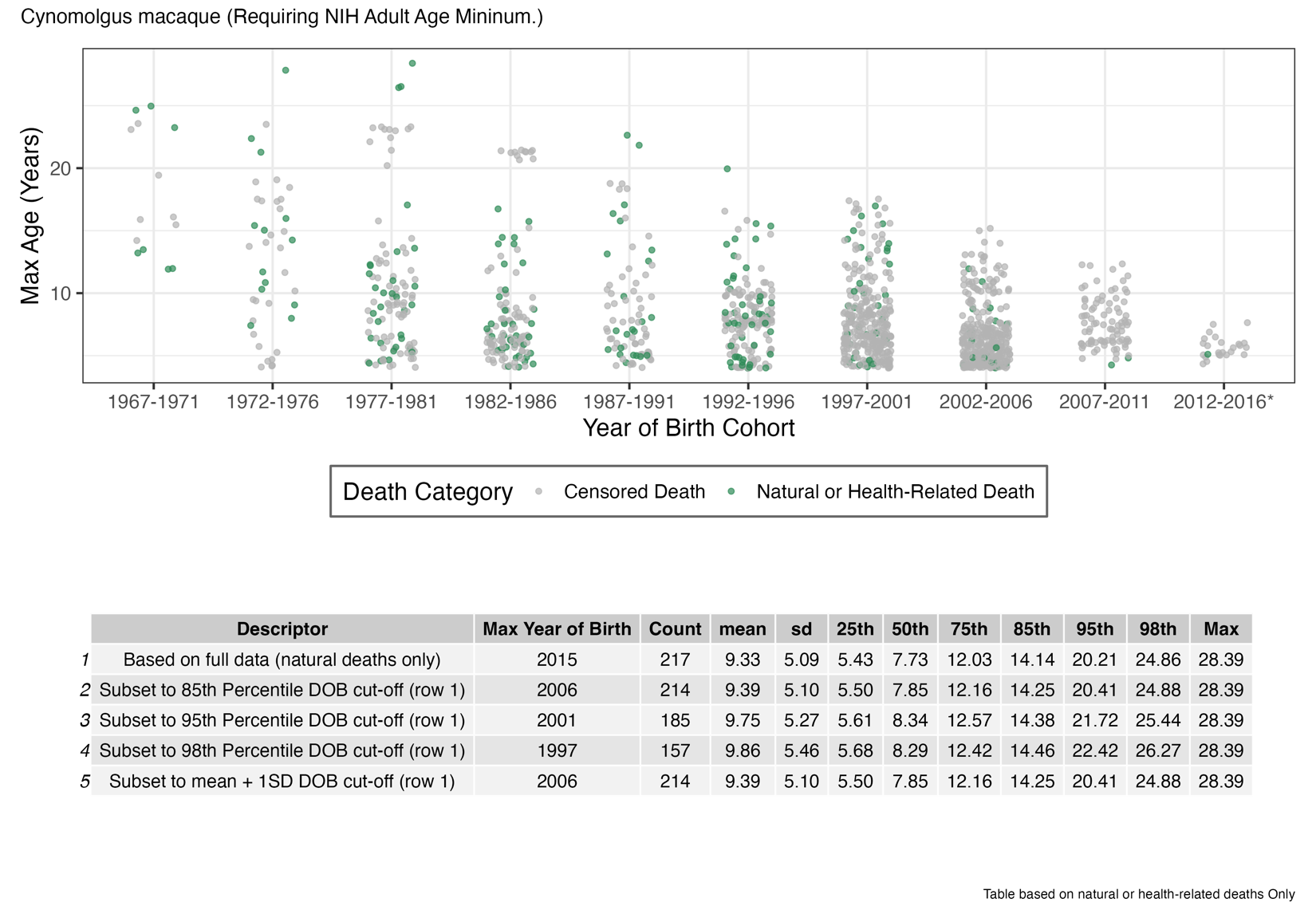

#### **Figure S1-H** Japanese macaque

Alive animals not included. Asterisk indicates timespan is less than 5 years.

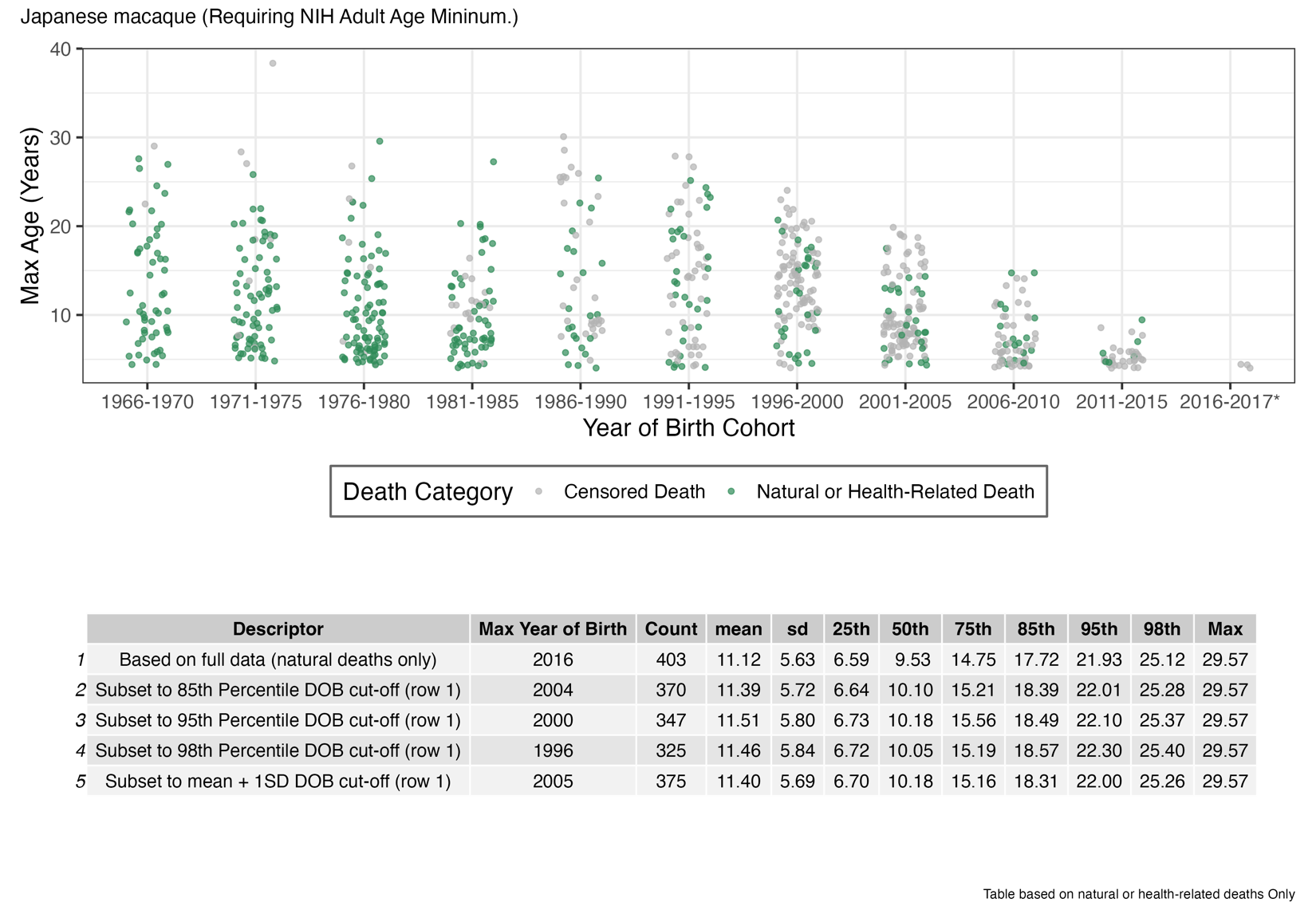

#### **Figure S1-I** Pigtail macaques

Alive animals not included. Asterisk indicates timespan is less than 5 years.

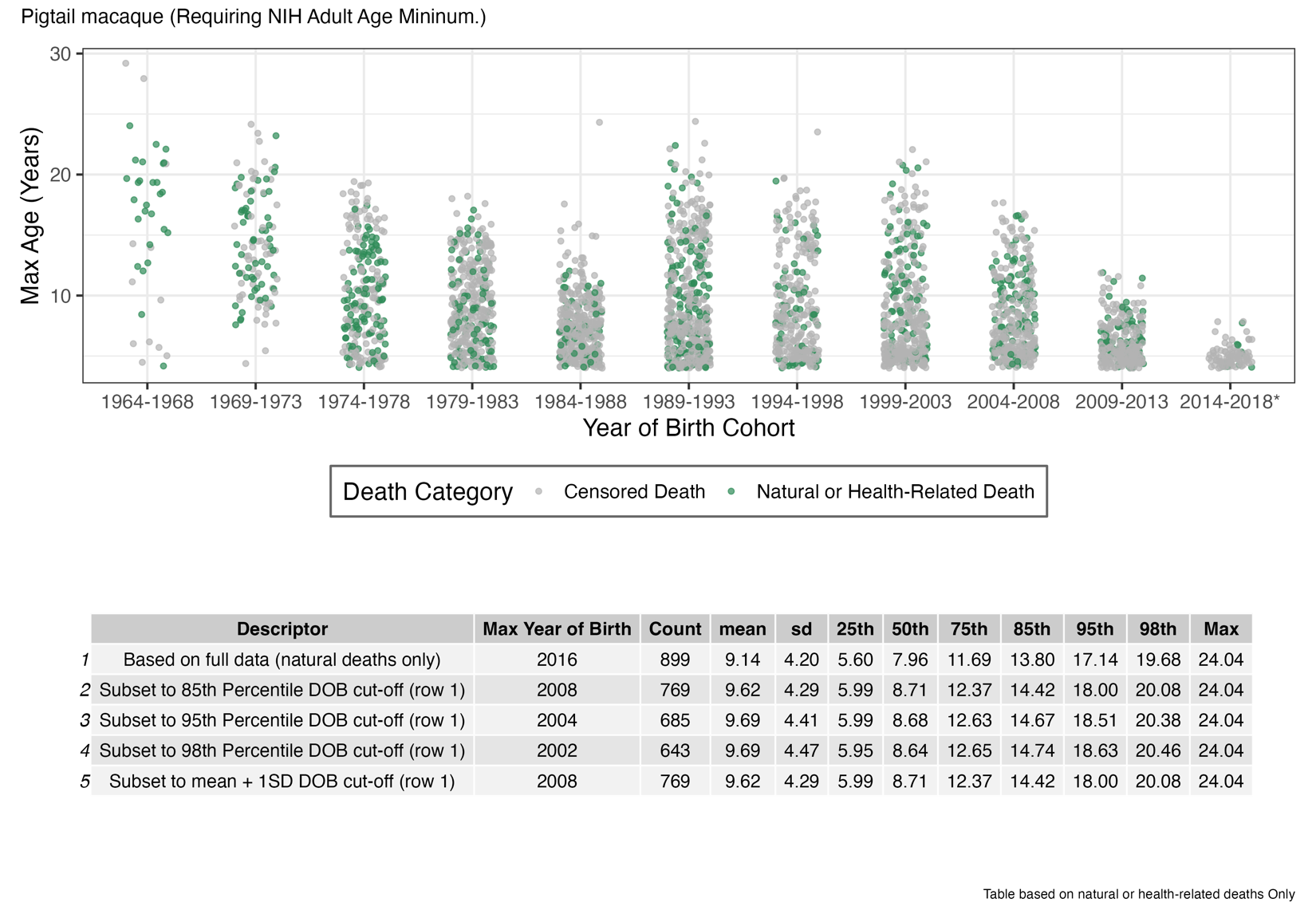

#### **Figure S1-J** Rhesus macaque

Alive animals not included. Asterisk indicates timespan is less than 5 years.

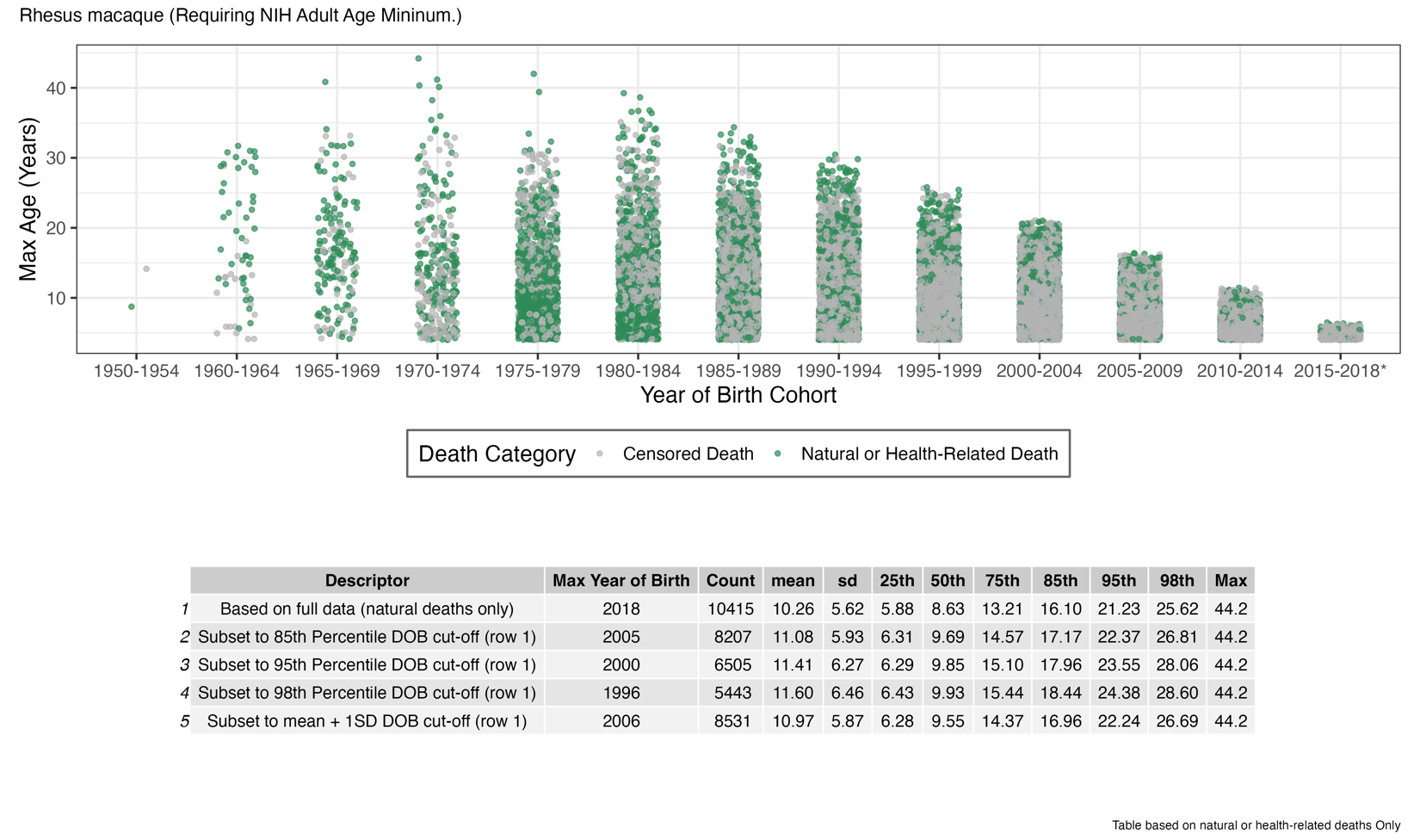

#### **Figure S1-K** Squirrel monkeys

Alive animals not included. Asterisk indicates timespan is less than 5 years.

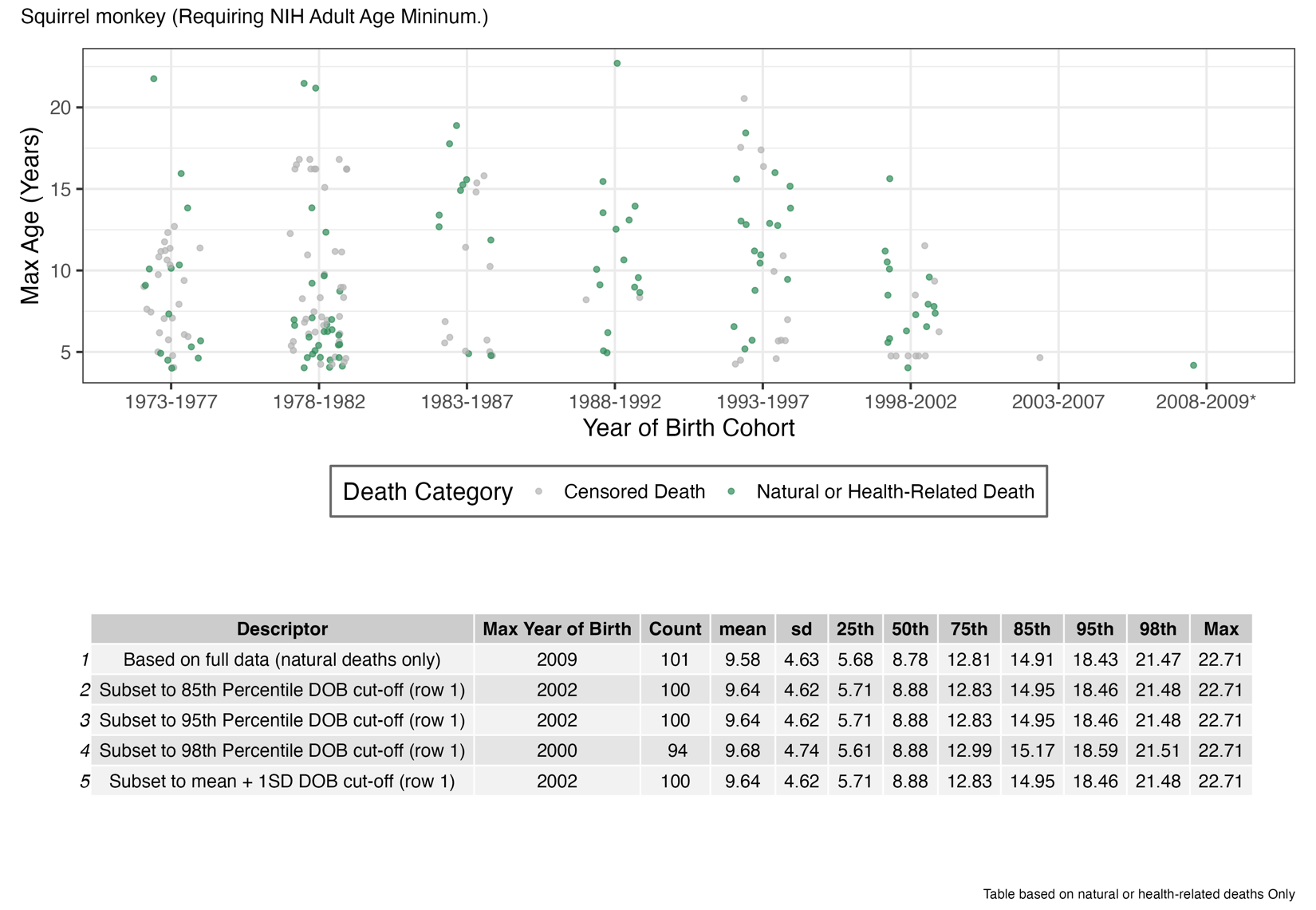

#### **Figure S1-L** Vervets/African green monkeys

Alive animals not included. Asterisk indicates timespan is less than 5 years.

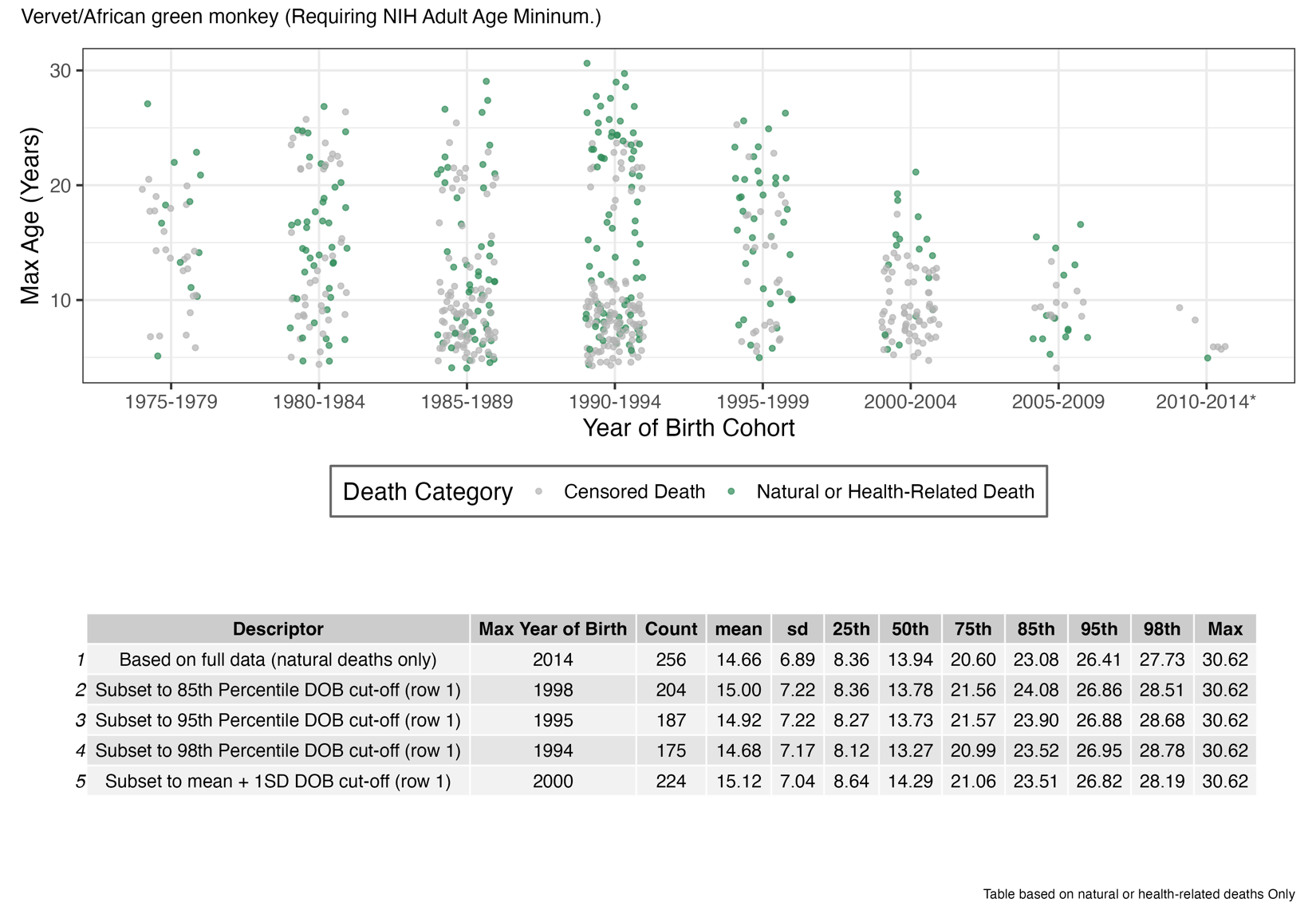

### **Figure S2.** Exponential models fit to first and last quartiles of Kaplan-Meier survival curves.

Exponential models are shown as dashed lined. Beta parameters from fitted models are shown. Survival curves are based on natural and health-related deaths, animals surviving to at least NIH-defined adulthood, and born before the species-specific DOB cut-off.

#### **Figure S2-A** Baboons

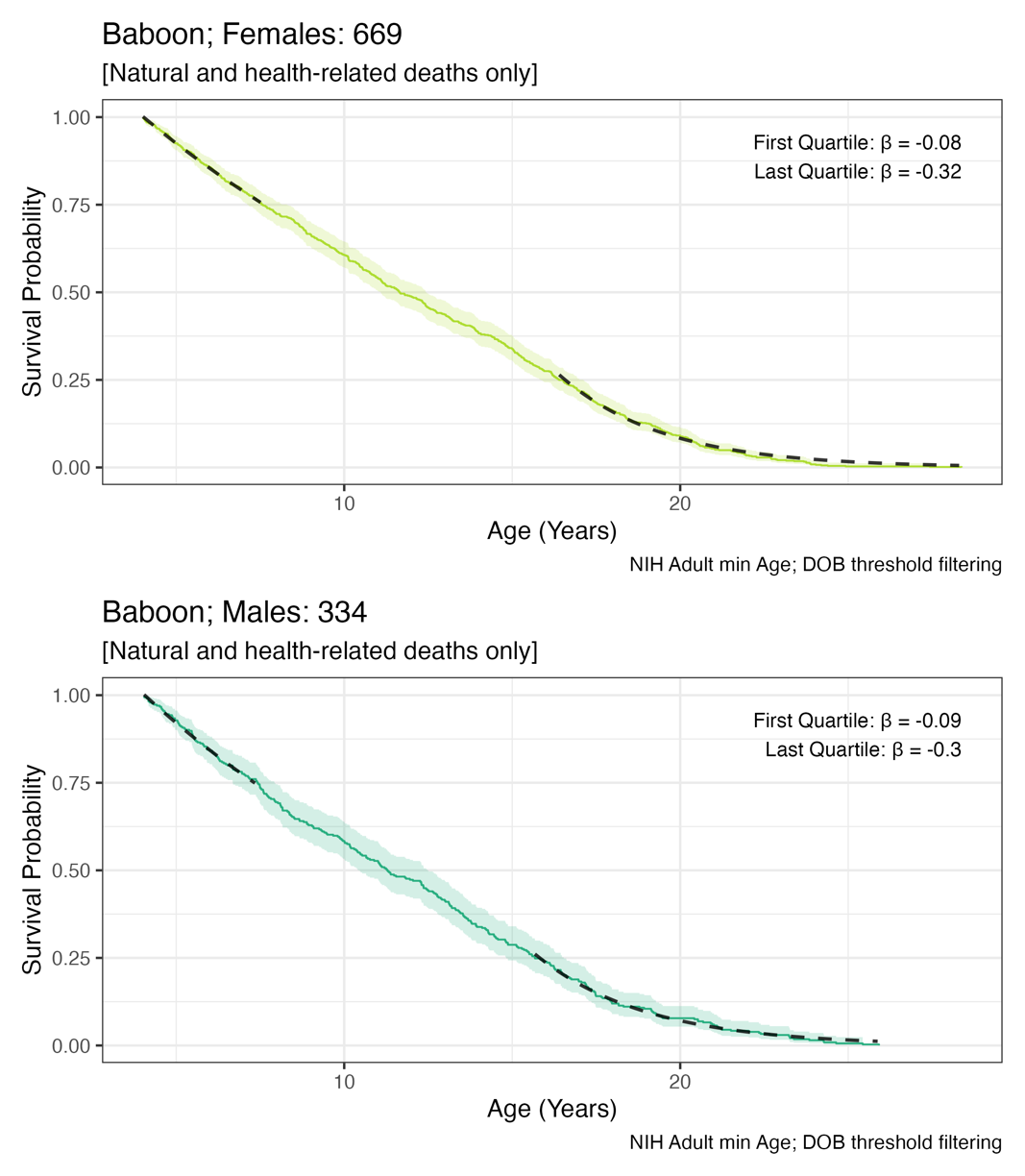

#### **Figure S2-B** Bonnet macaques

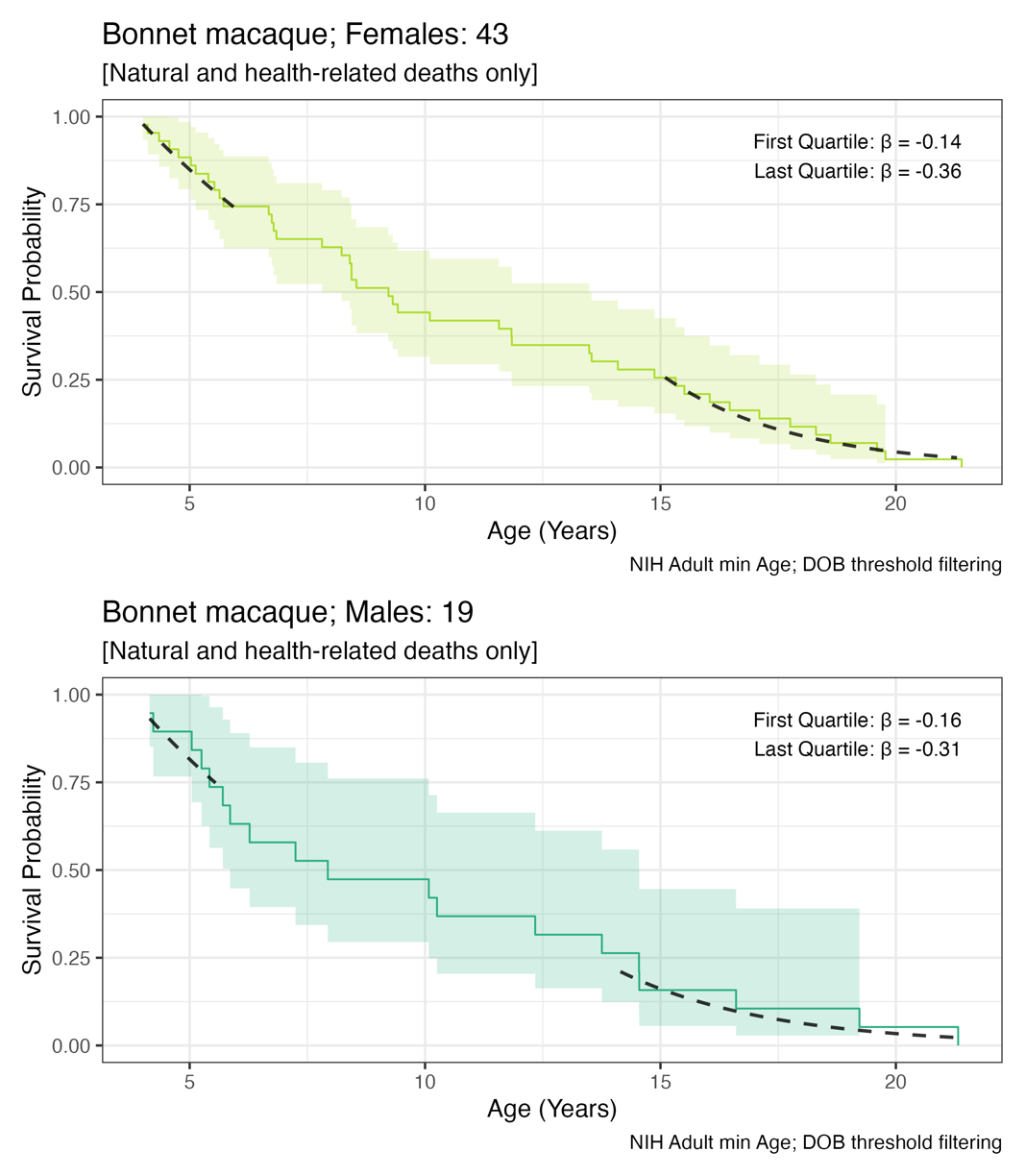

#### **Figure S2-C** Chimpanzees

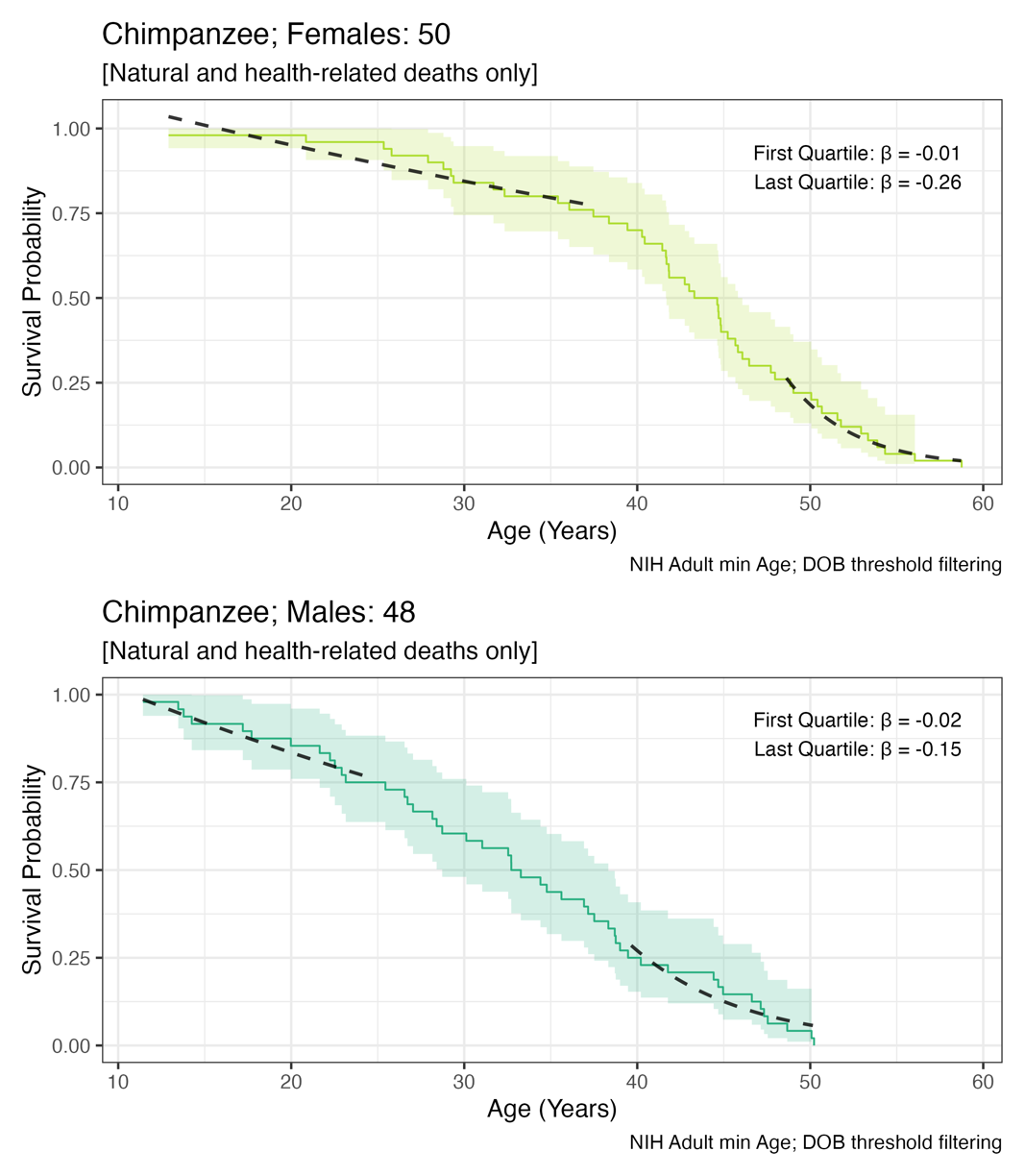

#### **Figure S2-D:** Common marmoset

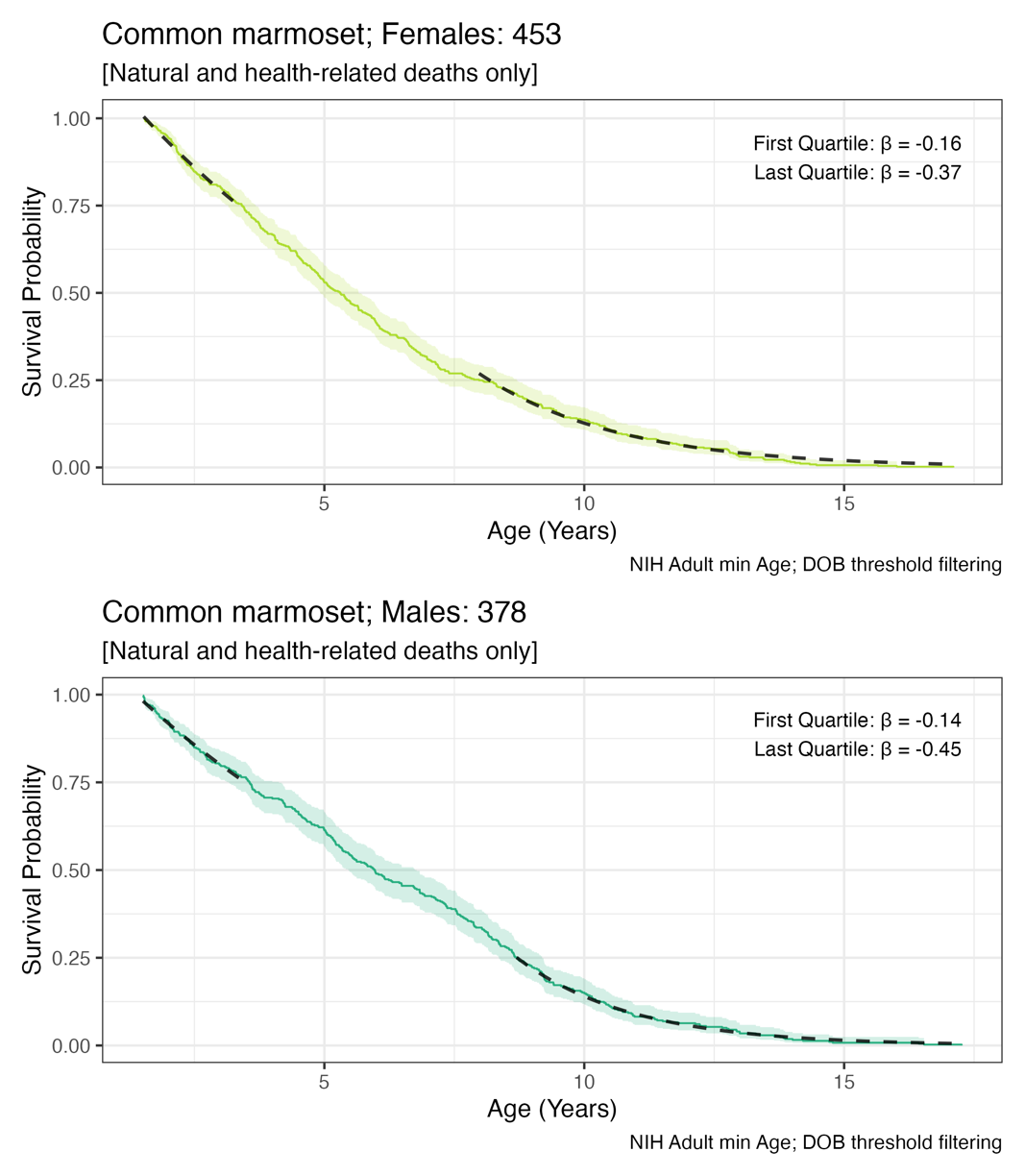

#### **Figure S2-E** Coppery titi monkeys

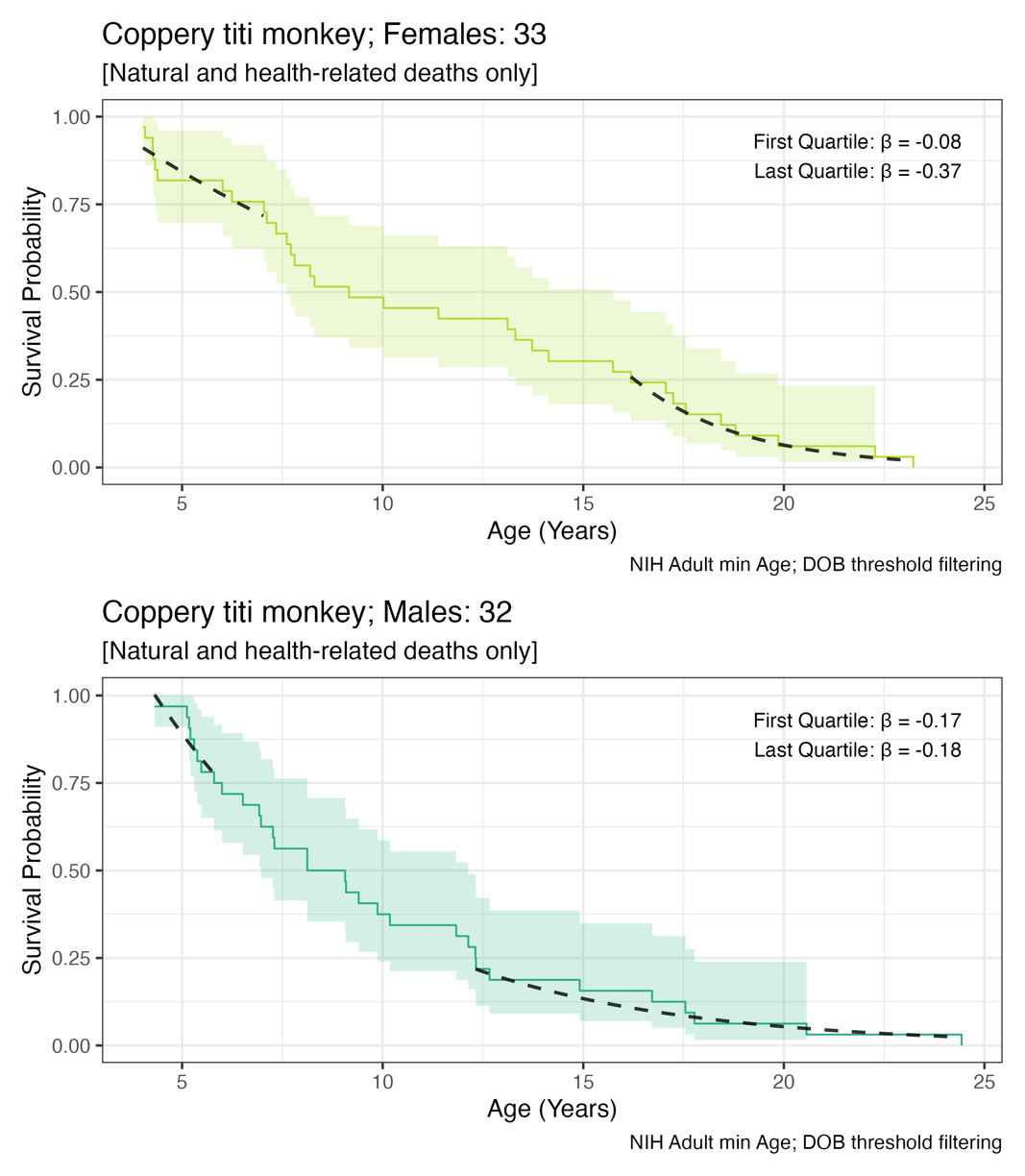

#### **Figure S2-F** Cotton-top tamarins

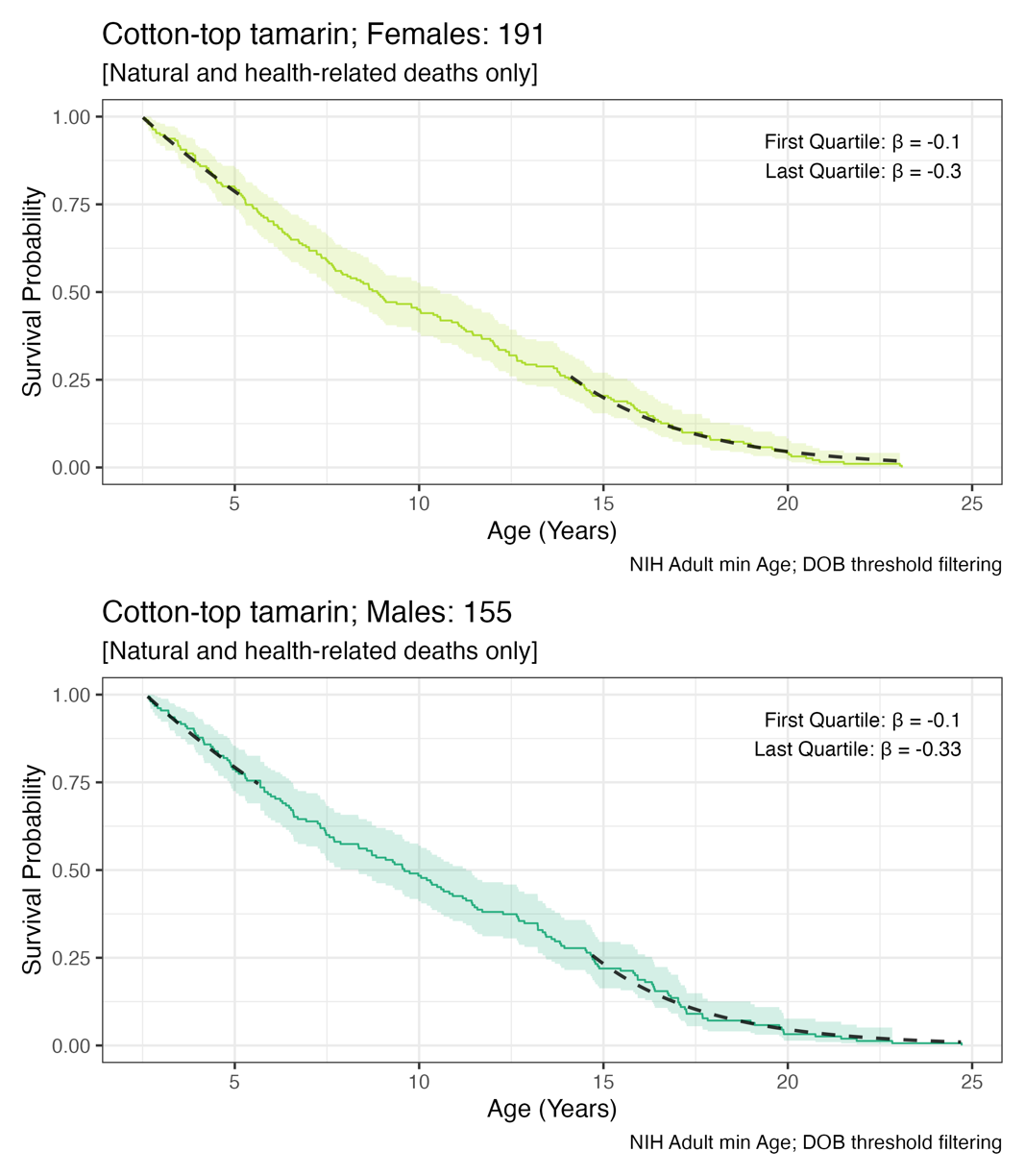

#### **Figure S2-G** Cynomolgus macaques

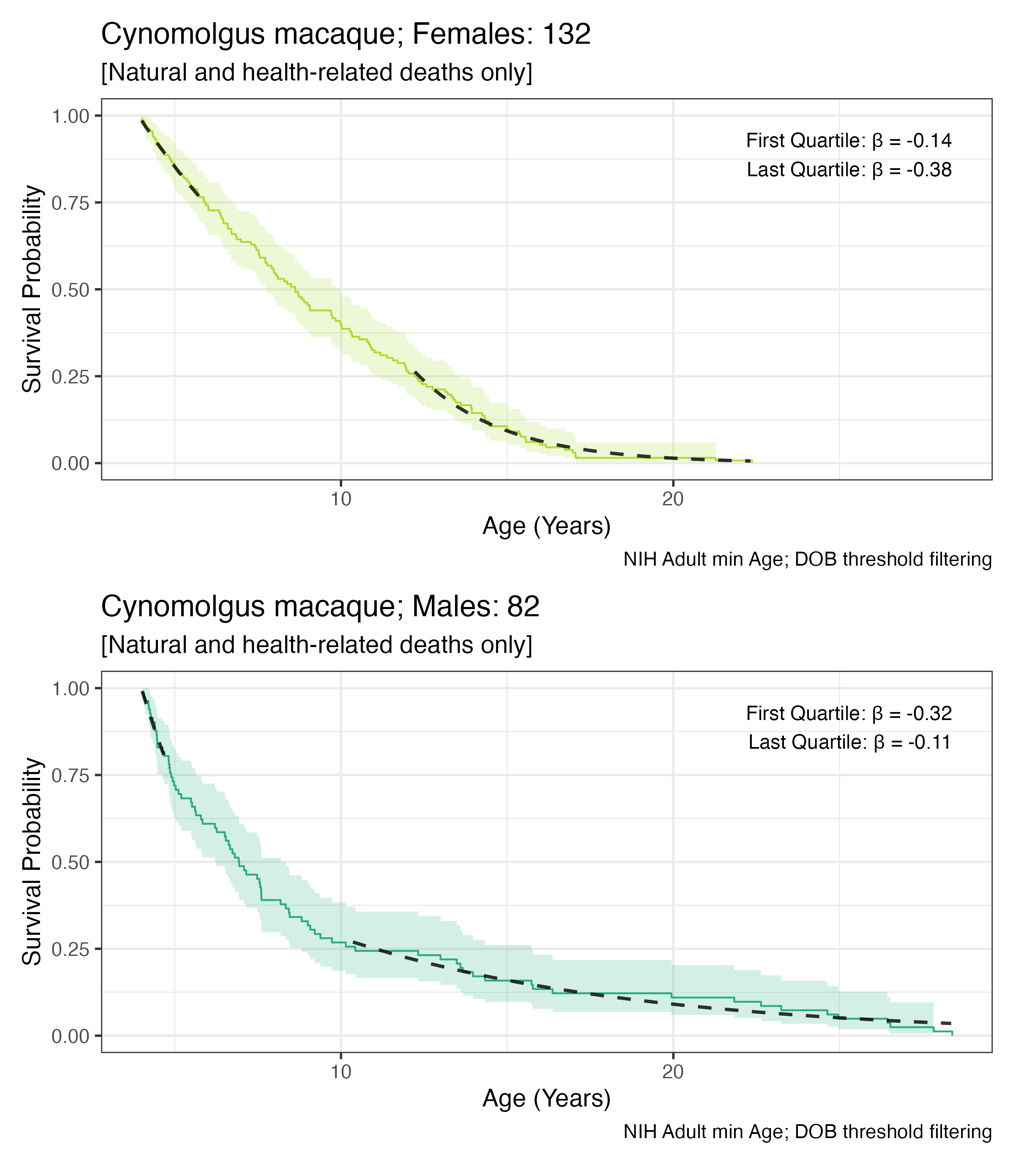

#### **Figure S2-H** Japanese macaques

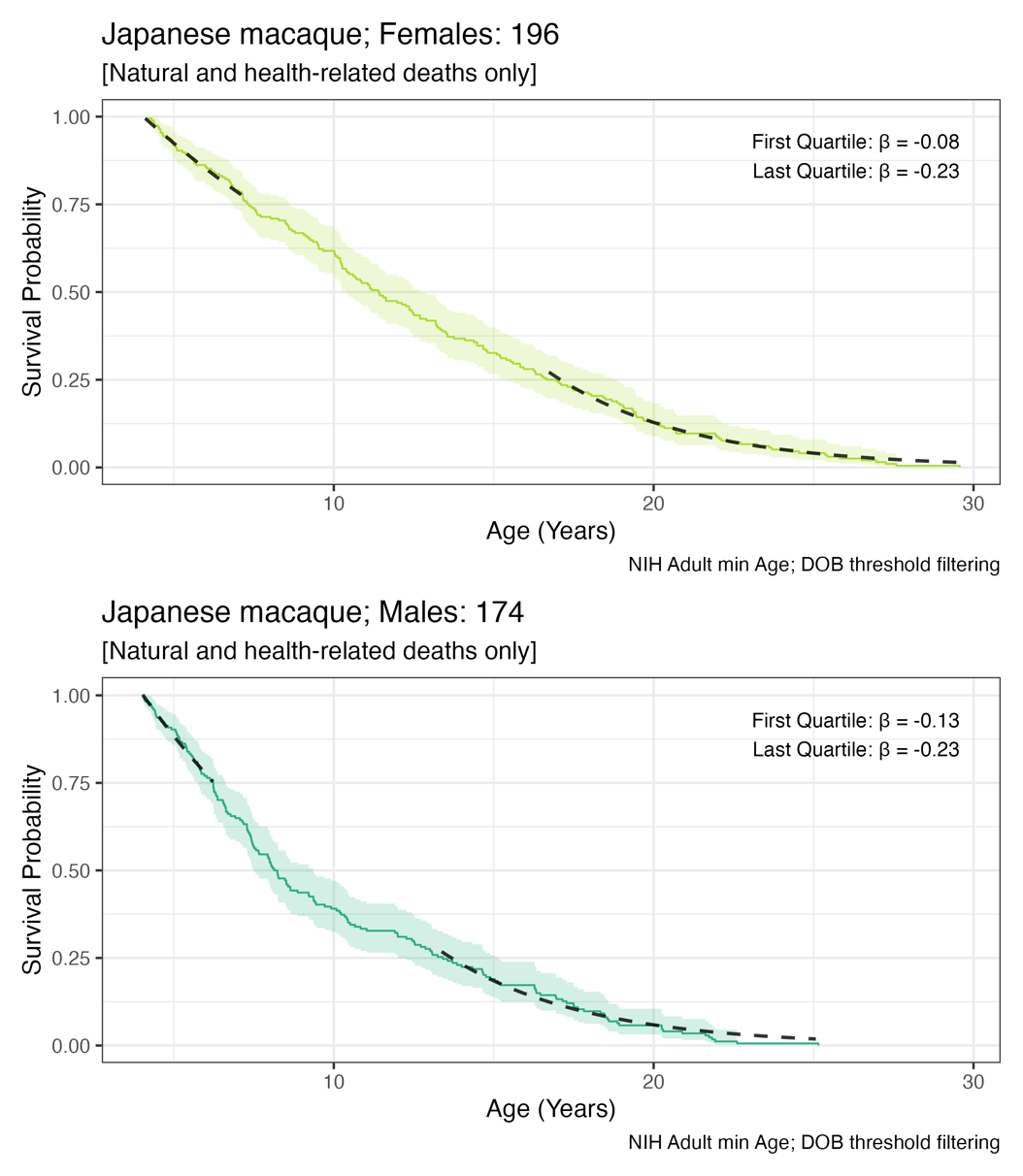

#### **Figure S2-I** Pigtail macaques

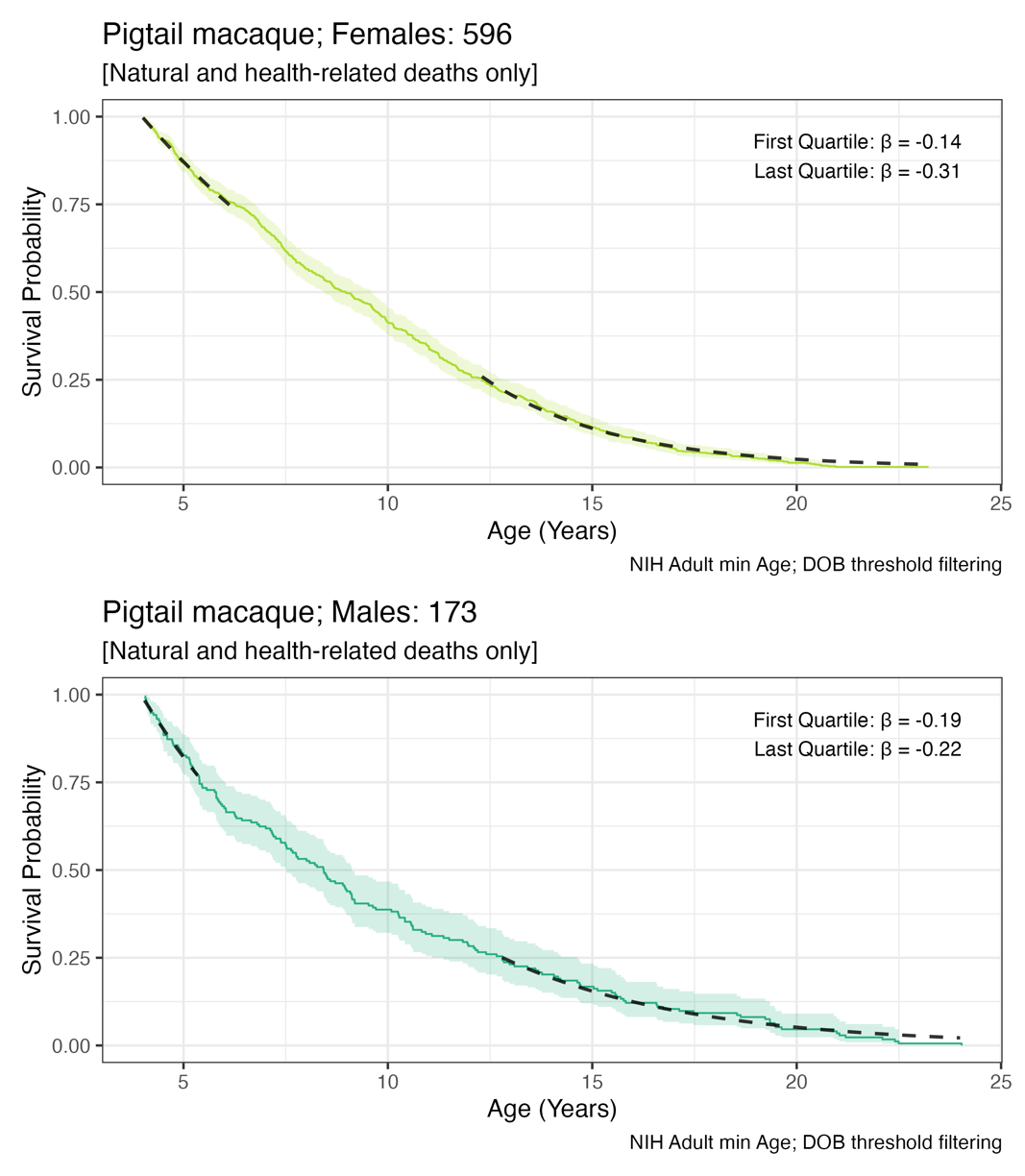

#### **Figure S2-J** Rhesus macaques

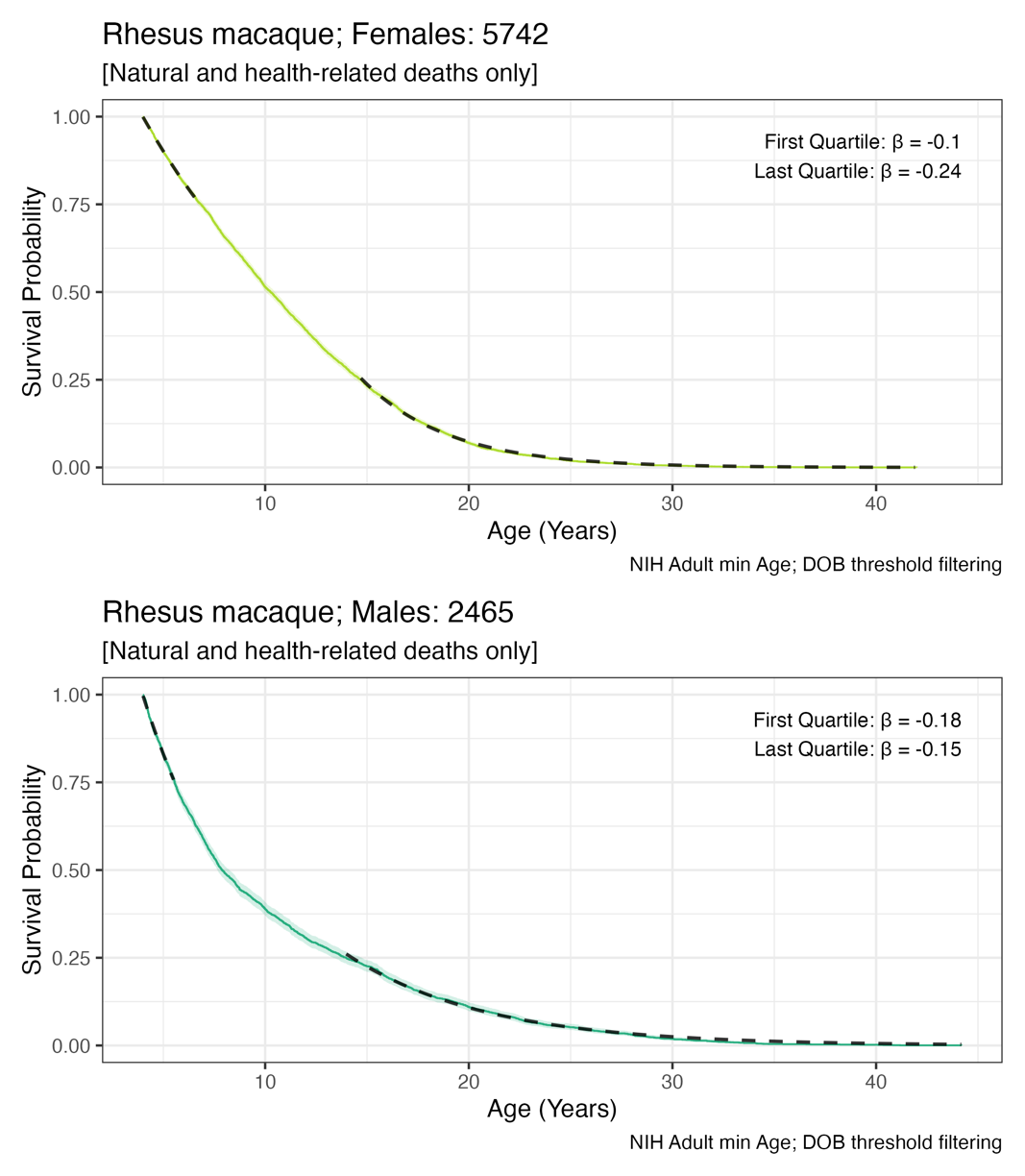

#### **Figure S2-K** Squirrel monkeys

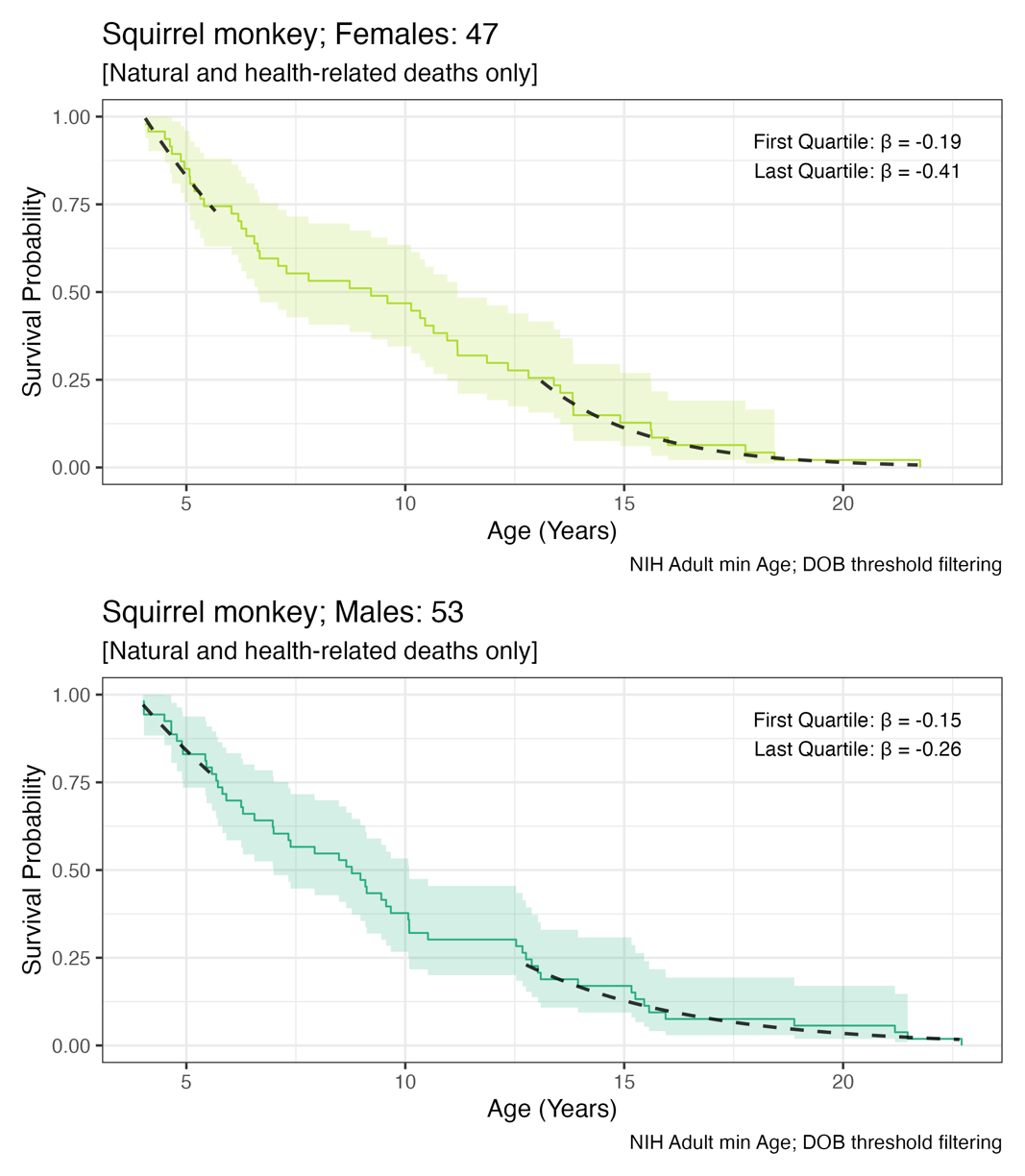

#### **Figure S2-L** Vervets/African green monkeys

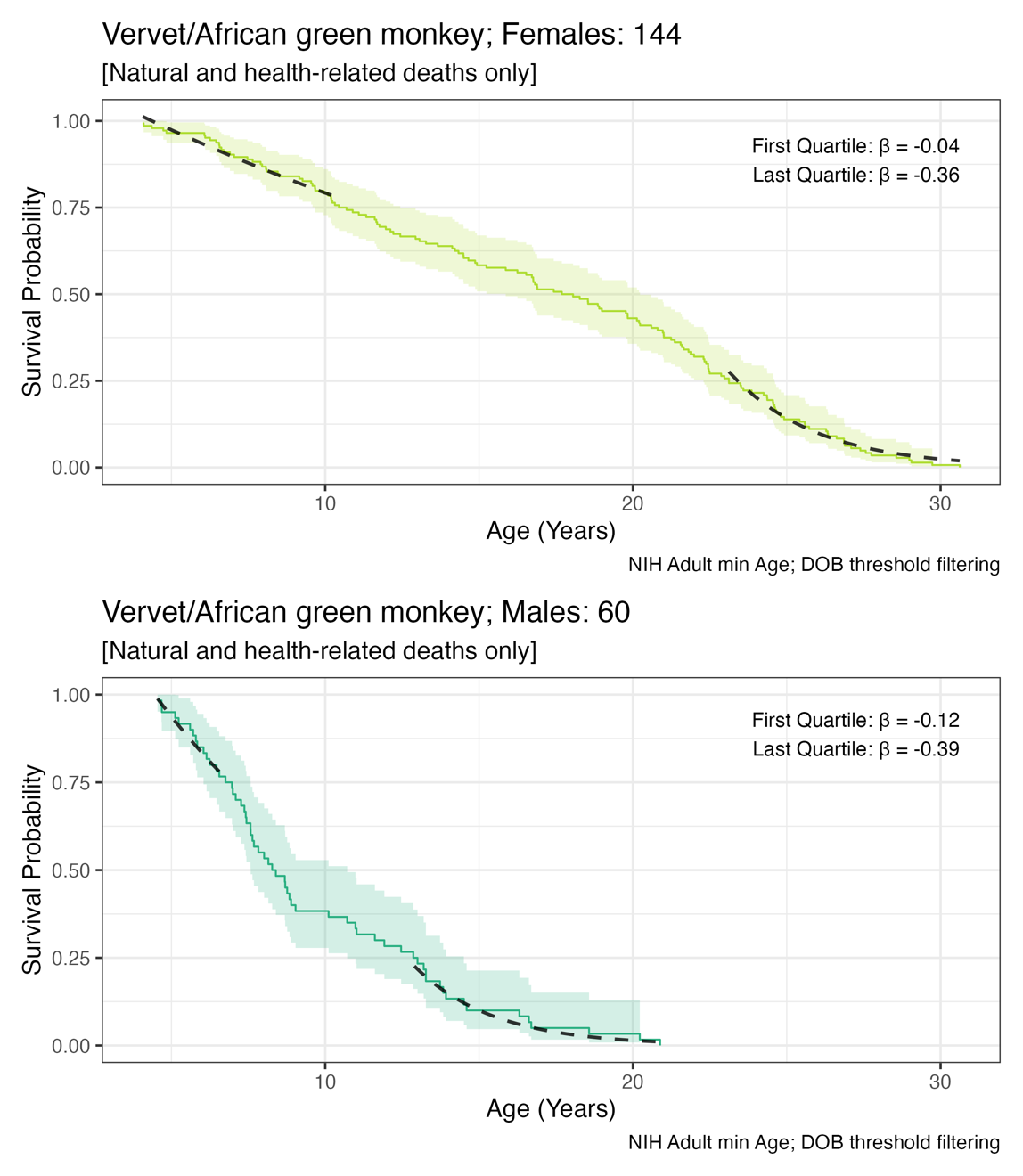

### **Figure S3:** Distribution and proportions of sex and censored data points for filtered data (n=32,616).

Deaths due to natural causes or health-related reasons are labeled as ‘natural.’ Deaths due to research sacrifice or colony management are labeled as ‘censored.’ Multiple species had very high proportions of censored events.

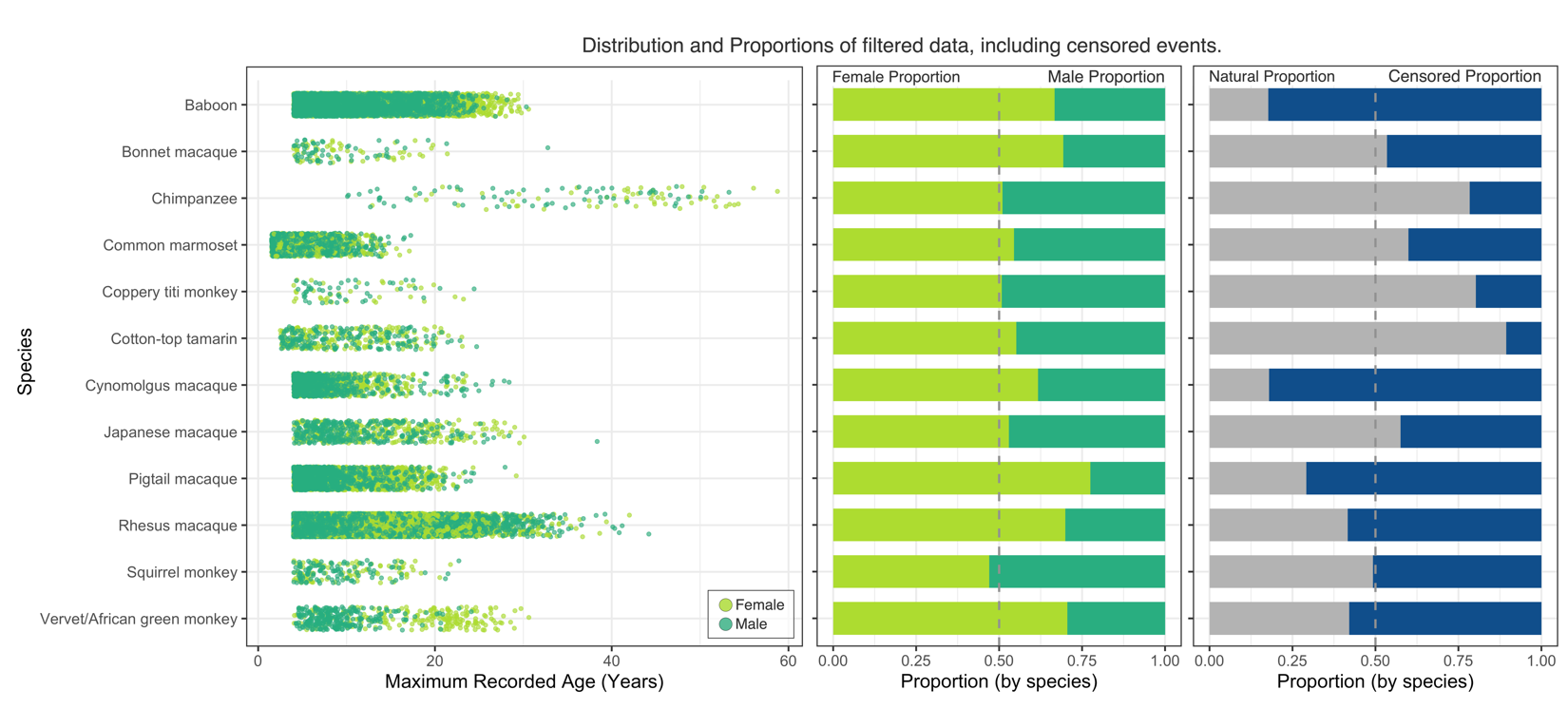

### **Figure S4:** Detailed Kaplan-Meier survivor curves for 12 species, by sex.

Here, we provide the Kaplan-Meier estimates of the survivorship function for all data passing Stage Two filtering. Presented are survival curves where deaths from research sacrifice and colony management are included as censored events, and for comparison, survival curves for the same data but limited to natural deaths and health-related deaths only. Importantly, censoring was informative of sex, so sex-specific comparisons were not computed for these data.

#### **Figure S4-A** Baboons

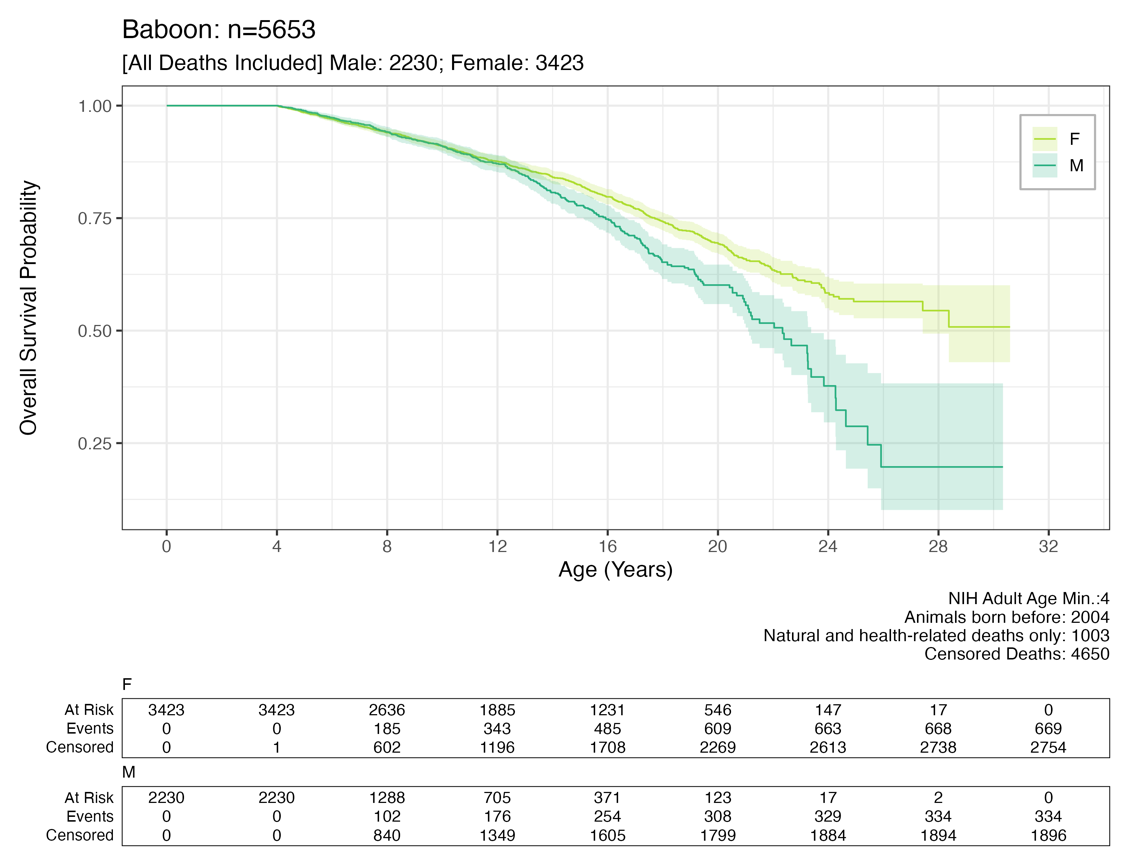

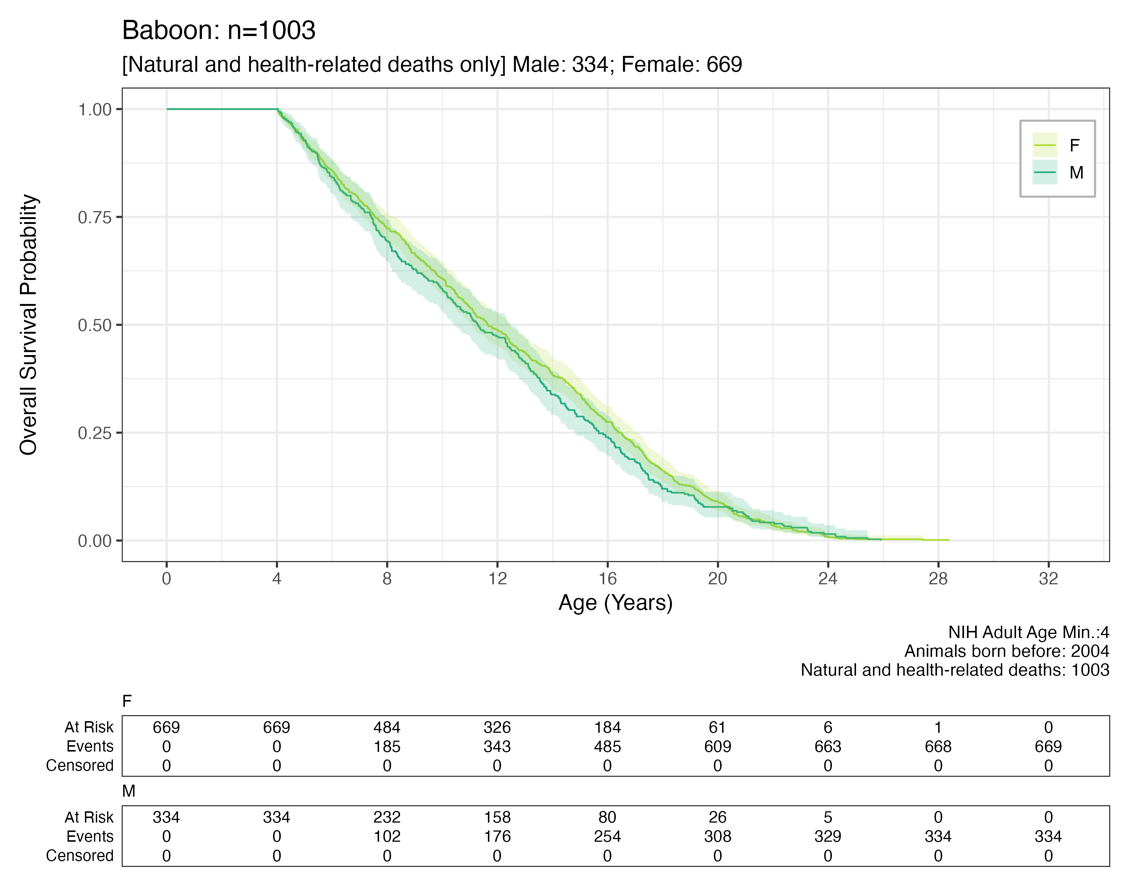

#### **Figure S4-B** Bonnet macaques

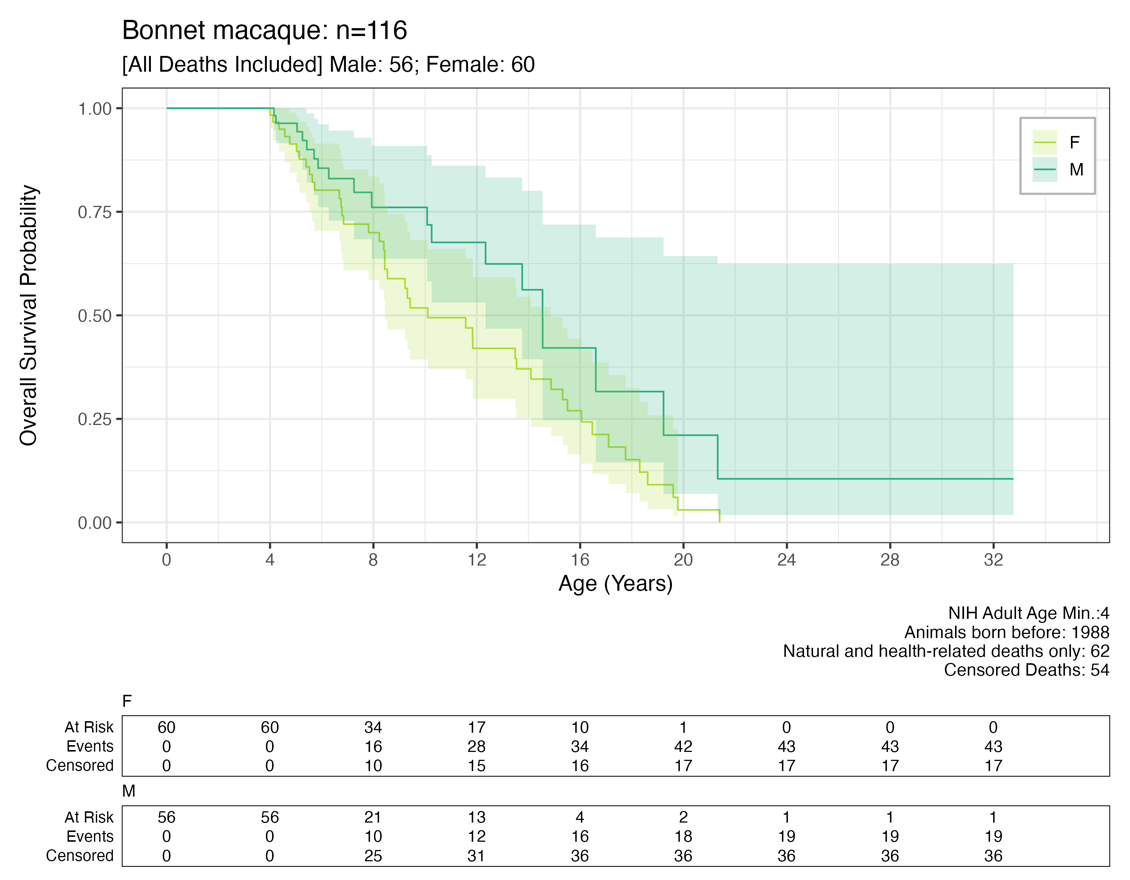

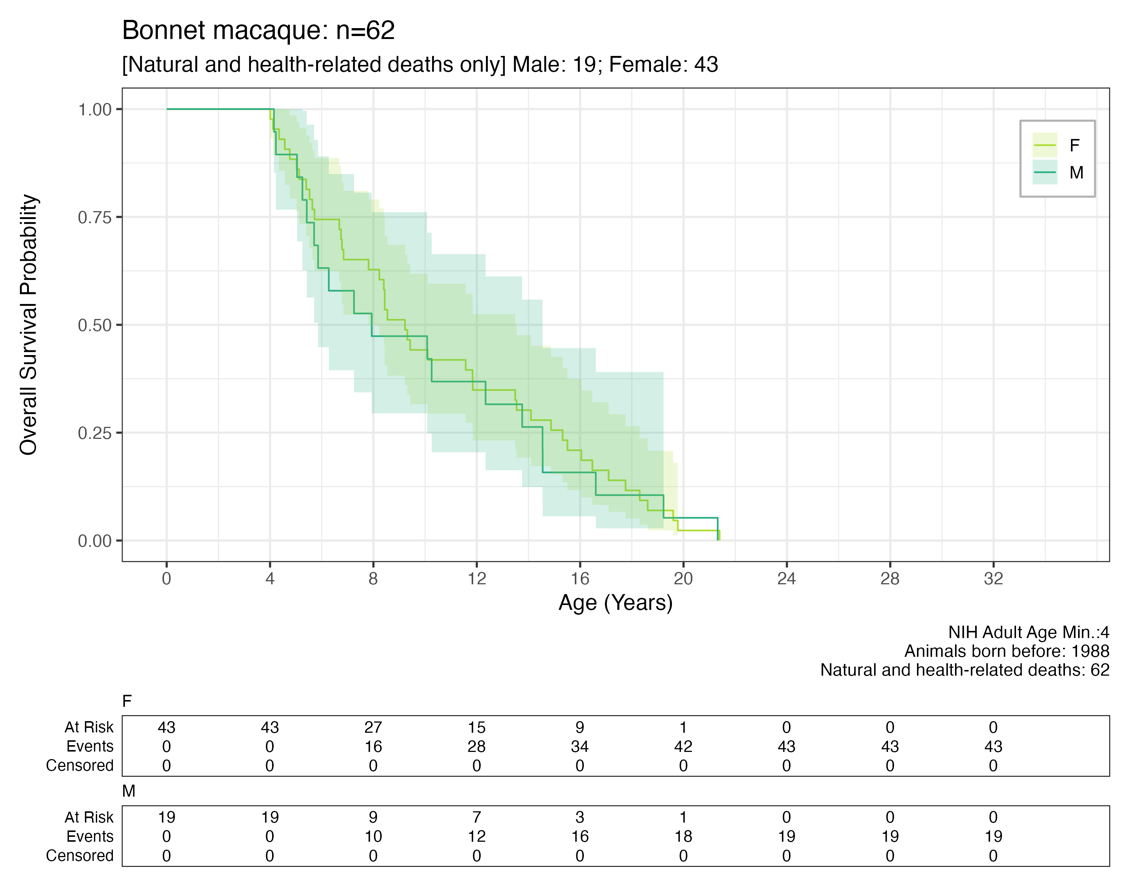

#### **Figure S4-C** Chimpanzees

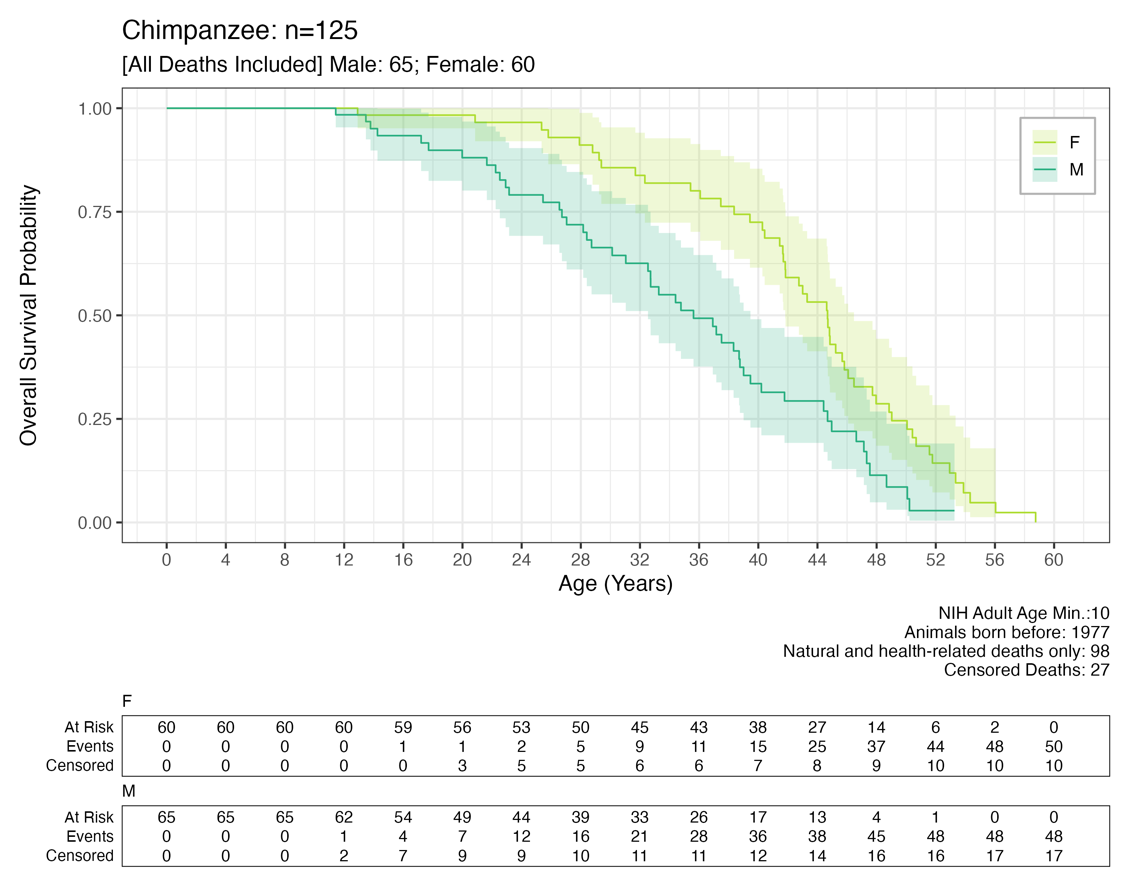

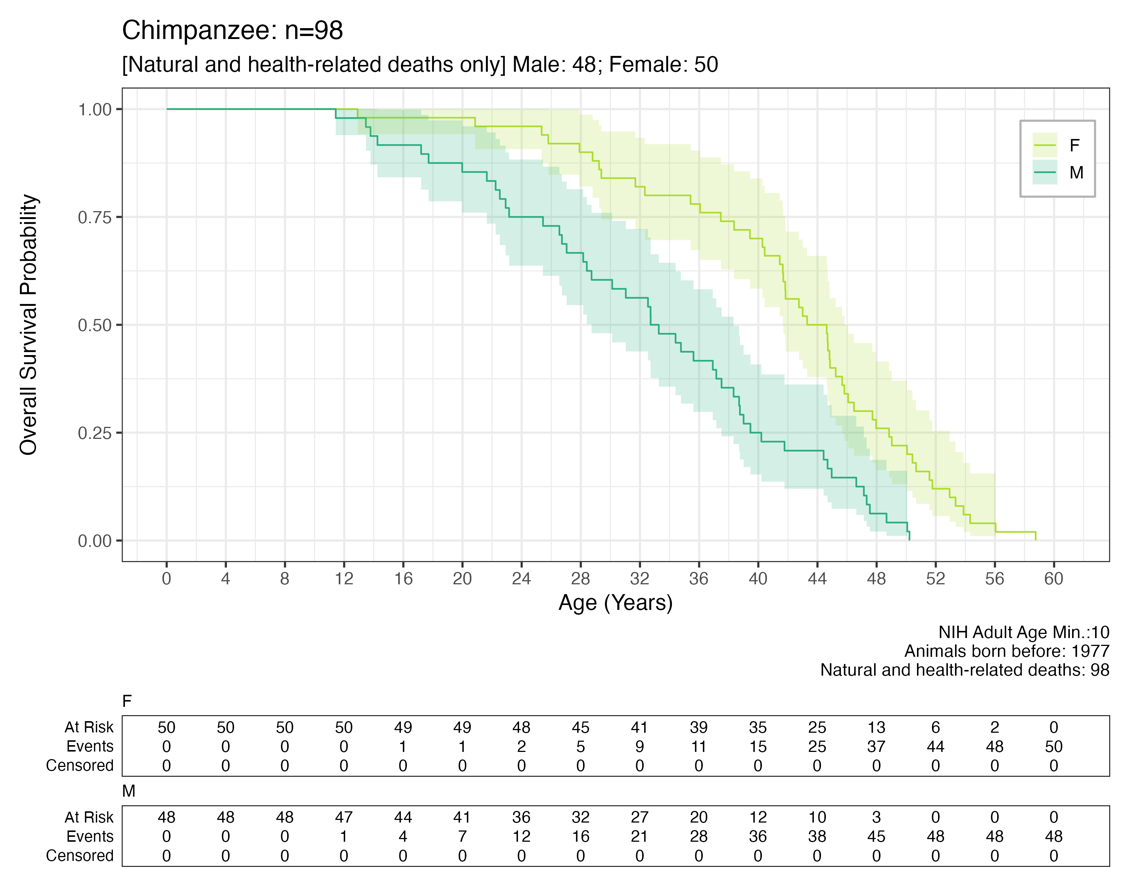

#### **Figure S4-D** Common marmoset

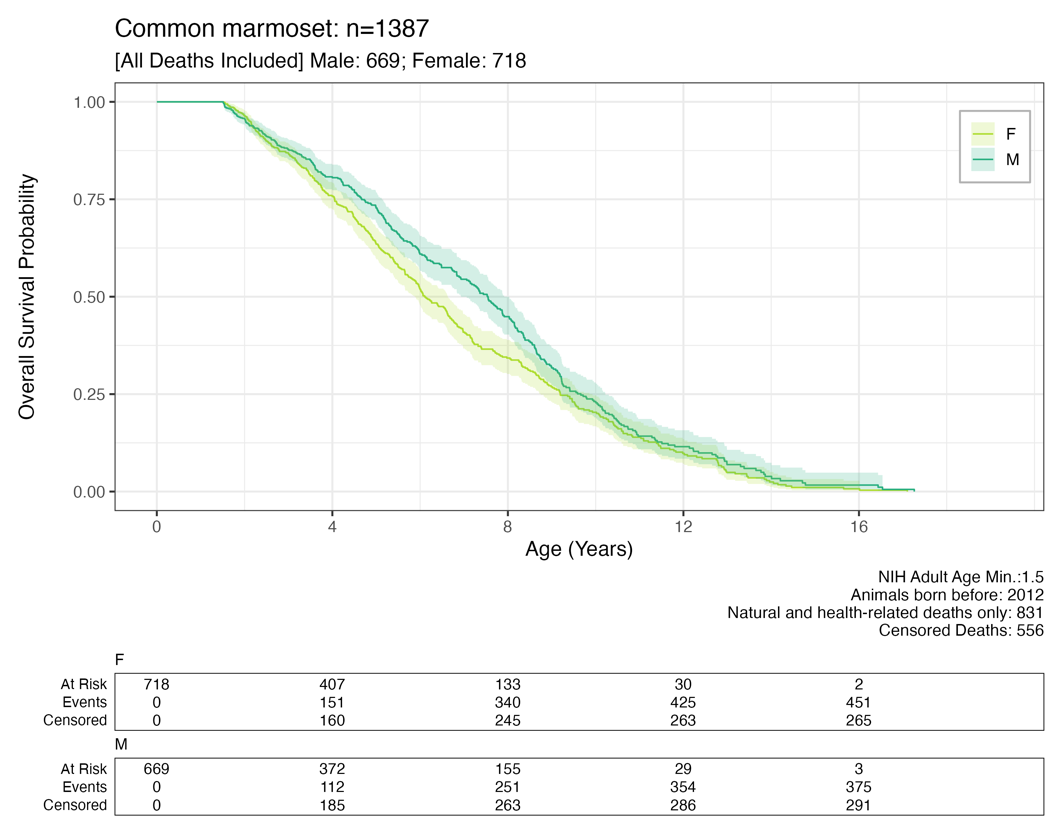

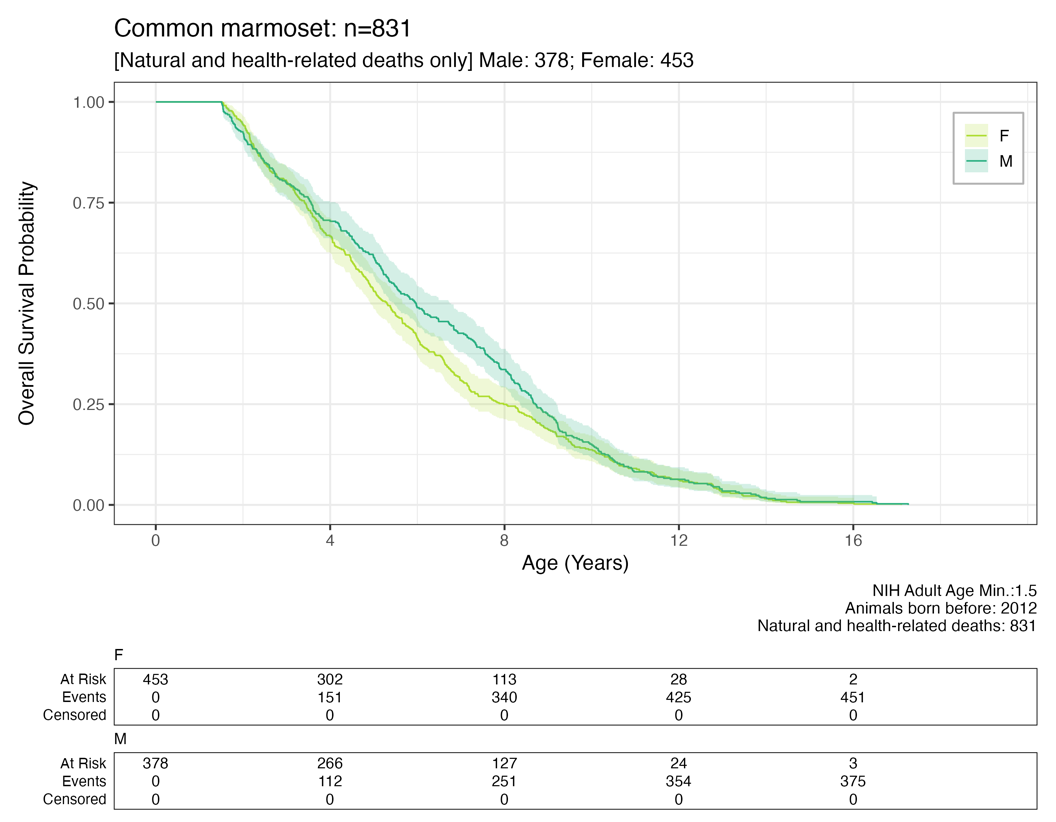

#### **Figure S4-E** Coppery titi monkeys

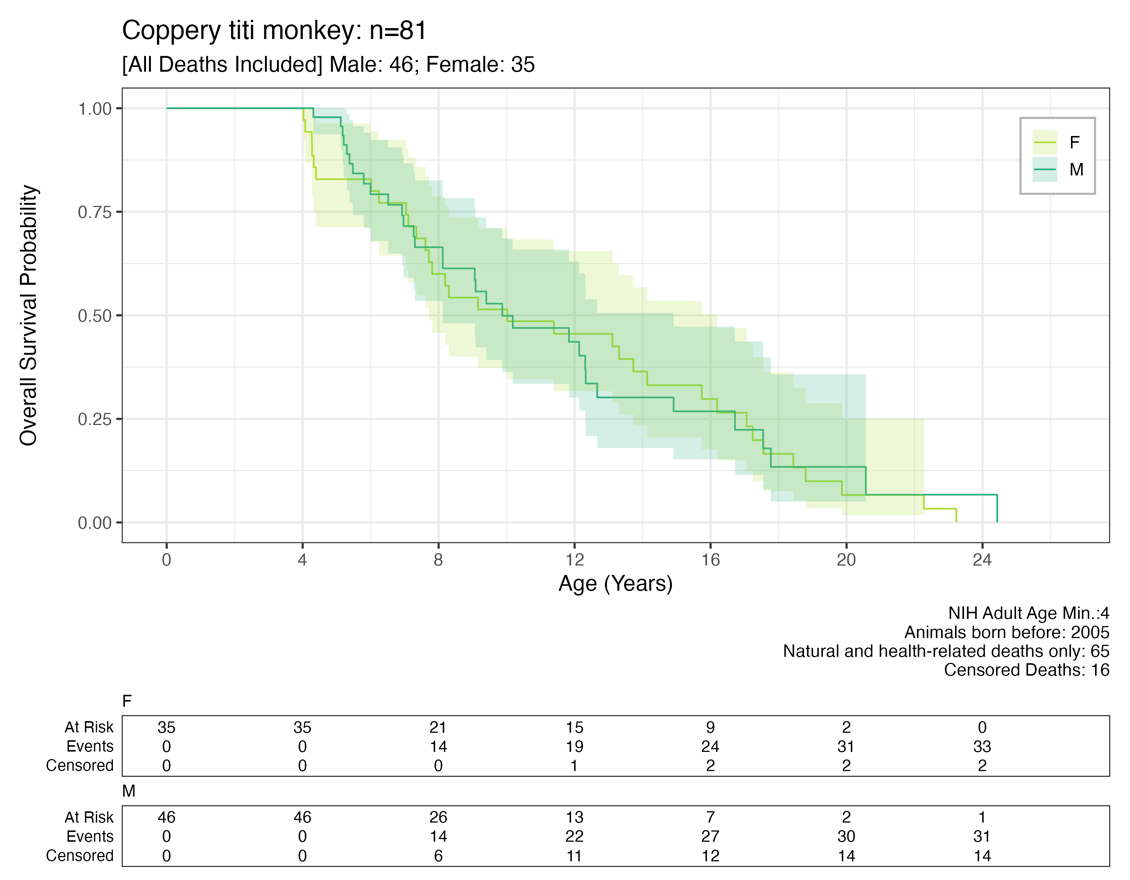

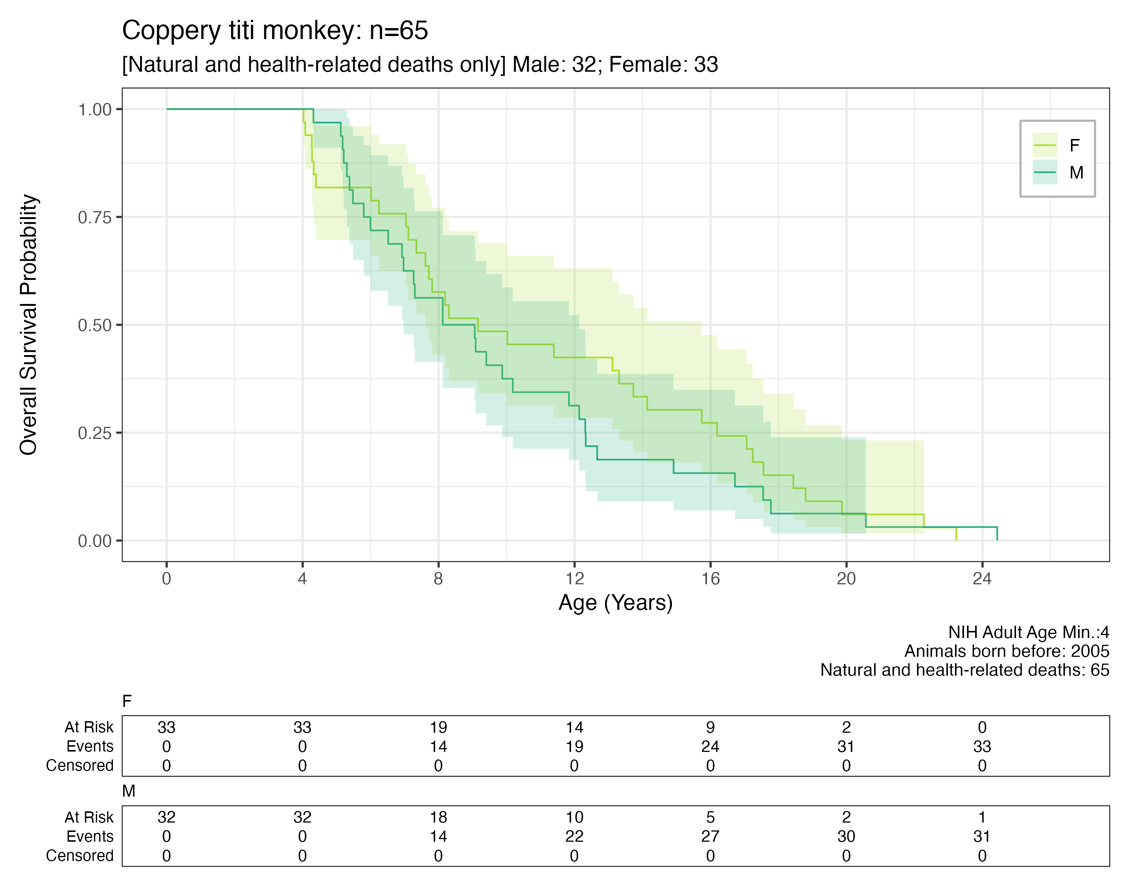

#### **Figure S4-F** Cotton-top tamarins

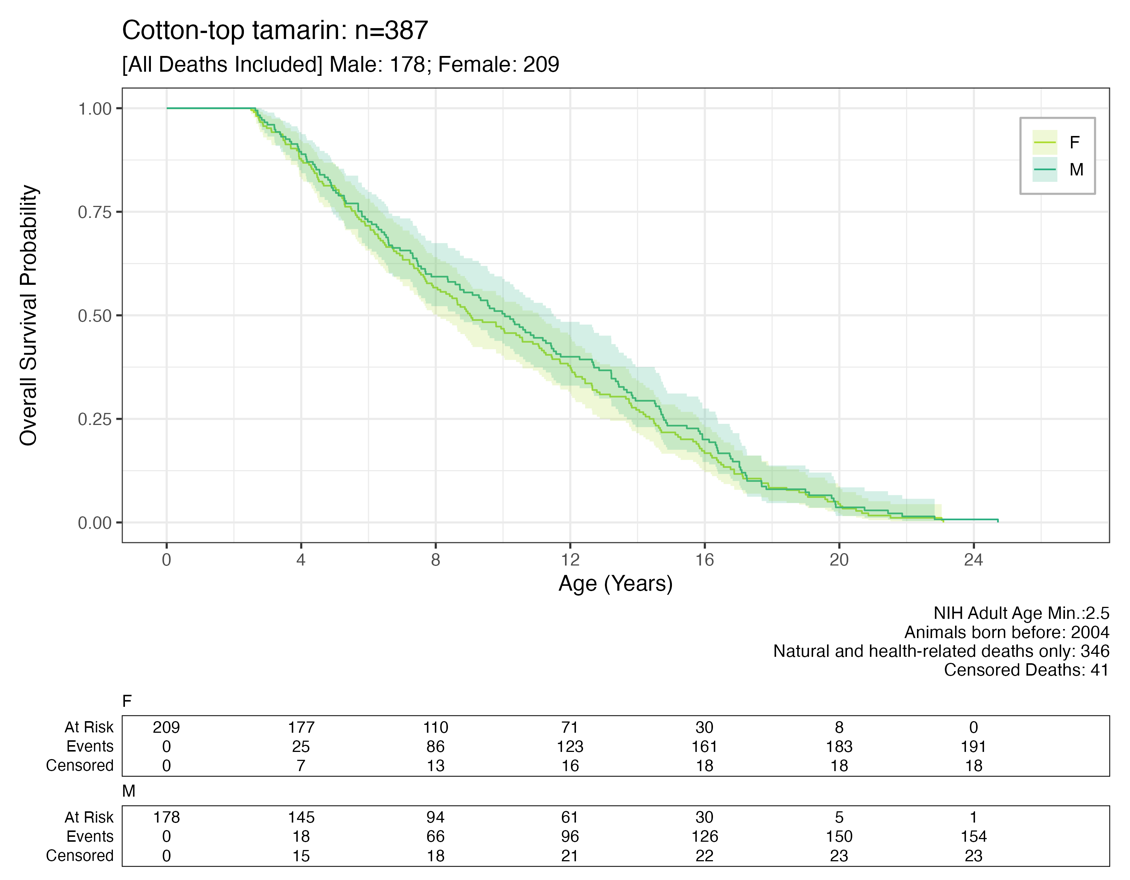

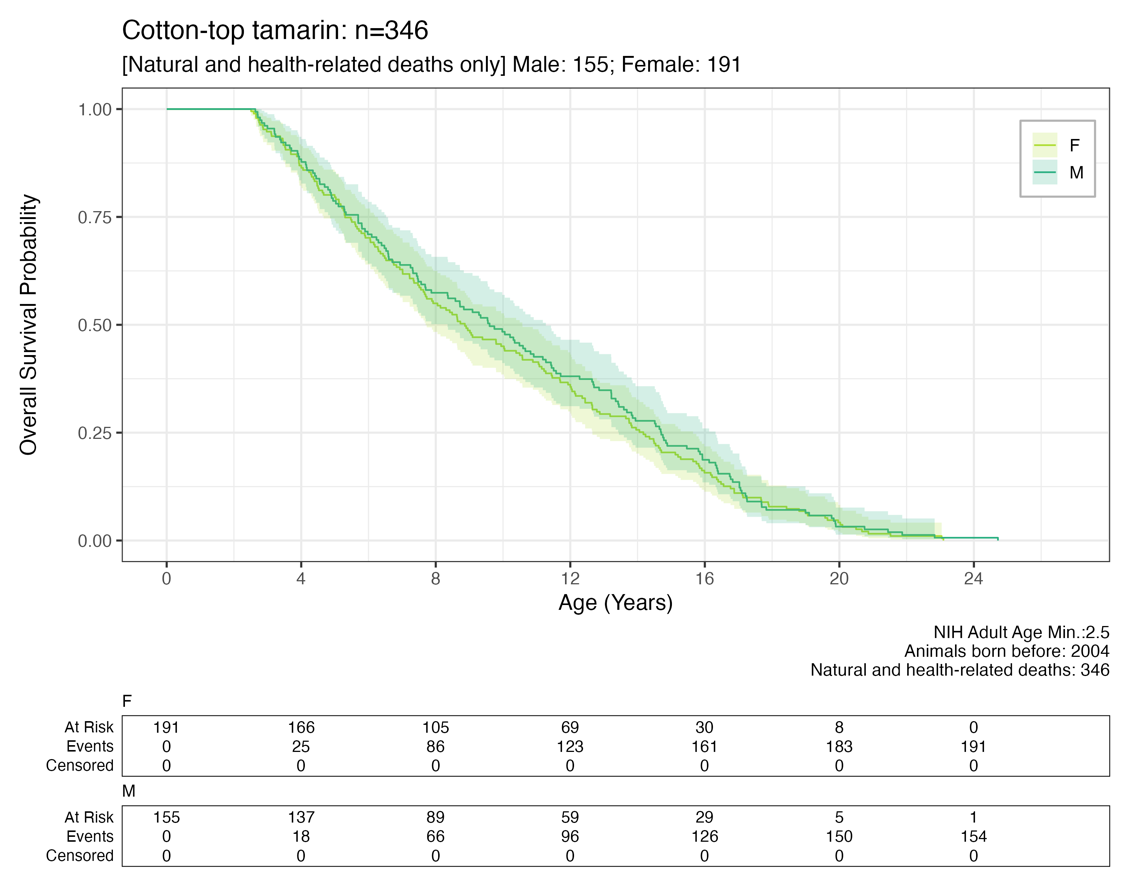

#### **Figure S4-G** Cynomolgus macaques

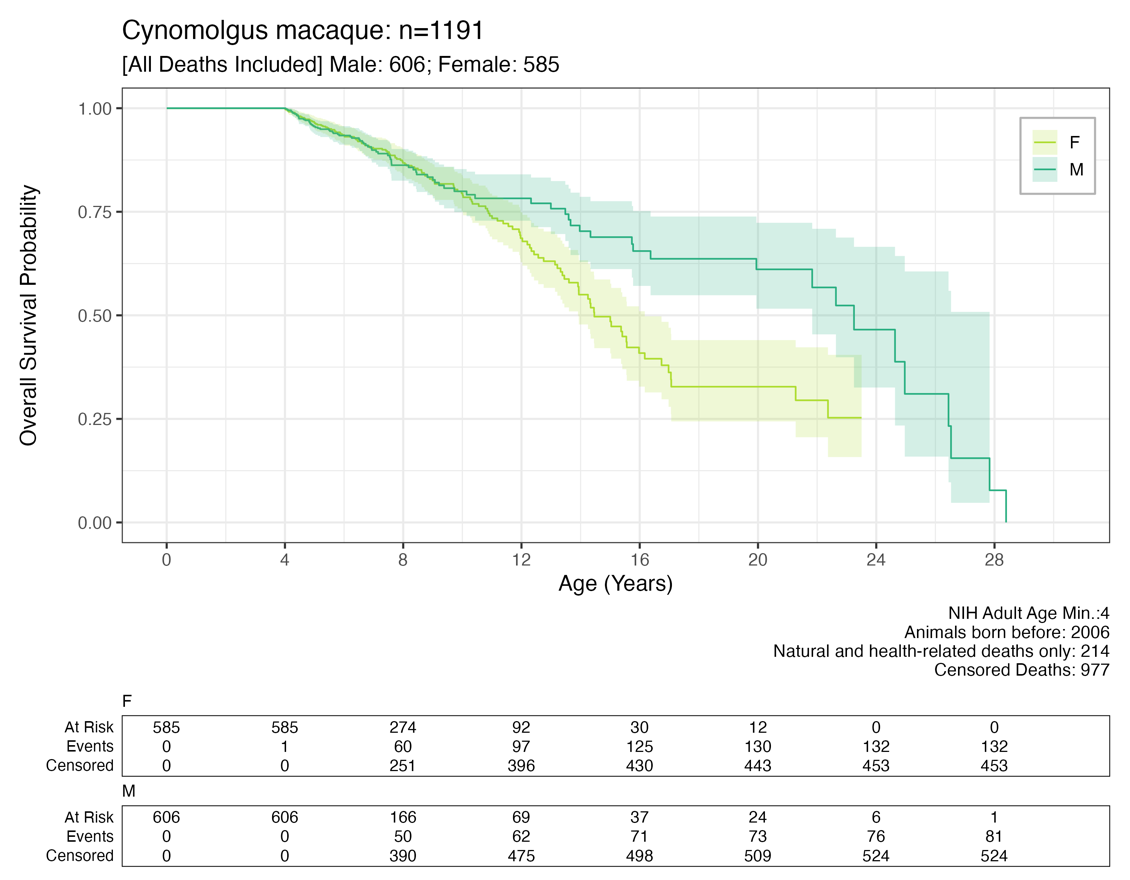

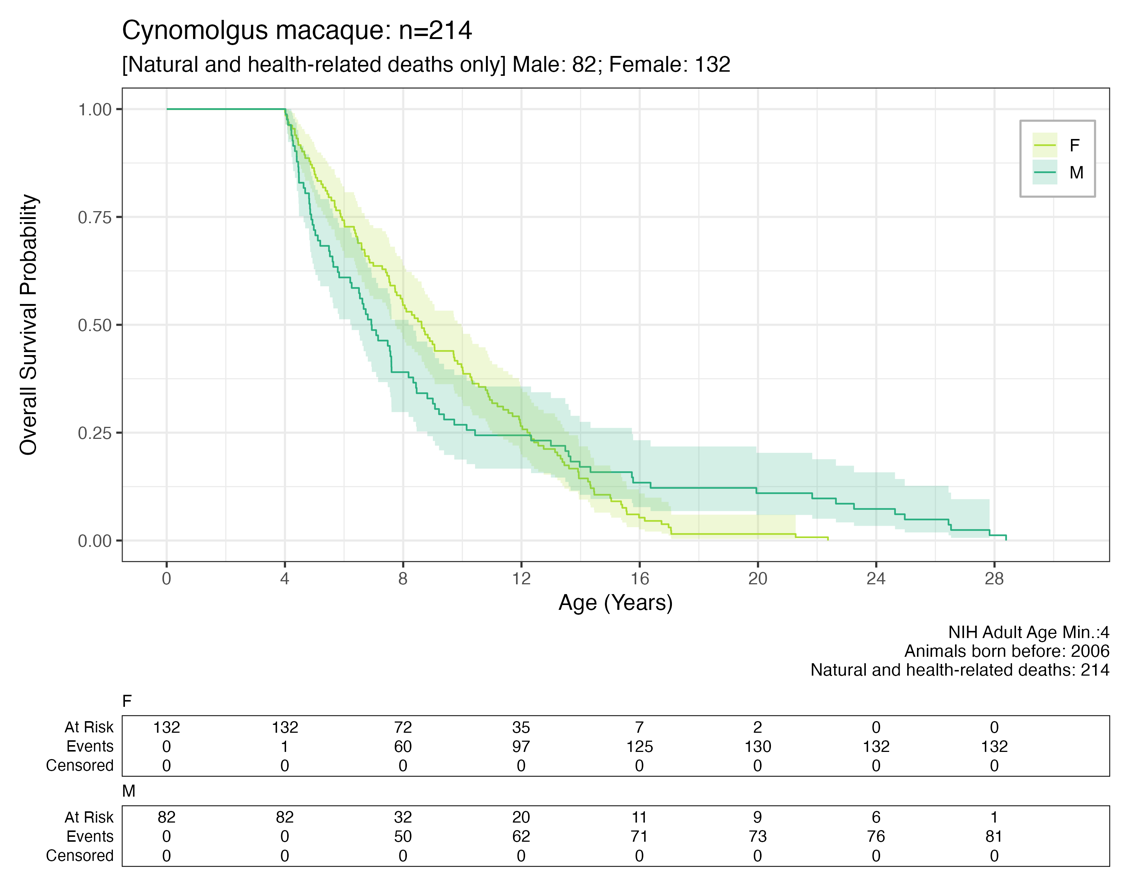

#### **Figure S4-H** Japanese macaques

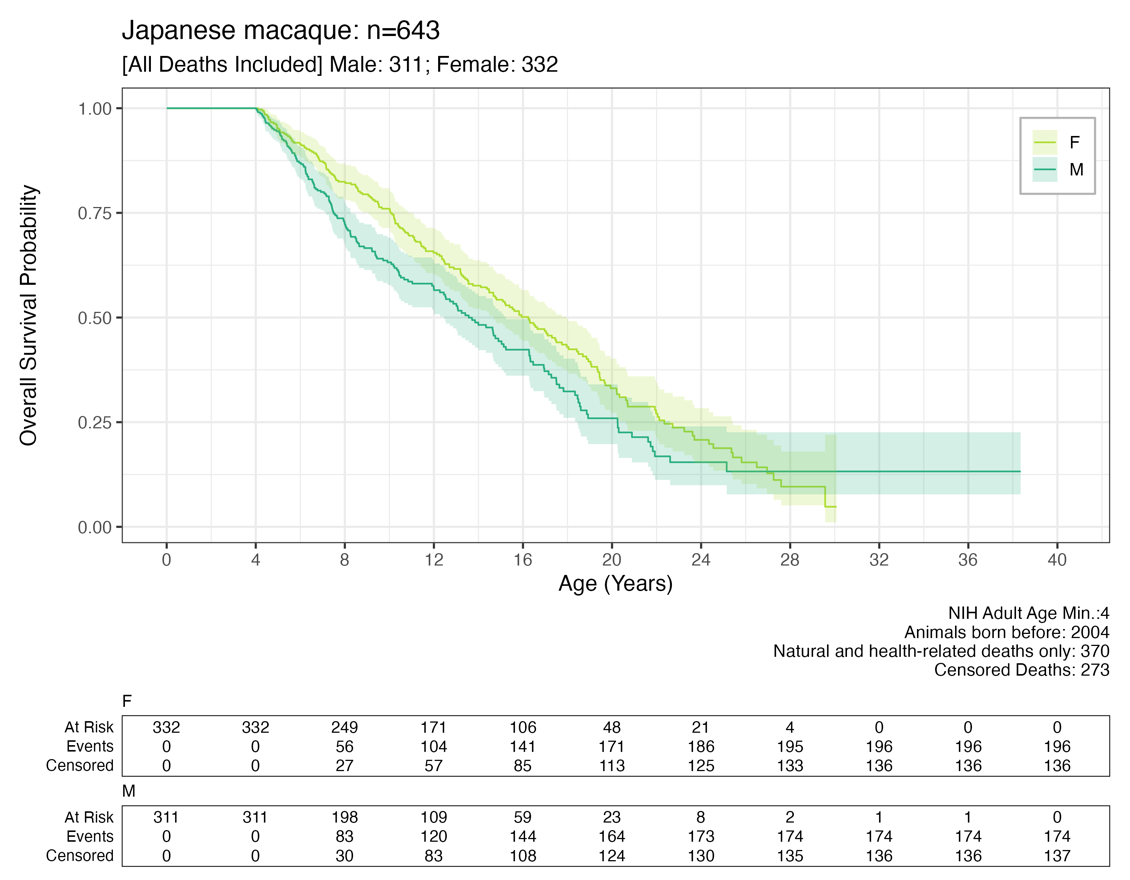

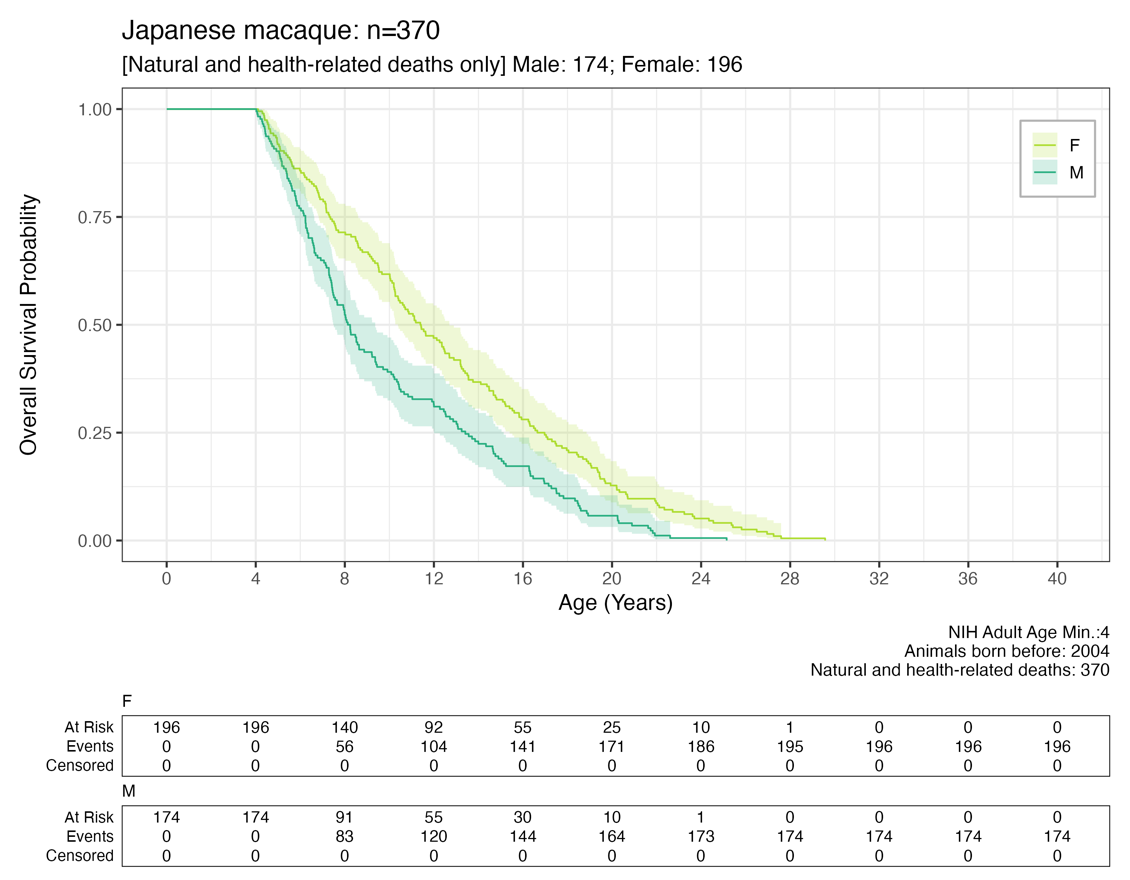

#### **Figure S4-I** Pigtail macaques

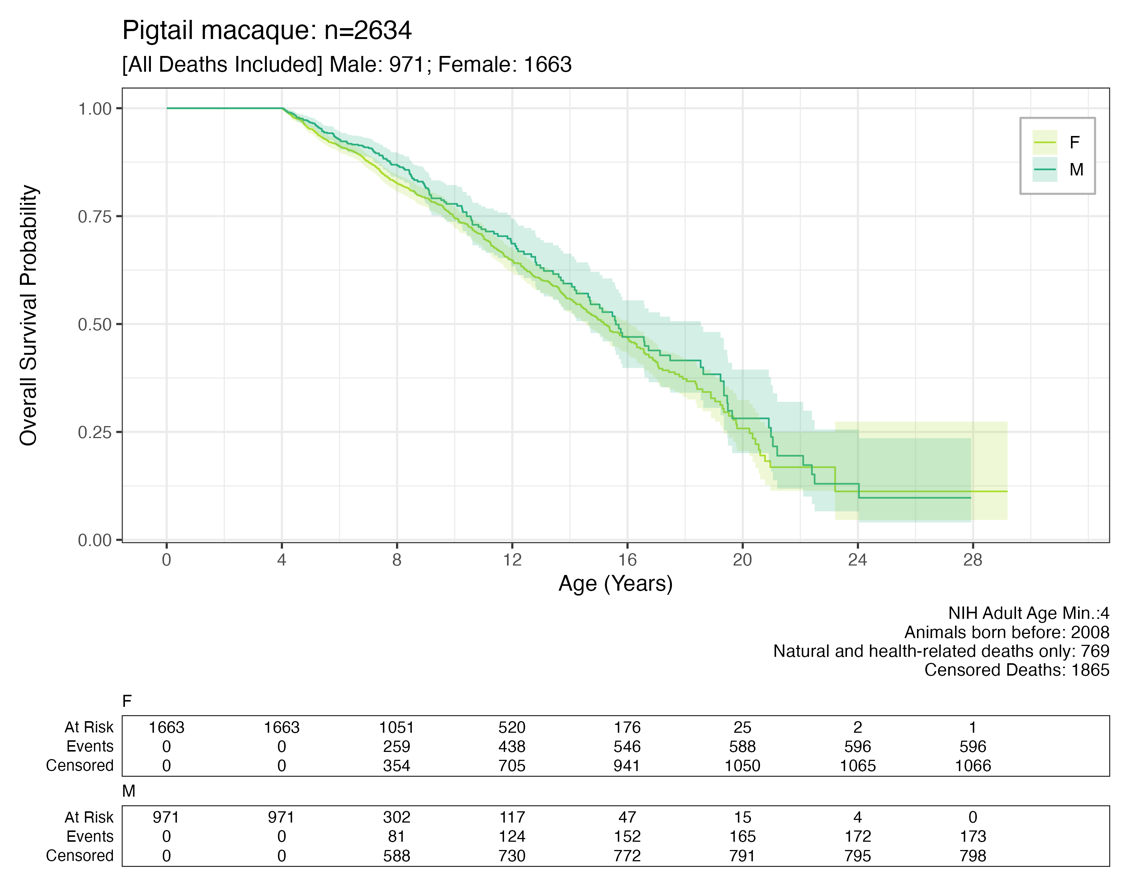

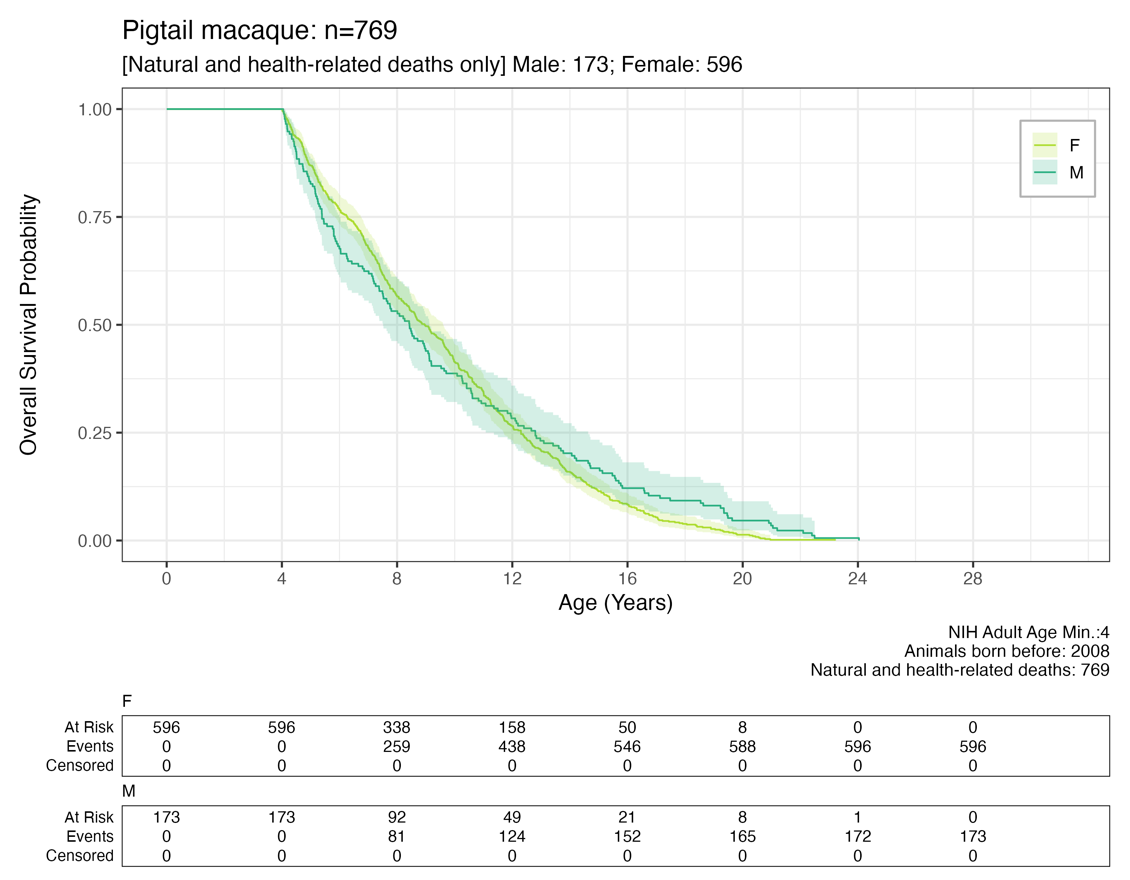

#### **Figure S4-J** Rhesus macaques

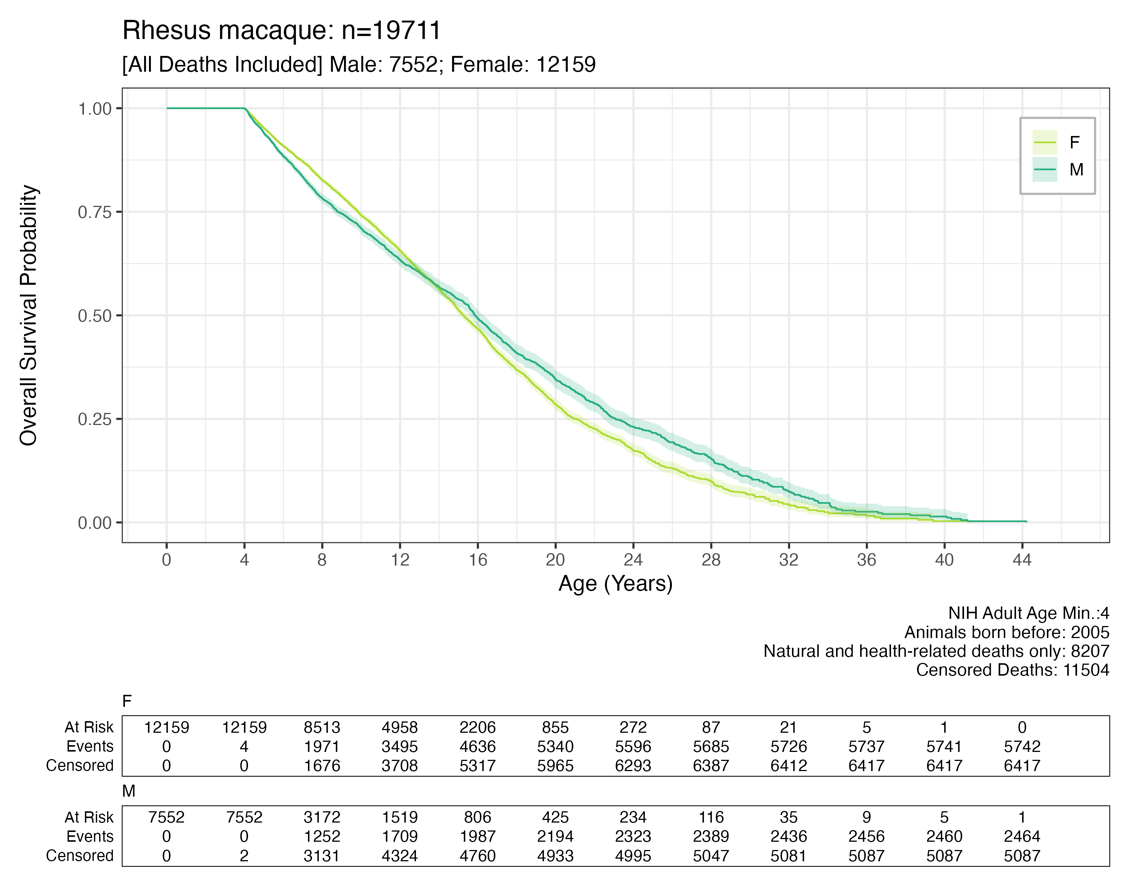

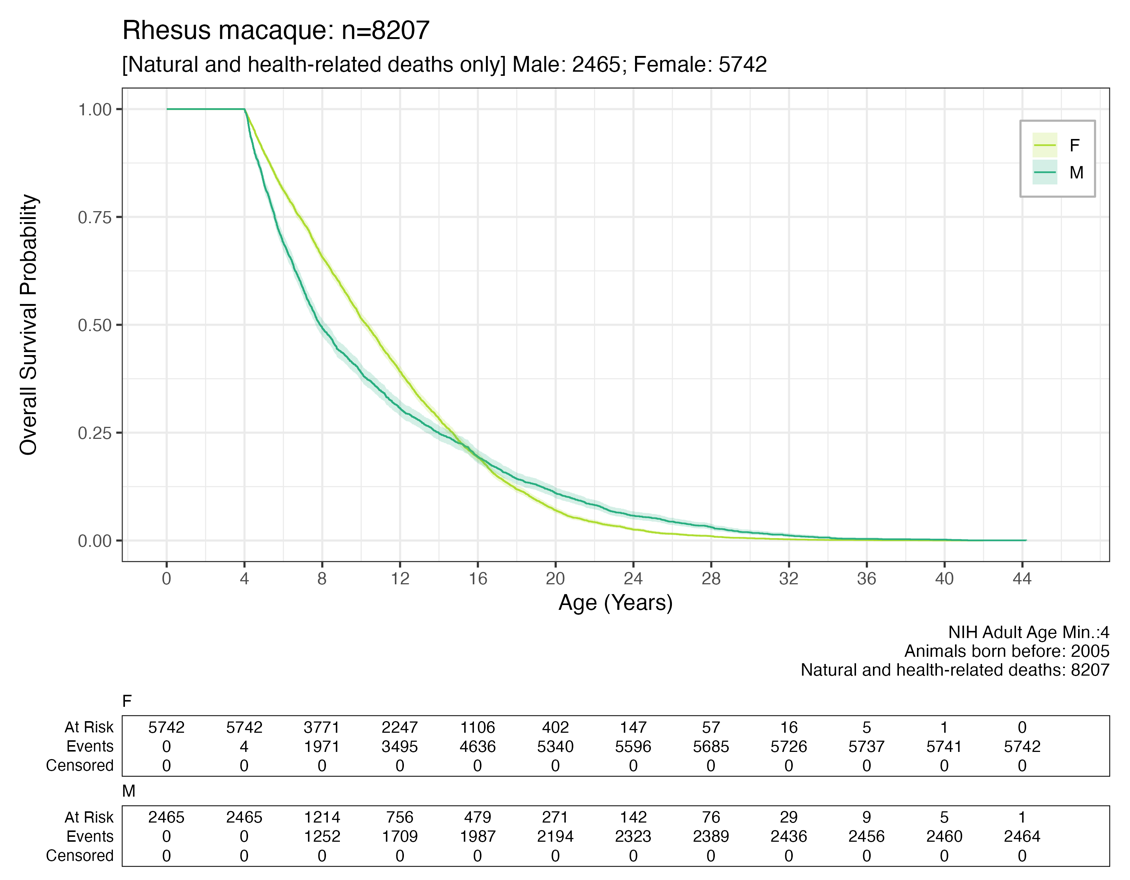

#### **Figure S4-K** Squirrel monkeys

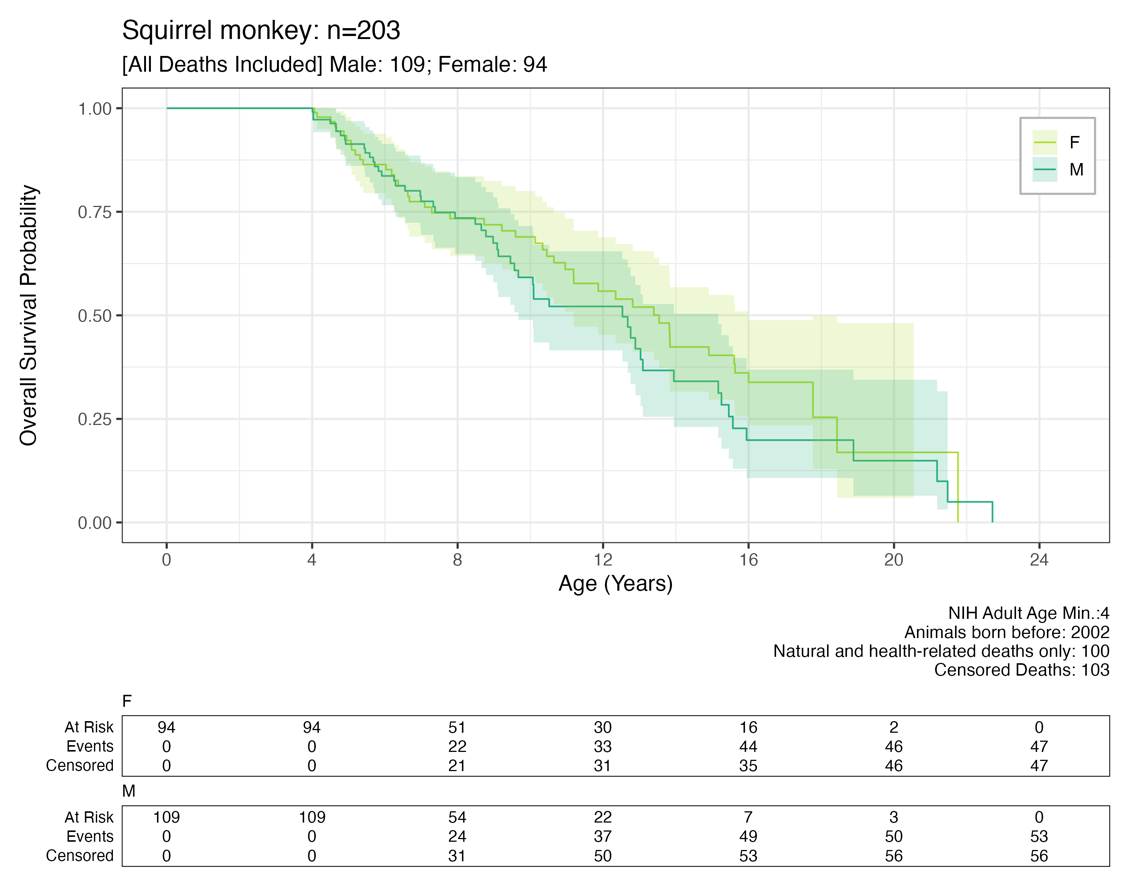

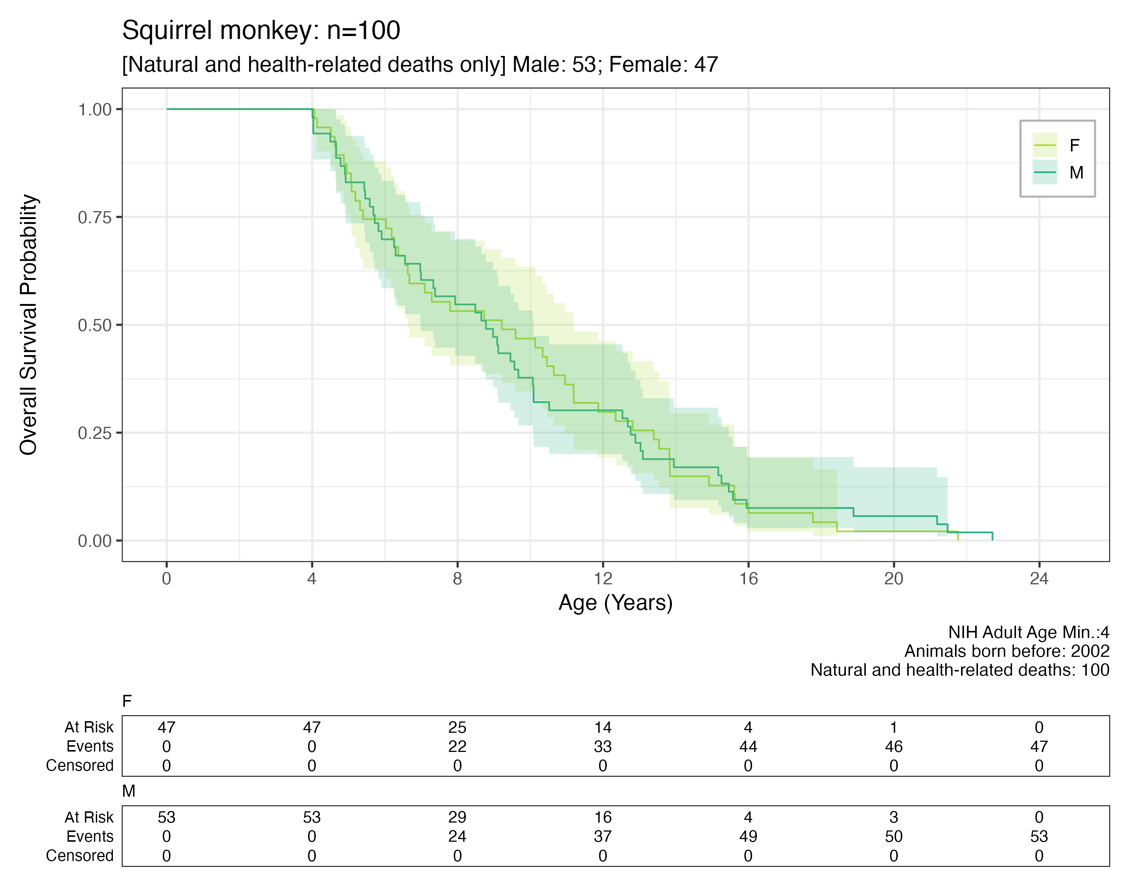

#### **Figure S4-L** Vervets/African green monkeys

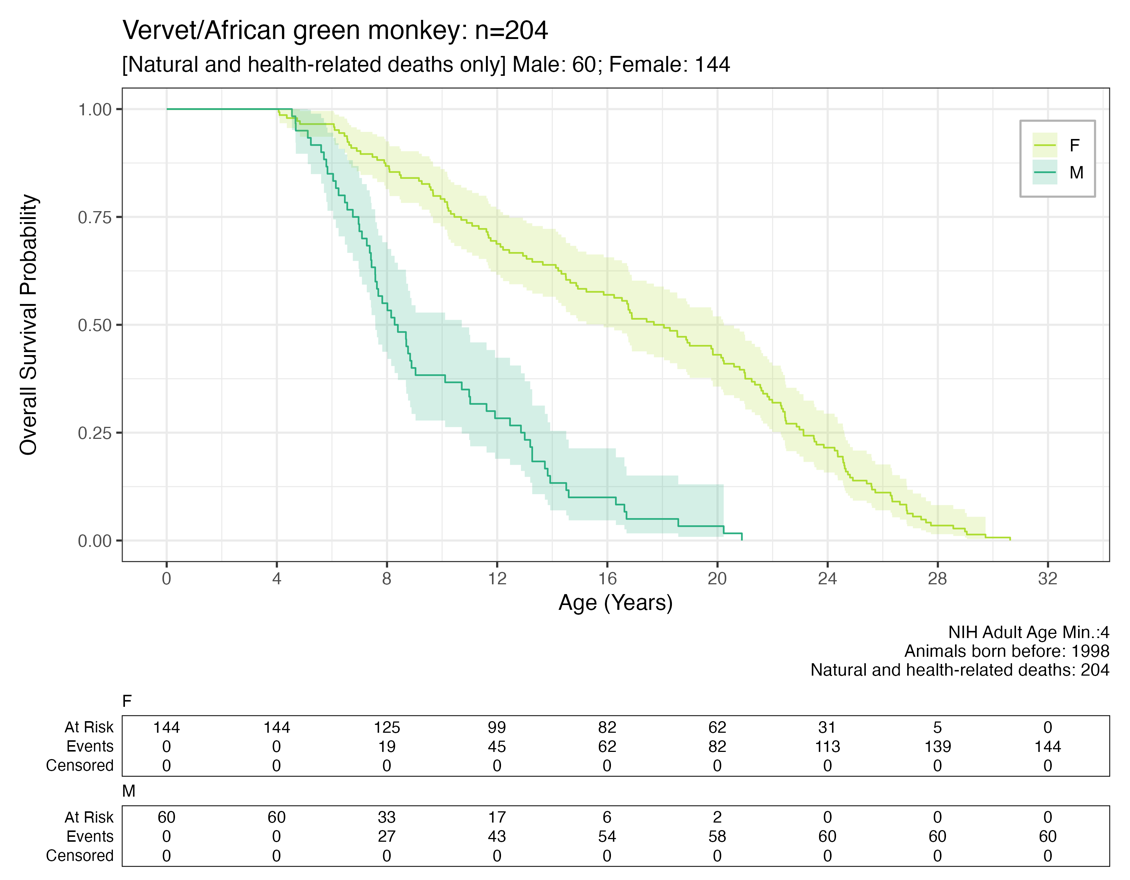

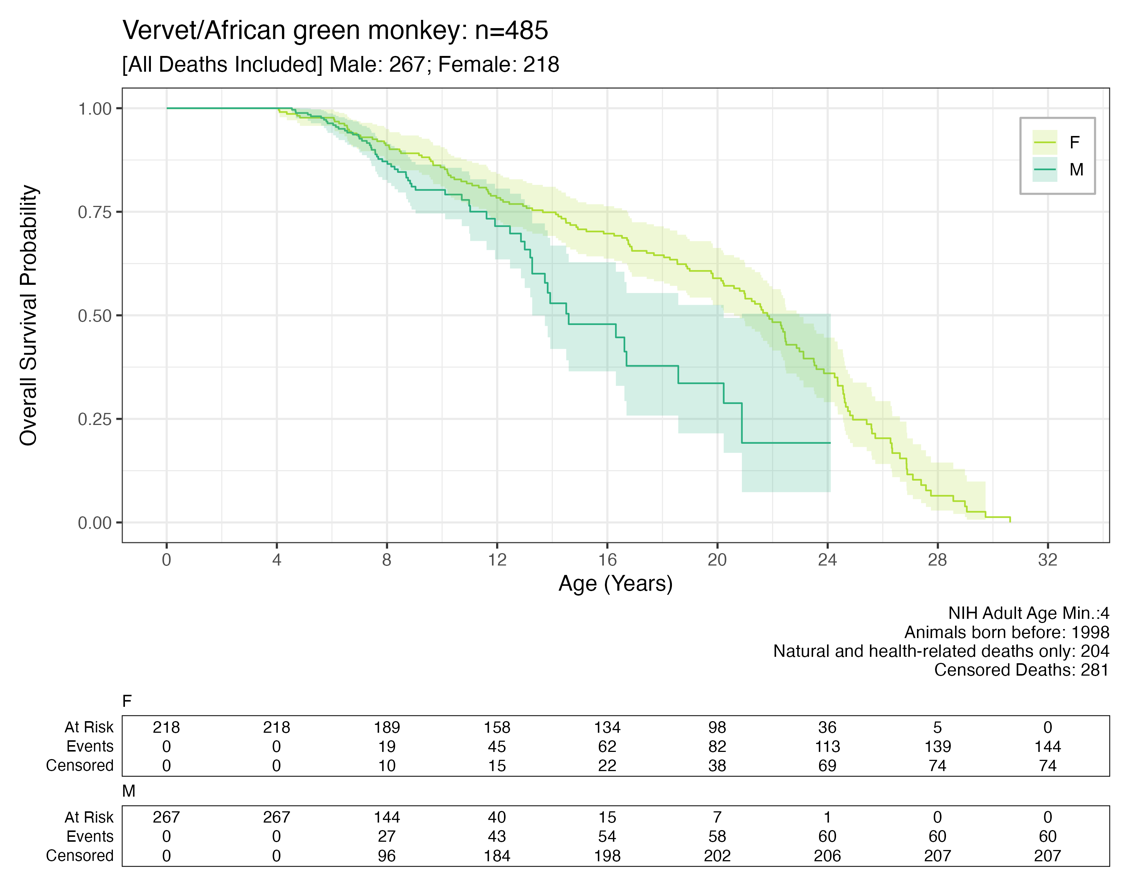

### **Supplementary Code 1.** Custom Code for fitting exponential curves to survival data.

**This code is also available via MIDAS (see Data Availability Statement).**

```
#R 4.1.2
#Comparative lifespan and healthspan of nonhuman primate species common to biomedical research
#(update for distribution: July 2024)

########################################################
## ABOUT DATA FOR THIS ANALYSIS                       ##
## -Natural deaths or health-related deaths only      ##
## -Minimum Age threshold = NIH adult age for species ##
## -Animals born after DOB threshold have been removed##
##  (to prevent skewing of data towards earlier deaths##
##   since alive animals are not included).           ##
## -Male (M) or Female (F)                            ##
## -No duplicated animals                             ##
## -12 species                                        ##
########################################################

###############################################
## SET SEED                                  ##
###############################################

set.seed(seed=49999)

##############################################
## LOAD LIBRARIES                           ##
##############################################

library(data.table) #for fread
library(ggplot2)    #for plot generation
library(patchwork)  #for combining plots
library(ggsurvfit)  #survival curve data generation


##############################################
## Open Full Dataset                        ##
## Includes censored events (to be removed) ##
##############################################
data_in <-fread("NHP_2024_postQCFilters_32616.tsv", header=T, sep="\t", stringsAsFactors = F)
nrow(data_in)
  #[1] 32616


##############################################
## LIMIT TO NATURAL DEATHS ONLY             ##
## censored coded as 0;                     ##
## non-censored (an event) coded as 1.      ##
##############################################

data_nat <-subset(data_in, censor_status==1)
nrow(data_nat)
  #[1] 12269

table(data_nat$censor_LABEL)
  #Natural or Health-Related Euthanasia
  #12269

table(data_nat$Species_name_common,data_nat$SEX_M_F)
       # F    M
       #  Baboon                       669  334
       #  Bonnet macaque                43   19
       #  Chimpanzee                    50   48
       #  Common marmoset              453  378
       #  Coppery titi monkey           33   32
       #  Cotton-top tamarin           191  155
       #  Cynomolgus macaque           132   82
       #  Japanese macaque             196  174
       #  Pigtail macaque              596  173
       #  Rhesus macaque              5742 2465
       #  Squirrel monkey               47   53
       #  Vervet/African green monkey  144   60

#list of species#
species_list_common <-as.vector(unique(data_nat$Species_name_common))


##########################################################
##                                                      ##
## FUNCTION TO FIT EXPONENTIAL CURVES TO FIRST AND LAST ##
## QUARTILES of SURVIVAL DATA;                          ##
## OUTPUT: PLOT AND TABLE OF EXP PARAMETERS.            ##
##                                                      ##
##########################################################

analyze_sex <- function(mydat, sex_value, sex_label, myMIN_AGE_CUTOFF,species_name,sex_color) {
  mydat_sex <- subset(mydat, SEX_M_F == sex_value)
  my_count <- nrow(mydat_sex)

  ##get 25th and 75th quartiles of max age (years): REMINDER: THESE DATA ARE ABSENT OF CENSORED EVENTS##
  ##These will be used to subset survival data prior to fitting exponential curve.
  my_25th <- as.numeric(quantile(mydat_sex$AGE_YEARS, probs = c(0.25)))
  my_75th <- as.numeric(quantile(mydat_sex$AGE_YEARS, probs = c(0.75)))

  ##generate dataframe of survival curve info##
  ##(note survival is using data for only a single Sex, as designated in function parameters)##
  mysurv <- survfit2(Surv(AGE_YEARS, censor_status) ~ SEX_M_F, data = mydat_sex, model = TRUE) %>%
    ggsurvfit()
    mysurv_dat <- as.data.frame(cbind(mysurv$data$time, mysurv$data$estimate))
    colnames(mysurv_dat) <- c("AGE_YEARS", "survival_prob")

  ###limit to relevant ages for each quartile group##
  ###reminder: we want to fit exponential curves STARTING at adulthood.
  ###          Because we have pruned animals less than adulthood, the
  ###          survival probability will=1 for ages less than adulthood (flat line).
  mysurv_dat_adults_25th <- subset(mysurv_dat, AGE_YEARS >= myMIN_AGE_CUTOFF & AGE_YEARS <= my_25th)
  mysurve_25_count <-nrow(mysurv_dat_adults_25th)
  mysurv_dat_adults_75th <- subset(mysurv_dat, AGE_YEARS >= my_75th)
  mysurve_75_count <-nrow(mysurv_dat_adults_75th)

  ##convert to vectors (for ease of computing)##
  X_myages_25 <- as.vector(mysurv_dat_adults_25th$AGE_YEARS)
  Y_mysurv_25 <- as.vector(mysurv_dat_adults_25th$survival_prob)

  X_myages_75 <- as.vector(mysurv_dat_adults_75th$AGE_YEARS)
  Y_mysurv_75 <- as.vector(mysurv_dat_adults_75th$survival_prob)

  ##fit exponential models and then save coefficients to a dataframe##
  ##--FIRST QUARTILE--##
  exp_model_25 <- nls(Y_mysurv_25 ~ a * exp(b * X_myages_25), start = list(a = 1, b = -0.1))
  print(summary(exp_model_25))
  mycoefficients_25 <- as.data.frame(t(as.data.frame(coef(exp_model_25))))
     colnames(mycoefficients_25) <-c("a_parameter","b_parameter")
     mycoefficients_25$sex <- sex_value
     mycoefficients_25$pct <- "25th"
  mycoeff_stderr_25 <-as.data.frame(t(summary(exp_model_25)$parameters[,"Std. Error"]))
     colnames(mycoeff_stderr_25) <-c("a_stdError","b_stdError")
  mycoefficients_25_tomerge <-cbind(mycoefficients_25,mycoeff_stderr_25)
  mycoefficients_25_tomerge$ResidStdErr <- as.numeric(summary(exp_model_25)$sigma) ## note --residual standard error (average goodness of fit)
  mycoefficients_25_tomerge$n_animals_quartile <-mysurve_25_count

  ##--LAST QUARTILE--##
  exp_model_75 <- nls(Y_mysurv_75 ~ a * exp(b * X_myages_75), start = list(a = 1, b = -0.1))
  print(summary(exp_model_75))
  mycoefficients_75 <- as.data.frame(t(as.data.frame(coef(exp_model_75))))
    colnames(mycoefficients_75) <-c("a_parameter","b_parameter")
    mycoefficients_75$sex <- sex_value
    mycoefficients_75$pct <- "75th"
  mycoeff_stderr_75 <-as.data.frame(t(summary(exp_model_75)$parameters[,"Std. Error"]))
     colnames(mycoeff_stderr_75) <-c("a_stdError","b_stdError")
  mycoefficients_75_tomerge <-cbind(mycoefficients_75,mycoeff_stderr_75)
  mycoefficients_75_tomerge$ResidStdErr <- as.numeric(summary(exp_model_75)$sigma) ## note --residual standard error (average goodness of fit)
    mycoefficients_75_tomerge$n_animals_quartile <-mysurve_75_count


  ##create annotation text that will be added to the plots##
  beta_25 <-as.vector(mycoefficients_25$b)
  beta_75 <-as.vector(mycoefficients_75$b)

  ##clean up annotation text formatting##
  annotation_txt_exp <-paste0("First Quartile: \u03b2 = ",round(beta_25,2),"\n",
                              "Last Quartile: \u03b2 = ",round(beta_75,2))


  ##combine exponential model info to a single dataframe##
  mycoefficients_out <- rbind(mycoefficients_25_tomerge, mycoefficients_75_tomerge)
  mycoefficients_out$Species_name_common <-species_name
  mycoefficients_out$model <-"Exponential"
  mycoefficients_out$count_bySex <-my_count

  ##############################################################################
  ##BEGIN preparation of plotting exponential curves ontop of survival curves ##
  ##############################################################################

  ##This will visualize the fit of the exponential model to the data.         ##
  ##Create a sequence of x values for the line (based on our data min and max ages)
  xx_25 <- seq(min(X_myages_25), my_25th, 0.1) #For some species, the youngest animal might be slightly older (~month) than Adult age threshold (hence, using min age, to capture smallest value)
  xx_75 <- seq(my_75th, max(X_myages_75), 0.1)

  ##create predicted values based on x values these will get plotted on top of survival curves##
  mypredicted_25 <- predict(exp_model_25, newdata = data.frame(X_myages_25 = xx_25))
  mypredicted_df_25 <- as.data.frame(cbind(xx_25, mypredicted_25))

  mypredicted_75 <- predict(exp_model_75, newdata = data.frame(X_myages_75 = xx_75))
  mypredicted_df_75 <- as.data.frame(cbind(xx_75, mypredicted_75))


  ###Create a plot that shows the fitted curve overlap###
  fitted_plt <- survfit2(Surv(AGE_YEARS, censor_status)~SEX_M_F, data=mydat_sex, model=T) %>%
    ggsurvfit(color=sex_color) +
    add_confidence_interval(fill=sex_color) +
    geom_line(data=mypredicted_df_25,aes(x = xx_25, y = mypredicted_25), color = "black", alpha=0.8,linewidth=0.9,linetype="dashed") +  ## Adding exp fit for 1st quartile
    geom_line(data=mypredicted_df_75,aes(x = xx_75, y = mypredicted_75), color = "black", alpha=0.8,linewidth=0.9,linetype="dashed") +  ## Adding exp fit for last quartile
    annotate("text", x = max(mydat$AGE_YEARS), y = 0.95, label = annotation_txt_exp, vjust = "inward", hjust = "inward", size = 4) +
    xlim(min(mydat$AGE_YEARS),max(mydat$AGE_YEARS)) +
    labs(title=paste0(species_name,"; ", sex_label,": ",my_count),
         subtitle="[Natural and health-related deaths only]",
         x = "Age (Years)", y = "Survival Probability",
         caption="NIH Adult min Age; DOB threshold filtering") +
    theme_bw(base_size=14)

  return(list(my_coeff_res=mycoefficients_out,
              my_plt=fitted_plt,
              my_25th_exp_model=exp_model_75))

} #end function analyze_sex


#############################################################
##                                                         ##
## BEGIN FITTING EXPONENTIALS TO EACH SPECIES DATA, BY SEX ##
##                                                         ##
#############################################################

for (myspecies in species_list_common) {

  ##remove non alphanumeric characters from name (easier for saving files)
  species_for_filename <-gsub(" ","_",gsub("[/]","",myspecies))


  print(myspecies)
  ## Prep data and info needed to run exponential fit function ##
  mydat <-subset(data_nat, Species_name_common==myspecies)
  nrow(mydat)

  myMIN_AGE_CUTOFF <-head(mydat$MIN_AGE_CUTOFF,1)
  my_species_count <-nrow(mydat)

  female_analysis <- analyze_sex(mydat, "F","Females", myMIN_AGE_CUTOFF,myspecies,"#addc30")
  male_analysis   <- analyze_sex(mydat, "M", "Males",myMIN_AGE_CUTOFF,myspecies,"#28ae80")

  ##output a table of the exponential coefficients##
  ##table included in manuscript.
  Comb_coefficients <-rbind(female_analysis$my_coeff_res, male_analysis$my_coeff_res)

  write.table(Comb_coefficients,
              file=paste0("EXP_OUT_TABLES/",species_for_filename,"_fittedEXPONENTIAL_PARAMETERS_BYSEX_adultDOBthreshold.txt"),
              col.names = T,
              row.names = F,
              sep="\t",
              quote=F
              )

  ##output male and female plot (this uses patchwork)
  ##these are included as supplementary plots in manuscript
  Comb_plot <- female_analysis$my_plt/male_analysis$my_plt


   ggsave(paste0("EXP_OUT_PLOTS/",species_for_filename,"_fittedEXPONENTIAL_BYSEX_adultDOBthreshold.txt.png"),
          plot=Comb_plot,
          units="in",
          width =8.0,
          height=9.2,
          dpi=300)

  } #for each species


######done!######
```
